# Subunit selective modulation of GABA_A_ receptors using pharmacogenetically tethered neurosteroids

**DOI:** 10.64898/2026.03.23.713679

**Authors:** Ajeet Kumar, Hong-Jin Shu, Mariangela Chisari, Mingxing Qian, Yuanjian Xu, Pyeonghwa Jeong, Brenda C. Shields, Jiyong Hong, Michael R. Tadross, Douglas F. Covey, Charles F. Zorumski, Steven Mennerick

## Abstract

Neurosteroids are powerful endogenous modulators of inhibition and emerging therapeutics for anxiety, epilepsy, and mood disorders, yet their actions at defined receptor subtypes and within specific neuronal populations remain poorly resolved. Here, we engineer a neurosteroid DART (Drug Acutely Restricted by Tethering) platform to deliver neuroactive steroid (NAS) activity with cellular precision and receptor-subunit selectivity. From a screen of seventeen NAS analogs, we identified seven scaffolds suitable for further engineering, and we discovered that linker attachment at the steroid C11 position uniquely preserves NAS positive allosteric modulation of GABA_A_ receptors, whereas C2 and C17 attachment abolished activity. C11-linked NAS-DARTs slowed IPSC decay kinetics and showed variable off-target modulation of NMDA and AMPA EPSCs. The lead DART compound, YX85.1^DART.2^, enhanced GABA-evoked currents in neurons expressing engineered α4/δ-containing GABA_A_ receptors but spared γ2-containing receptors. A complementary benzodiazepine DART BZP.1^DART.2^ showed the opposite selectivity. Together, these tools enable cell-restricted, subunit-resolved interrogation of neurosteroid action on inhibitory microcircuits and provide a strategy to dissect how distinct GABA_A_ receptor subclasses contribute to circuit function and therapeutic outcomes.

## 1. Introduction

Gamma-aminobutyric acid type A (GABA_A_) receptors are the principal mediators of fast synaptic inhibition in the central nervous system (CNS). These receptors are heteropentamers derived from 19 subunit genes and are finely regulated by numerous endogenous and exogenous modulators of therapeutic interest, some of which have subunit selectivity (Belelli and Lambert, 2005; Chua and Chebib, 2017; Thompson, 2024; Zuo et al., 2026). Among the most potent of these are endogenous neurosteroids (NS) such as allopregnanolone and exogenous/synthetic neuroactive steroids (NAS) like alphaxalone, positive allosteric modulators (PAMs) of GABA_A_ receptors (Covey et al., 2023; Maguire and Mennerick, 2023). These actions contribute to anxiolytic, sedative, anticonvulsant, and antidepressant properties of NAS. Questions remain about the cell types and receptor composition contributing to both the therapeutic and unwanted side effects of NAS. A significant challenge in the field remains restricting the actions of NAS to specific cell types or brain circuits to study or invoke precise behavioral effects.

Advances in neuropharmacological targeting technologies have led to the development of Drug Acutely Restricted by Tethering (DART), a platform that enables precise delivery of pharmacological agents to genetically defined neuronal populations (Shields et al., 2017). The latest evolution of DART allows for bidirectional synaptic modulation with thousandfold cellular specificity, achieved by covalently tethering bioactive small molecules to HaloTag protein-bearing (HTP^+^) cell membranes. This approach has demonstrated functional specificity for modulating α-amino-3-hydroxy-5-methyl-4-isoxazolepropionic acid (AMPA) and GABA receptor activity (Shields et al., 2017, 2024), as well as G-protein coupled receptors (Shields et al., 2017; Sanchez et al., 2025).

Benzodiazepines (BZPs) are PAMs with selectivity for certain α- and γ-subunit-containing GABA_A_ receptors (Pritchett et al., 1989; Rudolph et al., 1999). By contrast, NAS have broad-spectrum effects on GABA_A_ receptors, with some preference for receptors containing a δ subunit rather than a γ subunit (Wohlfarth et al., 2002; Stell et al., 2003). While benzodiazepine.1^DART.2^ (BZP.1^DART.2^) was recently developed (Shields et al., 2024), a NAS DART would extend the GABA_A_ PAM toolset to the full repertoire of GABA_A_ receptor subtypes, enabling comprehensive, cell-type specific deconstruction of the behavioral impact of GABA PAM modulation (Sequeira et al., 2019; Maguire and Mennerick, 2023; Thompson, 2024).

In this study, we integrate principles from the medicinal chemistry of NAS with the second-generation DART.2 platform to evaluate candidate NAS-DART compounds and directly compare their actions with a benzodiazepine DART. Guided by structural features of the NAS scaffold, we explored multiple positions for DART linker attachment and identified carbon-11 on the steroid C-ring as a privileged site, as only C11-linked NAS-DARTs, unlike C2- or C17-linked analogs, retained detectable functional activity. These findings establish a rational structure-function framework for engineering tethered NAS compounds. By combining molecular pharmacology, structural NAS chemistry, and cell-specific chemical-genetic targeting, our study introduces a novel strategy to interrogate GABA_A_ receptor modulation with precision that distinguishes GABA_A_ receptor subclasses. Together, these advances establish NAS-DART as a new class of tethered modulators and, when used alongside a BZP.1^DART.2^, enable subunit-resolved dissection of NAS versus benzodiazepine-mediated GABA_A_ receptor signaling, laying the foundation for mechanistic dissection of their circuit and behavioral effects.

## 2. Materials and Methods

### 2.1 Primary Neuronal Culture

All *in vitro* experiments were performed using primary cultures of rat hippocampal neurons. Hippocampi were isolated from 1-3 day-old Sprague Dawley rat pups of either sex. All procedures were carried out in accordance with the National Institutes of Health guidelines for the care and use of laboratory animals and were approved by the Washington University Institutional Animal Care and Use Committee (IACUC). Dissection was performed in ice-cold calcium and magnesium-free HBSS, and tissue was digested with papain (20 U/mL; Worthington Biochem) for 20-30 minutes at 37°C, as described in previous publications (Mennerick et al., 1995). Following enzymatic digestion, hippocampi were gently triturated using fire-polished Pasteur pipettes until a homogeneous single-cell suspension was achieved. Cells were pelleted by low-speed centrifugation, resuspended in Neurobasal-A medium supplemented with B27, GlutaMAX, and penicillin-streptomycin, and plated onto 12-mm poly-D-lysine-coated coverslips. Cultures were maintained at 37°C with 5% CO_2_. Neurons were used for experiments between day *in vitro* (DIV) 10 and DIV12, a developmental window that exhibits strong functional inhibitory and excitatory synaptic connectivity.

### 2.2 Viral Constructs and Neuronal Transduction

To validate the DART targeting system in rat hippocampal neurons, we used AAV2 constructs expression sfHTP-GGSGG8-GPI with either mGreenLantern or dTomato (AAV2-CAG-sfHTP-GGSGG8-GPI-IRES-mGreenLantern or AAV2-CAG-sfHTP-GGSGG8-GPI-IRES-dTomato). These constructs express a surface HTP fused with a long flexible glycine-serine linker (GGSGG8) to a glycosylphosphatidylinositol (GPI) anchoring sequence to ensure extracellular accessibility for DART tethering. The fluorophores mGreenLantern or dTomato enabled visualization of HTP^+^ neurons during imaging and electrophysiology. To assess subunit-specific modulation of GABA**_A_** receptors by PAM-DART compounds, we expressed a picrotoxin-resistant mutant δ subunit (δ*) of GABA_A_ receptor using AAV8-SYN-rGABAδT6’Y-IRES-GFP (Shu et al., 2021). Because δ-subunit-containing GABA**_A_** receptors typically co-assemble with α4 (Barrera et al., 2008; Suryanarayanan et al., 2011), neurons transduced with δ* were co-transduced with WT α4 (AAV8-SYN-hGABAα4-IRES-GFP). The T6’Y mutation confers resistance to picrotoxin (Gurley et al., 1995; Sedelnikova et al., 2006), enabling functional isolation of δ-mediated currents in the presence of picrotoxin during electrophysiology. To measure PAM-DART effects in γ2-containing receptors, we expressed a picrotoxin-resistant γ2 mutant using AAV8-SYN-Glob-mGABAγ2-T6’Y-IRES-GFP (γ2*)(Shu et al., 2021). All viral vectors were applied to hippocampal neurons at DIV4-5. Neurons were returned to the incubator and allowed 5-7 days for expression and membrane localization of the receptor subunits and HaloTag constructs. All imaging and electrophysiology experiments were performed between DIV10 and DIV12.

### 2.3 NAS-DART synthesis and conjugation strategy

NAS-inspired scaffolds were synthesized using multi-step organic synthesis pathways optimized to preserve the integrity of the steroid nucleus. Functionalization of hydroxyl groups at the C2, C11, or C17 steroid positions was used to introduce a linker attachment point. This was intended to introduce a chemical handle without disrupting core pharmacophores known to influence GABA_A_ receptor binding. A flexible spacer was introduced, capable of allowing the tethered NAS sufficient conformational freedom to access its receptor binding pocket even when anchored at the cellular surface, and terminating in an alkyne to permit click-chemistry conjugation to azide-PEG_36_-HTL.2 (azide^DART.2^), generating a NAS-DART molecule capable of covalent tethering to HTP^+^ neurons. Synthetic details for the seventeen NAS scaffolds prior to attachment of azide^DART.2^, and a general procedure for the click chemistry reaction are reported in the **Supplemental Materials** (see also Shields et al., 2024). All final constructs (MQ302^DART.2^, MQ303^DART.2^, MQ311^DART.2^, MQ369^DART.2^, MQ392^DART.2^, YX62^DART.2^, and YX85.1^DART.2^) were characterized by ^1^H NMR and HRMS (**Supplementary Information**). Stocks were prepared in dimethylsulfoxide at 1–10 mM and diluted in physiological solution immediately before experiments.

### 2.4 DART ligand application in HTP^+^ neurons

To assess DART specificity, HTP^+^ neurons were incubated with 30 nM Alexa647.1^DART.2^ in artificial cerebrospinal fluid (ACSF) for 15-20 minutes at room temperature. This concentration provides optimal labeling while minimizing off-target membrane accumulation (Shields et al., 2017, 2024). After incubation, neurons were washed three times with ACSF to remove unbound ligand. Labeling was performed on live cells to ensure that surface HTP remained accessible and that dye accumulation reflected true DART tethering rather than internalization or nonspecific binding. For functional validation of tethered ligand activity, the known BZP.1^DART.2^ ligand (Shields et al., 2024) was applied for 20 min under identical conditions, followed by ACSF washes. Newly designed NAS-DART molecules (e.g., YX85.1^DART.2^, YX62^DART.2^, and MQ392^DART.2^) were applied at 500 nM concentration using the same incubation and washing protocol. After tethering, neurons were transferred to the recording chamber. Tethering efficiency was verified by fluorescence imaging before electrophysiological recordings. Non-transduced (HTP-negative) neurons in the same field served as internal controls for IPSC recordings. For reasons described below, we used HTP^+^ neurons treated with blank.1^DART.2^ as a control for AMPA excitatory postsynaptic currents (EPSCs) and in subunit selectivity experiments.

### 2.5 Fluorescence imaging

Fluorescence imaging was performed in a subset of experiments to verify specific DART binding in HTP^+^ neurons. Live hippocampal neurons were imaged by epifluorescence and phase contrast microscopy after incubation with Alexa647.1^DART.2^ to verify successful ligand tethering. Alexa-647 fluorescence was visualized using a far-red filter set, and mGreenLantern/GFP or dTomato reporters were detected with appropriate FITC or TRITC filter cubes. Imaging settings, including exposure time, gain, and illumination intensity, were kept constant across experimental groups to allow quantitative comparison of Alexa647.1^DART.2^ binding intensity between HTP^+^ neurons and HTP^−^ neurons.

### 2.6 Electrophysiology

Neurons were mounted on an Eclipse TE2000S inverted microscope. Whole-cell patch-clamp recordings were performed using a Multiclamp 700B amplifier (Molecular Devices), digitized through an Axon Digidata 1550 Low-Noise Acquisition System (Molecular Devices), and acquired and analyzed using pClamp 10 software (Molecular Devices). For recordings of GABAergic postsynaptic currents (IPSCs) and GABA-induced currents, the intracellular pipette solution contained (in mM): 130 CsCl, 4 NaCl, 10 HEPES, 5 EGTA, and 0.5 CaCl_2_, adjusted to pH 7.25 with NaOH. For glutamatergic postsynaptic currents (EPSCs) recordings, CsCl was replaced with CsMeSO_4_. Typically, the extracellular recording solution consisted of (in mM): 138 NaCl, 4 KCl, 10 HEPES, 10 glucose, 2 CaCl_2_, and 1 MgCl_2_, while in the case of *N*-Methyl-*D*-Aspartic acid (NMDA) EPSCs MgCl_2_ was not used in the bath solution, and 10 µM D-serine was added. Post-synaptic currents were isolated with appropriate antagonists. For IPSCs, 25 μM D-APV and 1 μM NBQX were used to block glutamate-mediated EPSCs. For AMPAR-mediated PSCs, 25 μM D-APV and 10 μM gabazine were used, and in the case of NMDA-mediated PSCs, D-APV was replaced with 1 μM NBQX. Patch pipettes were pulled from borosilicate glass capillaries (World Precision Instruments) and had resistances of 3–6 MΩ. Neurons were voltage-clamped at -70 mV unless otherwise indicated. Drug application was carried out via a gravity-fed local perfusion system with a common outlet placed ∼500 µm from the recorded cell.

### 2.7 NAS precursor screening and hit identification

NAS-like analogues were screened prior to DART linker addition for their ability to modulate GABA_A_ receptor-mediated currents using whole-cell voltage clamp and a gravity-fed, local perfusion system. To assess precursor activity, responses to baseline GABA (0.5 µM) were compared to responses to GABA co-applied with test molecules (0.3 µM). Peak amplitudes from co-applications were normalized to the neuron’s baseline GABA response. Compounds that produced enhancement of GABA-evoked peak current at p < 0.05 were classified as functional hits.

### 2.8 Analysis of IPSC and EPSC Kinetics

Data analysis was performed using pClamp/Clampfit 10 software. For spontaneous IPSCs and EPSCs, constant event detection thresholds were maintained across recordings to ensure comparability. IPSC kinetics were extracted from well-resolved events free of overlap, and decay time values were obtained using single- and double-exponential fits as appropriate. Event frequency and amplitude were calculated across stable 2–4-minute epochs.

### 2.9 Subunit-Selectivity Assay

Hippocampal neurons were transduced with an engineered picrotoxin-resistant δ subunit (δ*; as noted above) co-expressed with wild-type α4 and HTP, or with a picrotoxin-resistant γ2 subunit (γ2*) co-expressed with HTP. Following 20 min with YX85.1^DART.2^, BZP.1^DART.2^, or Blank.1^DART.2^ (control) incubation (0.5 µM) and subsequent washout, exogenous GABA-evoked currents were recorded only in δ*/γ2* and HTP^+^ neurons under constant picrotoxin (PTX) blockade (Sedelnikova et al., 2006; Shu et al., 2021) to block endogenous receptors.

### 2.10 Statistical Analysis

All statistical analyses were performed using GraphPad Prism. For comparisons between two independent groups, unpaired Student’s *t* tests were used. One-sample *t* tests were used to detect enhancement in precursor screening experiments. This initial precursor screen used a permissive threshold to capture weak modulators for further evaluation as DART constructs. (see below). Statistical significance was defined as *p* < 0.05 for all analyses. The ROUT method (robust regression and outlier removal) in GraphPad Prism was used to identify outliers. Data are reported in text and figures as mean ± SEM. Sample sizes (*N*) represent the number of neurons recorded, with each neuron considered an independent sample and samples were drawn from at least three independently prepared cultures.

## 3. Results

### 3.1. Identification of NAS-like molecules for DART linker conjugation

To identify potential NAS-like molecules suitable for tethering via the DART system, we screened 17 synthetic compounds without the DART linker on GABA_A_ responses in neurons. The goal of this experiment was to provide candidates for subsequent DART linker conjugation. Whole-cell patch-clamp recordings were performed in hippocampal neurons using 0.5 μM GABA to evoke baseline currents, followed by co-application of 0.3 μM of each candidate with GABA. Among the 17 compounds tested (**Figure 1A**), seven molecules were selected for DART conjugation. Six clearly potentiated GABA-induced currents, reflected by increases in peak amplitude of GABA currents (**Figure 1B, C**): MQ303, MQ311, MQ369, YX62, YX85, and MQ392. Additionally, MQ302 exhibited minimal PAM activity, but it was also chosen for DART conjugation to test whether a high tethered drug concentration would restore PAM activity.

**Figure 1.**
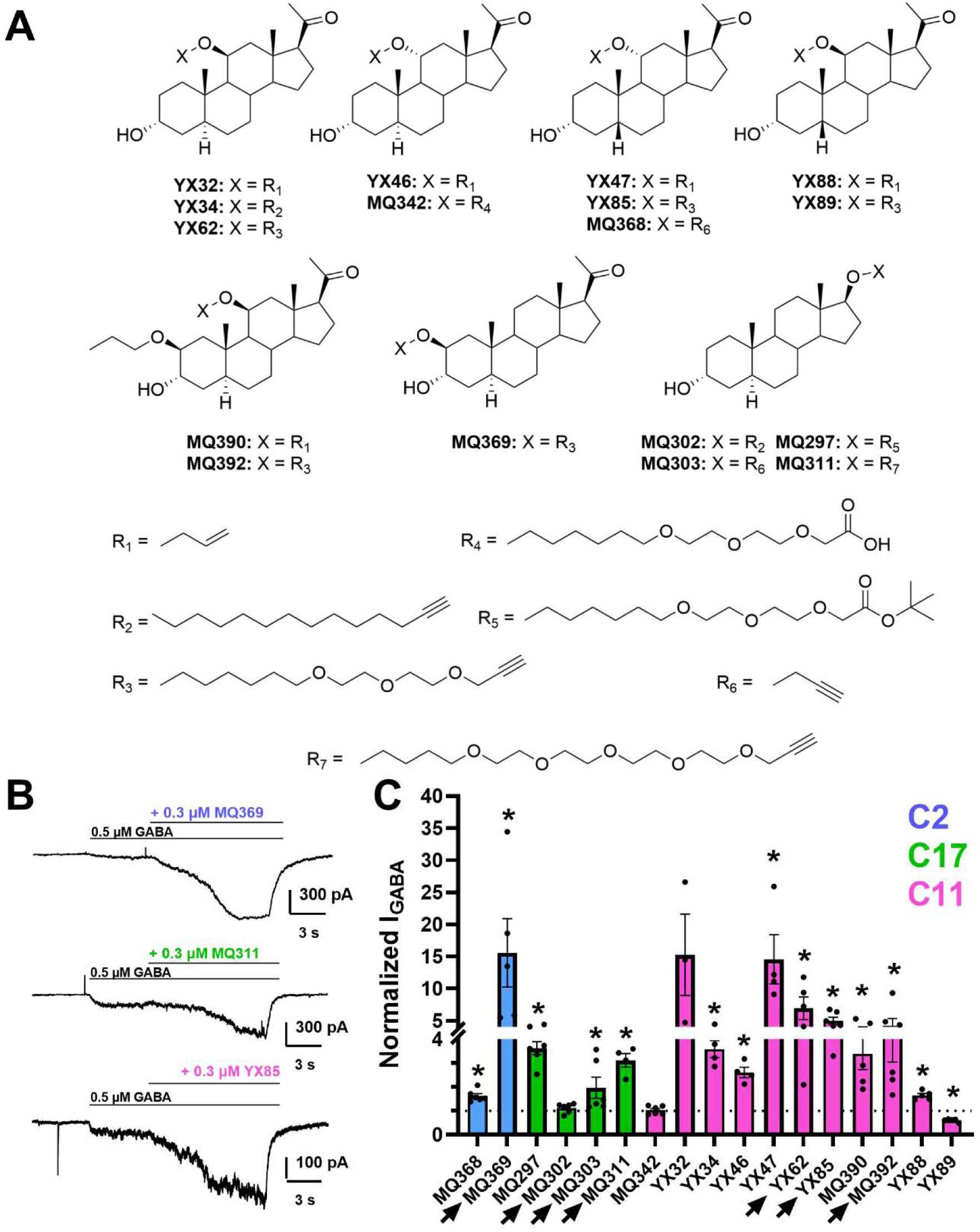
Chemical structures of synthesized NAS analogues and their functional screening for modulation of GABA_A_ receptor mediated currents. (A) NAS analogues with modifications for linker attachment at defined carbon positions. Compounds were either 5α reduced or 5β reduced, as both diastereomers result in active NAS (Covey et al., 2001). (B) Each analogue was tested for its ability to potentiate GABA-evoked currents using whole-cell patch-clamp recordings. Potentiation was quantified as the normalized peak current in response to that elicited by GABA alone (n = 5-6 cells per compound). * Indicate p <0.05 by one-sample t-test. Arrows next to compounds name indicate compounds advanced to DART linker/ligand conjugation.

### 3.2. Conjugation of NAS-like precursors to HaloTag-reactive linkers

To enable precise, cell-type-specific targeting of these prioritized molecules, we conjugated each selected NAS to the DART.2 linker system that contains a second-generation HTL.2 chloroalkane, which allows efficient tethering to HTP^+^ neurons, and a polyethylene glycol (PEG_36_) linker, which allows tethered drugs to span the distance between HTP and the native receptor of interest (**Figure 2A).** The attachment site varied among the seven molecules: C17 in three molecules, C2 in one molecule, and C11 in three molecules (**Figure 2B)**. These variations were designed to test whether the stereochemical position of attachment to the steroid backbone affects the ability of the tethered NAS to modulate GABA_A_ receptor function.

**Figure 2:**
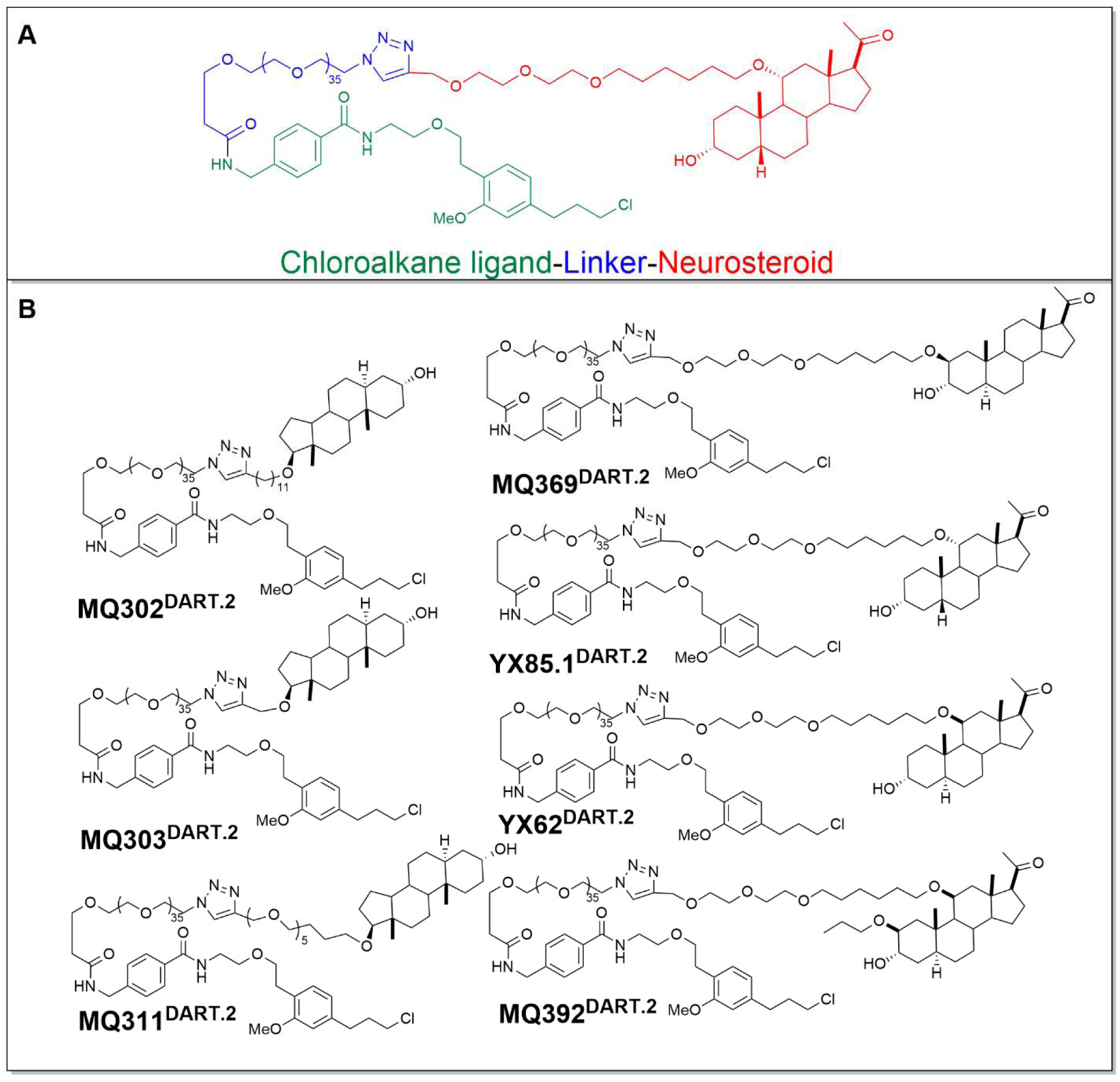
Conjugation of NAS-like molecules to HaloTag-reactive chloroalkane ligand. (A) Illustration of NAS-DART molecules, neurosteroid (red) attached with HaloTag reactive chloroalkane (green) by a large linker (blue) (B) Chemical structure of all tested NAS-DARTs showing conjugation of HaloTag-reactive linkers at different carbon positions.

### 3.3. Validation of cell-specific DART tethering in primary cultures of rat hippocampal neurons

To validate the fidelity of the DART platform in rat hippocampal neurons, we first confirmed cell-specific labeling following DART-ligand delivery. Selective delivery of DART conjugates to genetically defined neurons was achieved by transducing AAV2-CAG-sfHTP-GGSGG8-GPI-IRES-mGreenLantern to express surface HTP in cultured rat hippocampal neurons. Application of Alexa647.1^DART.2^ (30 nM), followed by wash, produced a strong membrane-associated fluorescence signal that precisely co-localized with mGreenLantern in HTP^+^ neurons, whereas no detectable Alexa-647 signal was observed in non-transduced neurons even under extended exposure conditions (**Figure 3**). This all-or-none labeling pattern (**Figure 3A**) demonstrated highly specific and background-free DART tethering to targeted neurons.

**Figure 3:**
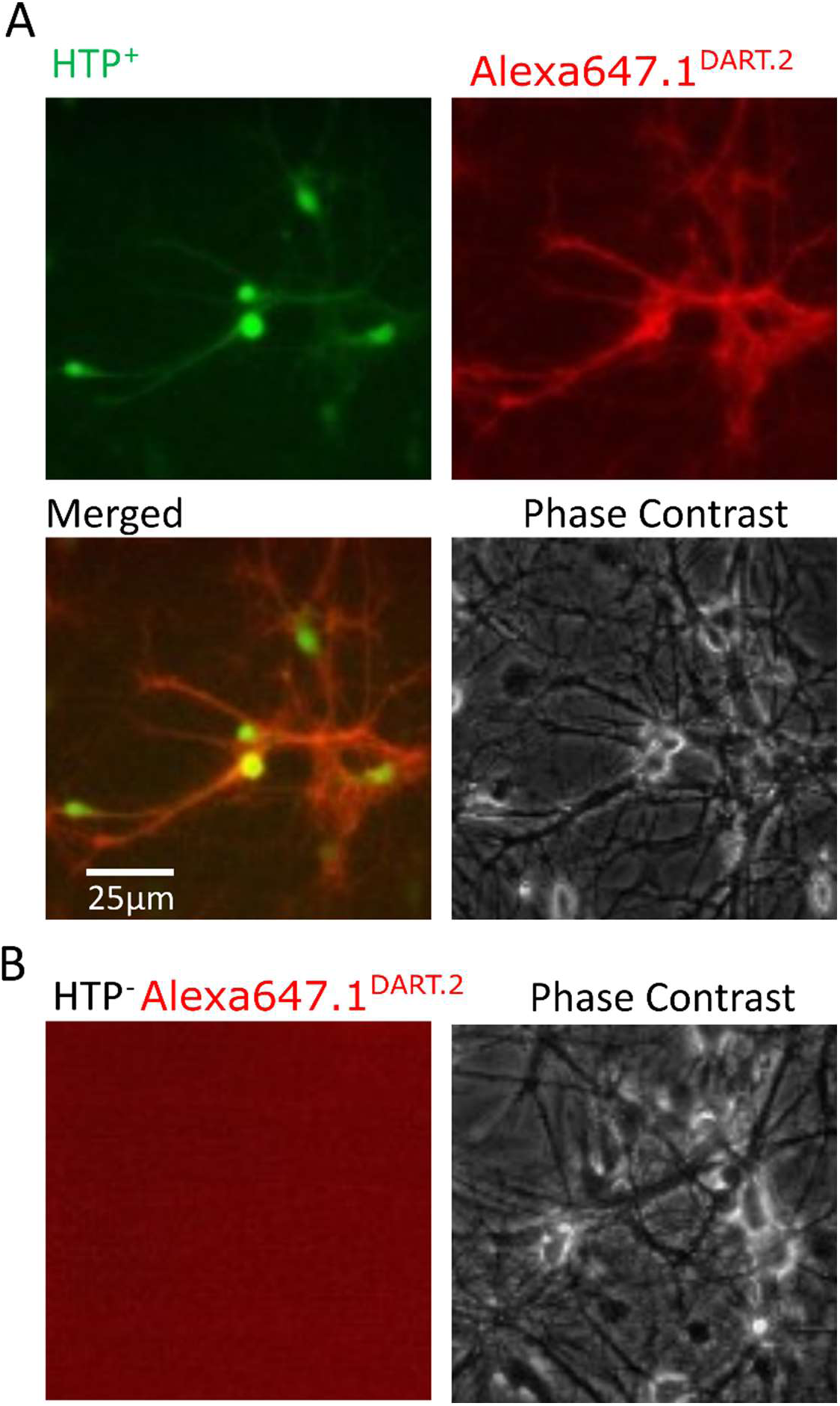
Validation of DART targeting primary rat hippocampal neuron cultures. (A) Representative fluorescence images showing selective labeling of HTP expressing (mGreen) hippocampal neurons by Alexa647.1^DART.2^ fluorescence dye (Red) and phase contrast image of same area. (B) Undetectable labeling with Alexa647.1^DART.2^ in non-transduced neurons at same exposure and display gain as in panel A.

To develop a screening workflow for GABA_A_R PAM NAS-DART ligands, we employed a previously validated BZP.1^DART.2^. We focused on spontaneous IPSC (sIPSC) decay as a readout because required incubation time and inherent variability of responses to exogenous GABA complicated within-cell assays, such as in Figure 1B. Specifically, sIPSC decay represents a relatively invariant parameter across cells and is robustly modulated by many classes of GABA_A_ receptor PAM (Otis and Mody, 1992; Chakrabarti et al., 2016). We incubated HTP-transduced neurons with 0.5 µM BZP.1^DART.2^ for 20 min, washed, and subsequently recorded spontaneous IPSCs in HTP^+^ and non-transduced neurons (HTP^−^) from the same culture dish. BZP.1^DART.2^ selectively prolonged the decay time constant of sIPSCs in HTP^+^ neurons with no effect on sIPSC amplitude or frequency (**Figure 4**). Importantly, HTP transduction alone did not produce any significant changes in sIPSC parameters when comparing HTP^+^ and HTP^−^ neurons (**Figure S1**), establishing a robust platform for screening. We evaluated off-target effects on excitatory transmission by evaluating NMDA **(Figure S2**), and AMPA EPSCs (**Figure S3**). Most parameters stayed intact, but we detected alteration of the decay time constant of AMPA EPSCs in HTP^+^ versus HTP^−^ neurons, which was traced to viral transduction rather than to an effect of HaloTag ligand (**Figure S3**). This observation was accounted for in the protocol for subsequent experiments examining EPSCs by using HTP+ neurons incubated in blank.1^DART.2^ as a control (Shields et al., 2024). Together, these results establish both the targeting fidelity and functionality of the DART platform in neuronal cultures, providing a validated foundation for subsequent exploration of NAS-DART compounds.

**Figure 4:**
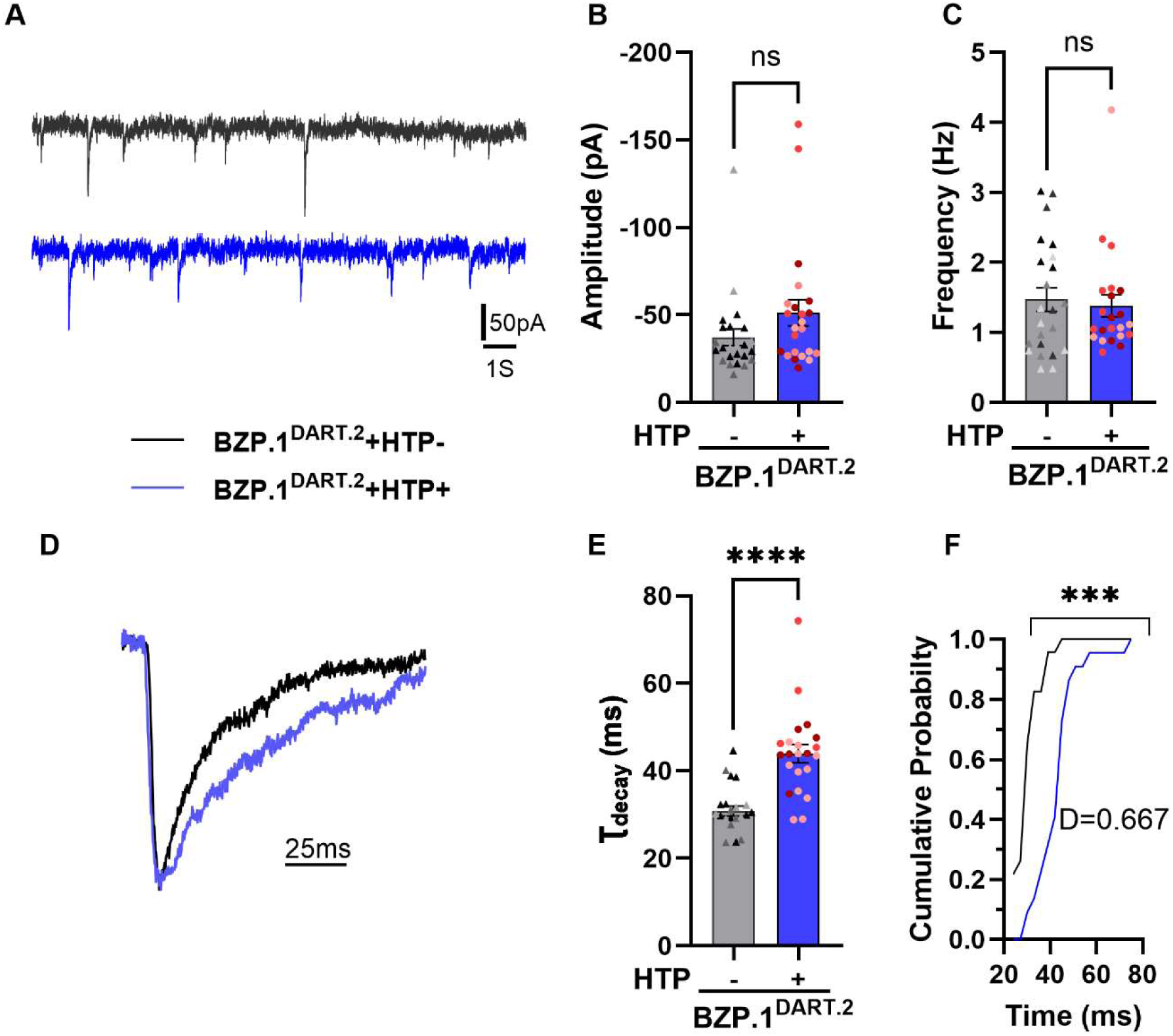
BZP.1^DART.2^ selectively modulates IPSCs in HTP neurons, validating the DART platform. (A) Representative raw traces of spontaneous IPSCs recorded from HTP^−^ (black trace) and HTP^+^ (blue trace) hippocampal neurons after application of BZP.1^DART.2^. (B) and (C) show summaries of amplitude and frequency respectively, derived from analysis of individual events as in panel A. Student’s *t* test showed p > 0.05 for both. Color shading of data points denote cells from independent culture. (D) Representative traces of peak-normalized, average IPSC waveform from HTP^−^ neurons (black trace) and HTP^+^ neurons (blue trace) treated with BZP.1^DART.2^, (E) Summary of BZP.1^DART.2^ on IPSCs event decay time constant (Student’s *t test,* *** p <0.001). (F) Re-plot of summary IPSC decay data as cumulative probability distributions (Kolmogorov-Smirnov test, **p <0.0007, Max difference D=0.6667). 1-2 cell(s) were excluded based on outlier criteria described in the methods.

### 3.4. Native cell subunits

Ultimately, we are interested in subunit selectivity, especially actions at δ-subunit containing receptors for NAS and γ-subunit containing receptors for benzodiazepines. Although γ contributions to IPSCs are expected, previous work from our group and others suggested a small but measurable contribution of δ-containing receptors to exogenous GABA responses in hippocampal cultures (Mangan et al., 2005; Shu et al., 2021). To confirm, we challenged sIPSCs with the selective δ PAM DS2 (1 µM) to query δ-receptor contributions to IPSCs. We found a small but measurable potentiation of sIPSC decay time constant (**Figure S4 A, B**), suggesting that δ-containing receptors contribute to IPSCs in culture neurons and provide a possible substrate for neurosteroids acting at δ-containing receptors. As expected, 0.5 µM diazepam (DZP), a γ-selective modulator, also prolonged hippocampal neurons IPSCs (**Figure S4 C, D**). These results suggest that hippocampal IPSCs are sensitive to both δ and γ selective modulators.

### 3.5. Functional profiling of NAS-DART to identify optimal conjugation site for preserving NAS activity

We next evaluated NAS-DART molecules using sIPSCs. Following 20 min incubation in 0.5 µM compound, as with BZP.1^DART.2^, C11 conjugated molecules YX62^DART.2^, YX85.1^DART.2^, and MQ369^DART.2^ prolonged the decay time constant of IPSCs in HTP^+^ neurons compared to HTP^−^neurons, with no detectable change in amplitude or frequency (**Figure 5**). Several molecules conjugated at C17 (e.g., MQ302 and MQ311) or C2 (MQ392) failed to measurably alter sIPSC decay kinetics when delivered through the DART system (**Figure S5-7**). This loss of activity indicates that the structural orientation and spatial presentation of the NAS relative to the receptor are critical, and that inappropriate linker positioning can prevent effective engagement of GABA_A_ receptor allosteric sites. These experiments revealed a striking position-dependent effect of DART conjugation on inhibitory synaptic function. This selective prolongation of IPSC kinetics, in the absence of changes in amplitude or frequency, is consistent with positive allosteric modulation of GABA_A_ receptor-mediated currents and demonstrates that C11 represents the optimal conjugation site for preserving NAS-mediated inhibitory potentiation.

**Figure 5:**
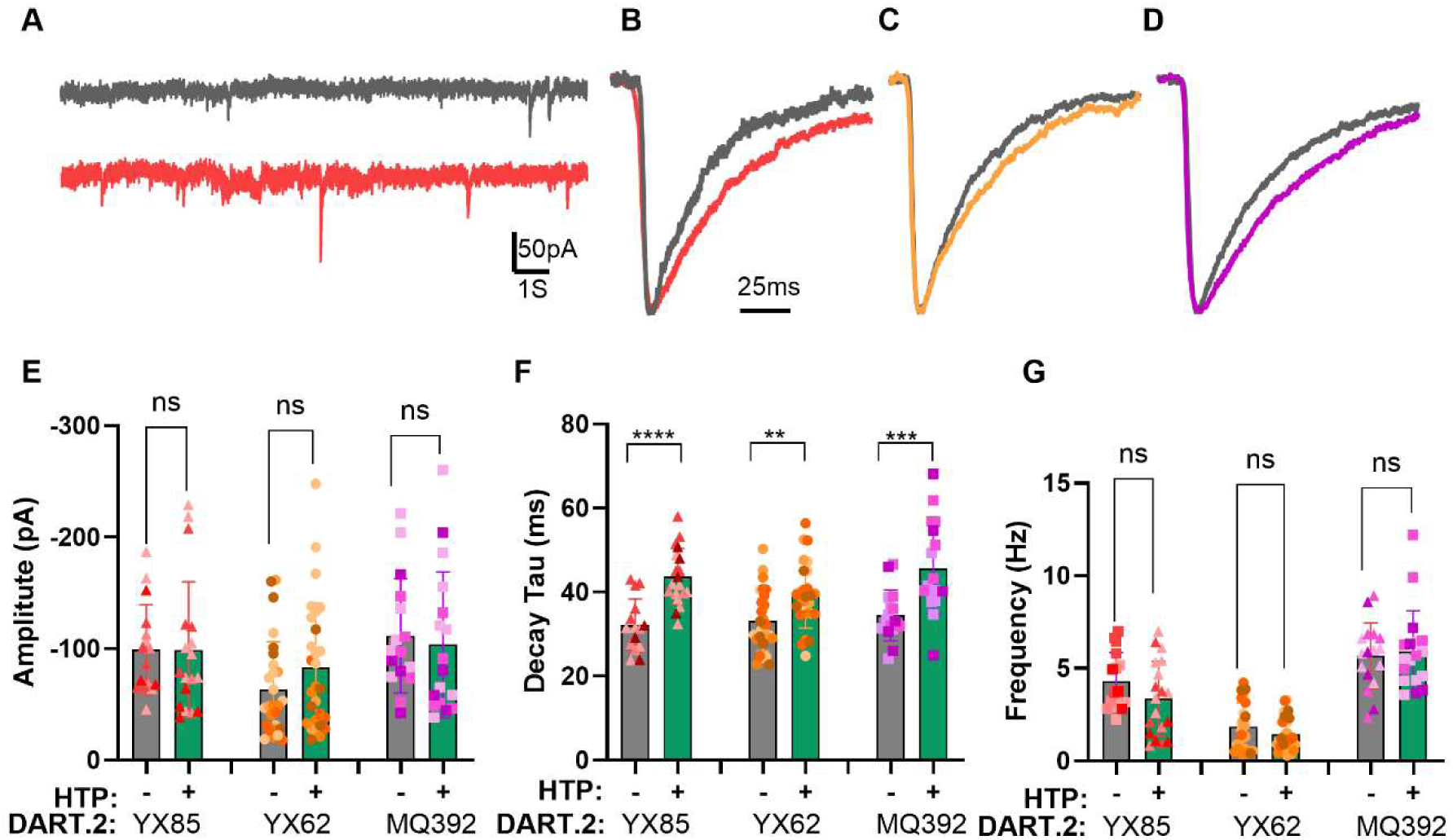
Functional profiling identifies optimal linker positions that preserve NAS activity. (A) Representative raw traces of spontaneous IPSCs recorded from HTP^−^ (black trace) and HTP^+^ (red trace) hippocampal neurons after application of YX85.1^DART.2^. (B-D) Representative traces of normalized average IPSCs events of HTP^−^ neurons (black trace) and HTP^+^ neurons (colored trace) treated with (B) YX85.1^DART.2^, (C) YX62^DART.2^ and (D) MQ392^DART.2^. (E-G) bar graphs of summary plots showing IPSCs amplitude, decay time constant and event frequency for each DART molecule in HTP^+^ neurons (green bars) with respective controls (HTP^−^ neurons, gray bar). Color shading of data points denote cells from independent platings. Statistical significance was assessed using Student’s *t* test (** p <0.01, *** p <0.001, **** p <0.0001), 1-2 cell(s) were excluded based on outlier criteria described in the methods.

Although NAS are best known for potentiating GABA_A_ receptors, several endogenous and synthetic NAS also modulate NMDA and AMPA receptors, either enhancing or inhibiting excitatory postsynaptic currents (EPSCs) (Mennerick et al., 2001; Borovska et al., 2012; Maguire and Mennerick, 2023). We evaluated EPSCs to help evaluate selectivity for GABA_A_ receptor effects. Therefore, all GABA-active NAS-DARTs were advanced to excitatory synapse assays. YX62^DART.2^, and YX85.1^DART.2^ did not show effects on NMDA receptor-mediated EPSCs, including amplitude, frequency, and decay time constant, in HTP^+^ neurons compared to HTP^−^ neurons treated in parallel (**Table 1**, **Figure S8)**. By contrast, MQ392^DART.2^ significantly increased NMDAR EPSC amplitude and frequency in HTP^+^ neurons compared to HTP^−^ neurons **Table 1**, **Figure S8**. Because AMPA EPSCs decay was intrinsically slower in HTP^+^ neurons (above and **Figure S3**), active NAS-DART effects were quantified relative to HTP^+^ neurons treated with blank.1^DART.2^ rather than HTP^−^ neurons (**Figure S9**). YX62^DART.2^ significantly increased AMPA EPSC amplitude and frequency without affecting the decay time constant (**Figure S9**), while MQ392^DART.2^ reduced the amplitude and prolonged AMPA EPSC decay without affecting frequency (**Figure S9**). In contrast, YX85.1^DART.2^ did not alter any features of AMPA EPSCs **Table 1**, **Figure S9**, demonstrating selective action at GABA_A_ receptors.

**Table 1:**
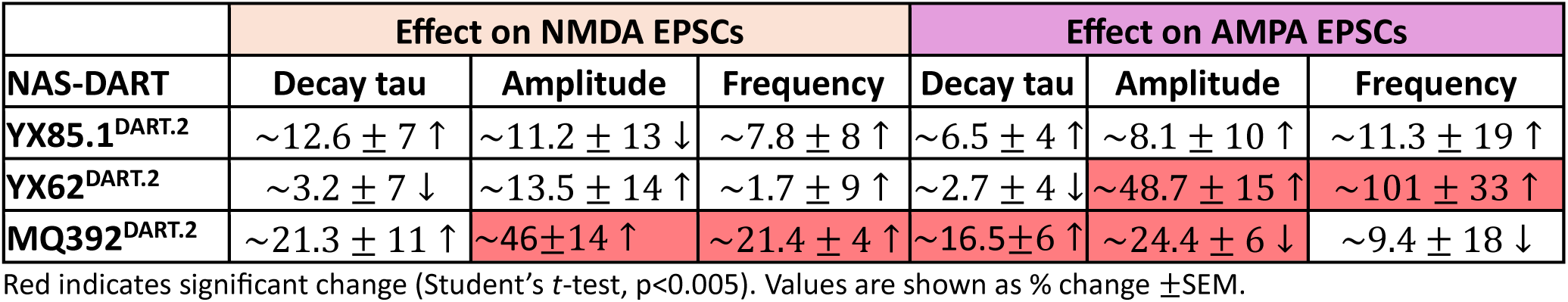
Activity of NAS-DARTs on EPSCs parameters.

Together, these findings show that excitatory modulation is not a universal property of NAS-DART molecules, but rather a scaffold-specific feature within the C11-linked active subgroup. To optimize selectivity for GABA_A_ receptor PAM actions, we focused subsequent studies on YX85.1^DART.2^, which exhibited robust effects on sIPSCs with no detectable effect on EPSCs.

### 3.6. Subunit Selectivity of active DART molecule and comparison with BZP.1^DART.2^

To compare the differential utility of BZP.1^DART.2^ with that of NAS-DART, we compared the subunit selectivity of both DART molecules. Whereas benzodiazepines require a γ subunit for effects (Günther et al., 1995), NAS may exhibit a preference for δ-containing receptors (Wohlfarth et al., 2002; Stell et al., 2003). **Figure S4** shows that IPSCs in cultured hippocampal neurons respond to both δ and γ modulators. To test subunit selectivity, we employed a subunit-engineering strategy that enabled pharmacological isolation of α4/δ or γ2-subunit-associated GABA responses within similar neuronal environments. Neurons were co-transduced with HTP and with either a picrotoxin (PTX)-resistant mutated (T6’Y) version of the δ subunit (δ*) (paired with preferred α4 subunit) or a mutated version of the γ2 subunit (γ2*). This allowed pharmacological isolation of the desired subunit-containing receptors. We measured exogenous GABA current density for these studies because of the tendency of δ-containing receptors to avoid synapses (Farrant and Nusser, 2005; Brickley and Mody, 2012). A concentration of GABA near the expected EC_50_ for each receptor class was adopted. In δ* receptors in presence of PTX (100µM), BZP.1^DART.2^ failed to affect GABA responses (0.5 μM GABA), but YX85.1^DART.2^ enhanced GABA current under the same conditions (**Figure 6**). In the complementary test, BZP.1^DART.2^ potentiated γ2* current density to GABA (10 µM) while YX85.1^DART.2^ failed to potentiate (**Figure 7**).

**Figure 6:**
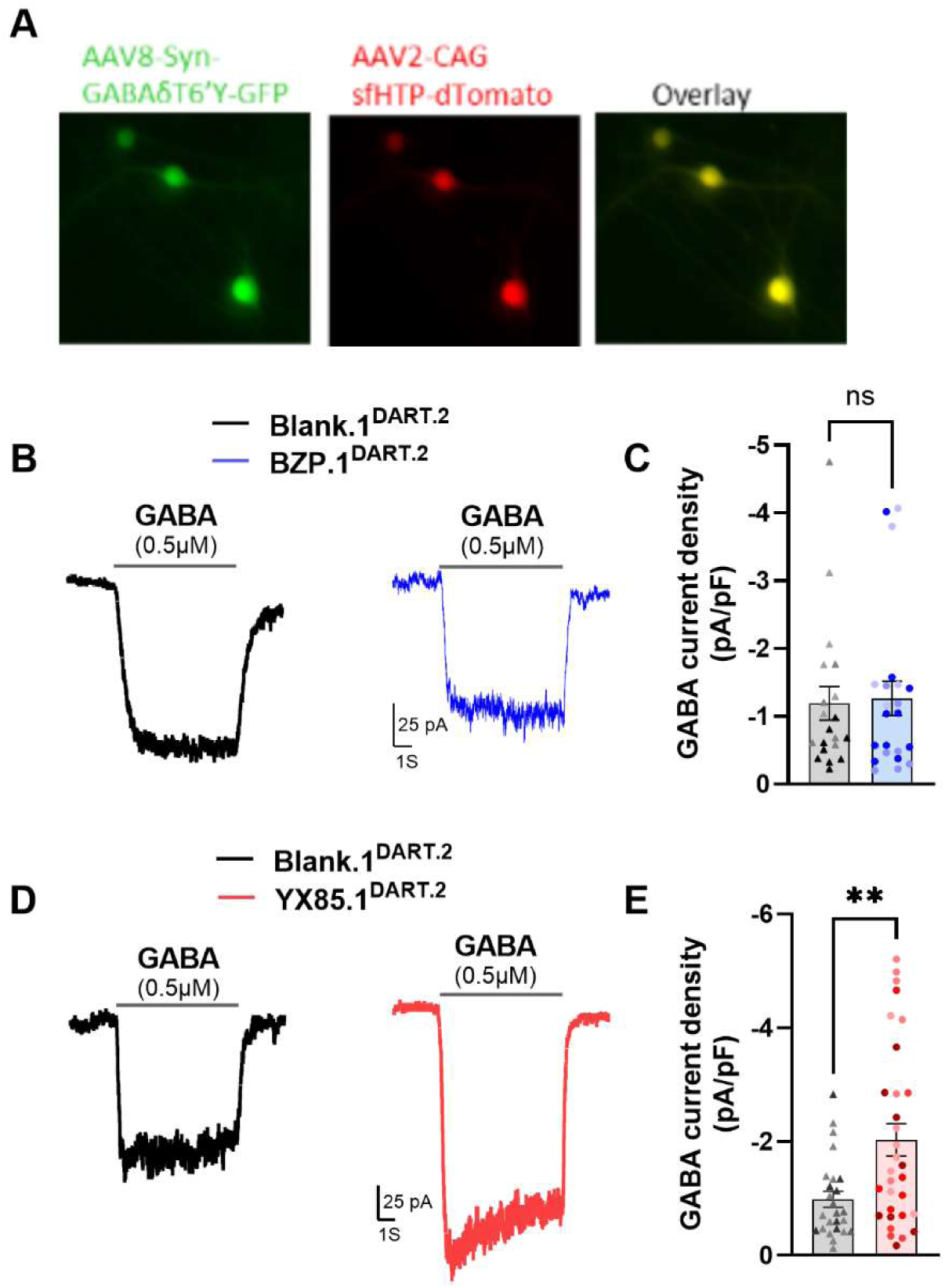
Comparison of the effects of NAS-DART and BZP.1^DART.2^ on δ subunit-containing GABA_A_ receptors. (A) δ* and HTP positive hippocampal neurons transduced with AAV8-Syn-GABAδT6’Y and AAV2-CAG-sfHTP-dTomato (B) Representative raw traces of GABA-evoked currents (0.5 µM, 5 s) recorded in the presence of 100 µM picrotoxin from α4, δ* (δT6′Y)-containing HTP^+^ neurons previously treated (20 min before recording) with either blank.1^DART.2^ (black trace) or BZP.1^DART.2^ (blue trace). (C) Quantification of GABA current density with color shading indicating cells from common plating (Student’s *t* test, p >0.05). (D) Representative raw traces of GABA-evoked currents (0.5 µM, 5 s) recorded in the presence of 100 µM picrotoxin from α4, δ (δT6′Y)-containing HTP^+^ neurons previously treated (20 min before recording) with either blank.1^DART.2^ (black trace) or BZP.1^DART.2^ (red trace). (E) Quantification of GABA current density (Student’s *t* test, **p <0.01). 1-2 cell(s) were excluded based on outlier criteria described in the methods.

**Figure 7:**
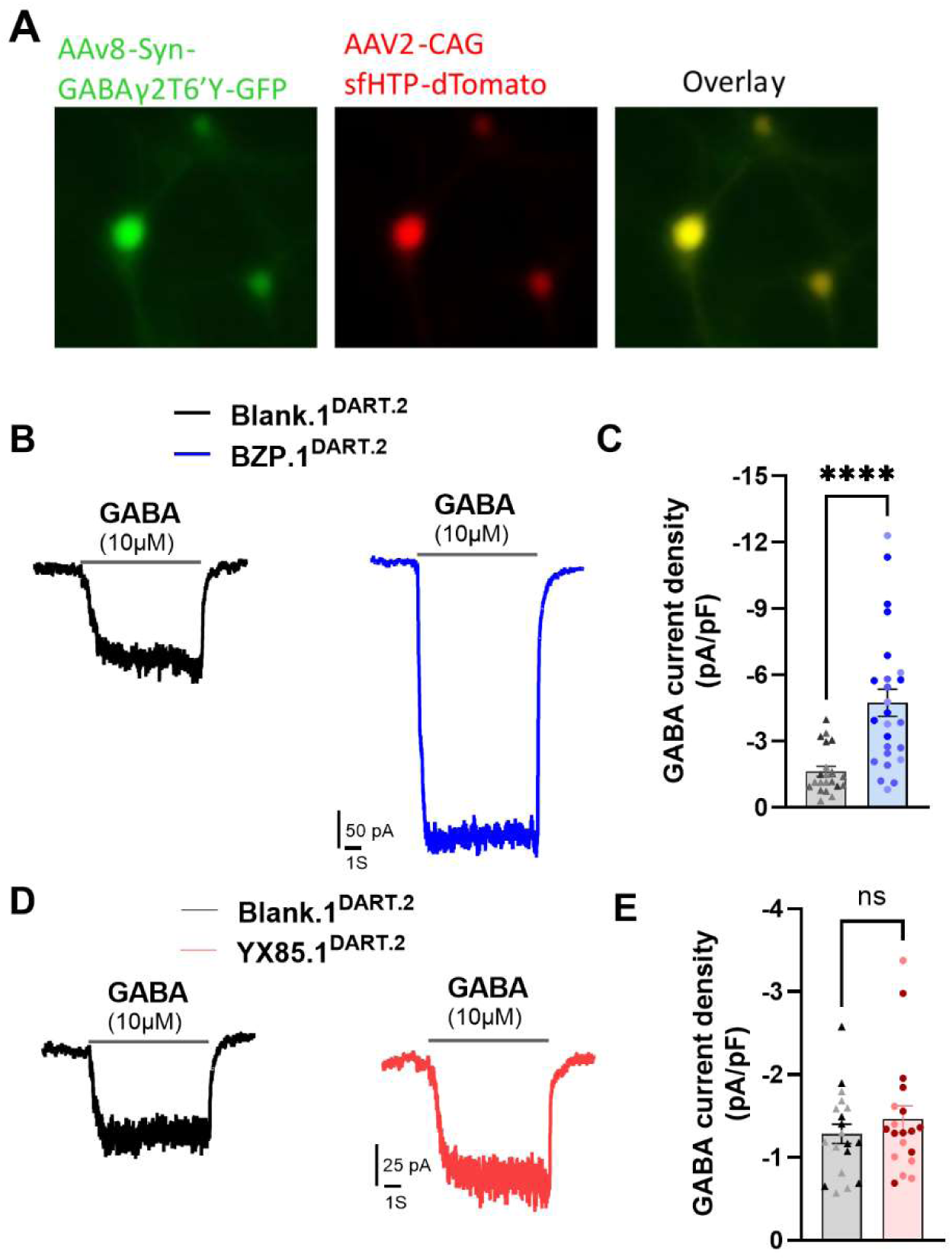
Comparison of the effects of NAS-DART and BZP.1^DART.2^ on γ2 subunit-containing receptors. (A) γ2* and HTP positive hippocampal neurons transduced with AAV8-Syn-GABAγ2T6’Y and AAV2-CAG-sfHTP-dTomato. (B) Representative raw traces of GABA-evoked currents (10 µM, 5 s) recorded in the presence of 100 µM picrotoxin from γ2* (γ2T6′Y) containing HTP^+^ neurons previously treated (20 min before recording) with either blank.1^DART.2^ (black trace) or BZP.1^DART.2^ (blue trace). (C) Quantification of GABA current density (Student’s *t* test, ****p <0.0001). (D) Representative raw traces of GABA-evoked currents (0.5 µM, 5 s) recorded in the presence of 100 µM picrotoxin from γ2* (γ2T6′Y)-containing HTP^+^ neurons previously treated (20 min before recording) with either blank.1^DART.2^ (black trace) or YX85.1^DART.2^ (Red trace). (E) Quantification of GABA current density (Student’s *t* test, p >0.05). 1-2 cell(s) were excluded based on outlier criteria described in the methods.

## 4. Discussion

NAS compounds are among the most potent and efficacious PAMs of GABA_A_ receptors, yet their widespread access to multiple neuronal populations, diverse receptor subtypes, and intracellular targets has obscured the mechanisms underlying their therapeutic and behavioral actions (Belelli and Lambert, 2005; Balan et al., 2019, 2023; Ishikawa et al., 2021, 2022; Kumar et al., 2025). Furthermore, a growing list of intracellular targets for NAS has been identified, (Balan et al., 2019, 2023; Ishikawa et al., 2022; Kumar et al., 2025) reflecting their ability to readily penetrate cell membranes and accumulate within cells (Jiang et al., 2016; Zorumski et al., 2025). Currently, there is no good way to limit NAS actions to membrane receptors. In this study, we introduce a NAS-DART platform that overcomes the above limitations by enabling cell-type-specific and receptor-subtype-resolved delivery of NAS activity, limited to the membrane surface. By integrating HaloTag-based genetic targeting (Shields et al., 2017, 2024), structure-guided chemical conjugation, and electrophysiological interrogation across δ- and γ2-containing GABA_A_ receptors, we preserve core NAS pharmacology, identify a privileged conjugation site, and demonstrate δ-selective modulation with a level of precision difficult to achieve through conventional bath application of free ligand to native cells.

A fundamental requirement for any targeted pharmacology approach is that the system itself must not perturb baseline neuronal physiology. We did find that HaloTag receptor expression itself altered AMPAR EPSC kinetics (**Figure S3**). Although the basis of this effect remains unknown, we accounted for this potential confound by comparing the effect of blank.1^DART.2^ vs. NAS-DART on AMPAR EPSCs in HTP^+^ neurons (**Figure S9**). Further, some NAS-DART compounds exhibited off-target effects on excitatory transmission (**Figure S8-9**). However, the high specificity of YX85-DART mirrors the single-cell precision originally demonstrated in the DART framework (Shields et al., 2017, 2024; Sanchez et al., 2025). This confirms that NAS-DART can be used for selective, cell-restricted pharmacology. Results with YX85.1^DART.2^ validate NAS-DART as a stable and selective platform for dissecting inhibitory mechanisms in defined neuronal populations.

Our structure-activity analysis revealed that only C11-linked NAS-DART constructs preserved functional potency, whereas C2 and C17 modifications abrogated activity. This position-specific permissiveness is consistent with extensive structure-activity and structural analyses demonstrating that NAS potentiation depends critically on the C3 α-hydroxyl and C20 carbonyl groups, while the C11 position on the C ring is structurally tolerant of substitution (Akk et al., 2007; Laverty et al., 2017; Mortensen et al., 2025). Identification of C11 as a privileged attachment site therefore provides a clear design principle for engineering next-generation NAS probes, including tethered ligands, photopharmacological tools, and imaging-compatible analogs that preserve functional activity while enabling spatial control. Within the C11-linked class, excitatory modulation proved to be scaffold-dependent: only selected compounds influenced NMDA or AMPA EPSCs. This pattern could be consistent with the intrinsic diversity of NAS actions, in which some analogs, such as pregnenolone sulfate, modulate excitatory transmission, whereas others, including allopregnanolone-like compounds, remain predominantly GABAergic (Korinek et al., 2011; Borovska et al., 2012; Ziolkowski et al., 2020). However, we cannot exclude the possibility of off-target effects that would not be mimicked by free NAS.

The most striking finding was that YX85.1^DART.2^ preferentially potentiated δ-containing GABA_A_ receptors. Using HaloTag targeting together with picrotoxin-resistant T6’Y subunits to pharmacologically isolate α4/δ versus γ2 receptors, YX85.1^DART.2^ robustly enhanced GABA-evoked current density in α4/δ-expressing neurons but produced no measurable effect on γ2-containing receptors. This δ-selectivity is consistent with the known high sensitivity of δ receptors to neurosteroids and demonstrates that the DART approach can achieve receptor-subtype-resolved modulation. (Wohlfarth et al., 2002; Stell et al., 2003; Belelli and Lambert, 2005; Brickley and Mody, 2012). Although tethering places ligand near membrane receptors, the observed selectivity suggests the effective local concentration mimics that of low free NAS, because higher free-ligand concentrations typically reduce subunit selectivity (Wohlfarth et al., 2002). The apparent discrepancy between YX85.1^DART.2^’s prolongation of IPSC decay and its lack of γ2 potentiation may reflect several possibilities: contributions of δ receptors to some IPSCs (Ye et al., 2013; Jager et al., 2016; Sun et al., 2018, 2020), differing α/β subunit contexts, unanticipated effects of the T6’Y mutation, or restricted DART access at synaptic sites. The effect of the δ-selective PAM DS2 (**Figure S4**) suggests that δ receptor contribution to IPSCs in hippocampal cultures explains at least part of the discrepancy.

Importantly, BZP.1^DART.2^ displayed effects consistent with classical benzodiazepine-mediated positive allosteric modulation, selectively enhancing agonist responses at γ2-containing receptors (Buhr et al., 1996; Campo-Soria et al., 2006; Zhu et al., 2018). The complementary selectivity profiles of YX85.1^DART.2^ and BZP.1^DART.2^ therefore establish two orthogonal, non-overlapping pharmacological profiles. Thus, DART conjugation delivers modulatory activity with both spatial precision and receptor-subunit fidelity. Because subunit expression can be cell-type dependent, DART promises to interrogate the role of inhibition in specific neuronal populations in behavioral outcomes such as anxiolysis, antidepressant activity, seizure protection, or sensory gating, without off-target effects that may limit clinical utility (Gunn et al., 2004; Belelli and Lambert, 2005; Ziolkowski et al., 2020; Maguire and Mennerick, 2023). By integrating molecular pharmacology, structure-guided NAS chemistry, and advanced targeting technology, this work establishes a platform for precision NAS-mediated modulation of neuronal circuits. Looking forward, an important next step will be to test the behavioral relevance of cell-restricted GABA_A_ receptor modulation in vivo.

In summary, this study introduces a NAS-DART platform that preserves essential NAS pharmacology while enabling cell-type-specific and receptor-subunit-specific control of inhibitory signaling. By uncovering C11 as a privileged conjugation site, revealing scaffold-dependent inhibitory modulation, and demonstrating δ-selective potentiation by YX85.1^DART.2^ alongside complementary γ2-selective BZP.1^DART.2^ effects, we establish a foundation for the mechanistic dissection of inhibitory microcircuits and the development of next-generation NAS therapeutics with tailored spatial and pharmacodynamic profiles.

## Acknowledgements

The work was supported by MH122379, MH123748 and the Taylor Family Institute for Innovative Psychiatric Research, and by RF1-MH117055 and R01-MH135932 (to M.R.T.). The authors thank members of the Taylor Family Institute, Steve Traynelis, Lisa Monteggia, and Bernhard Lüscher for discussion.

---

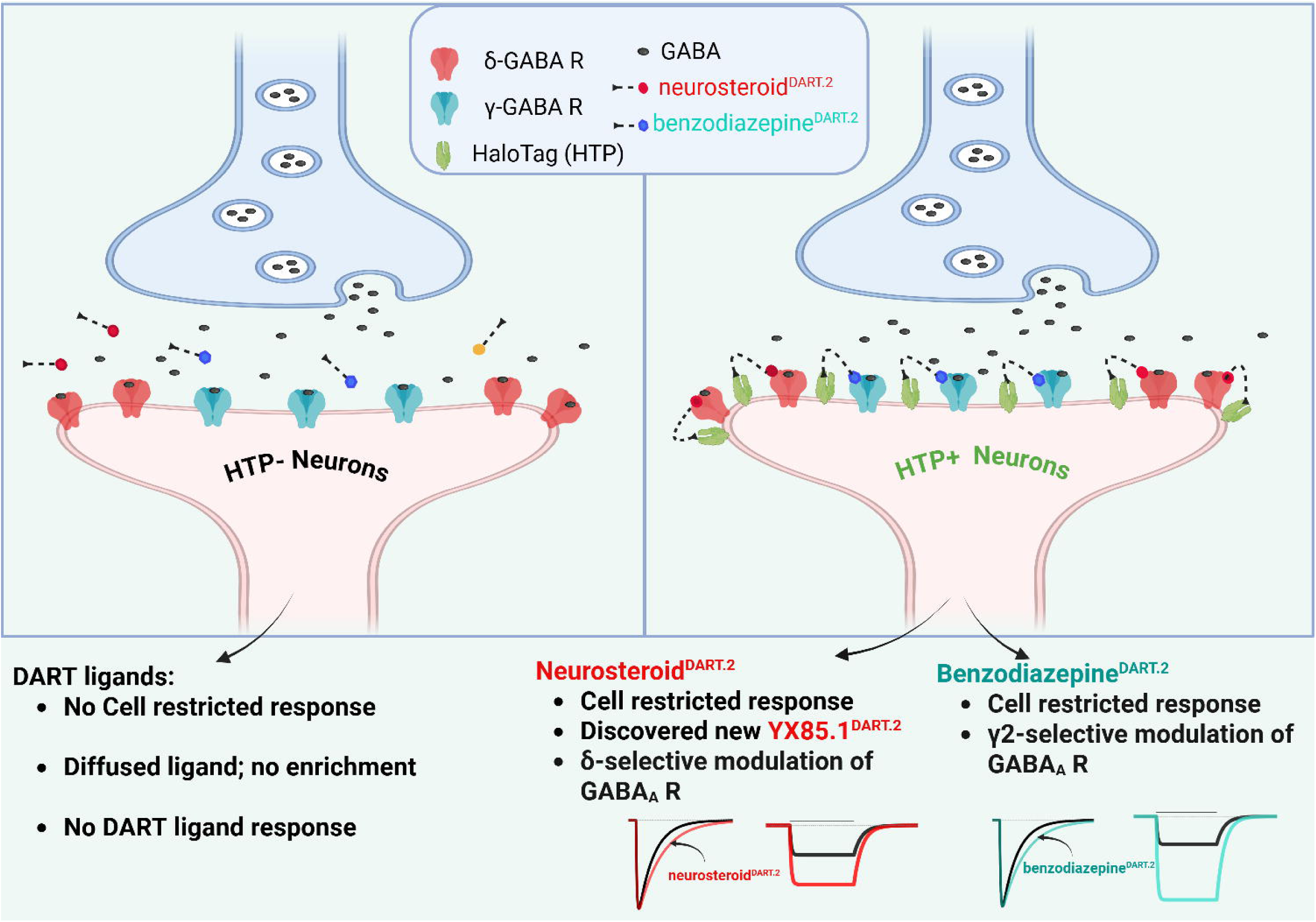

## References

Akk, G., Covey, D.F., Evers, A.S., Steinbach, J.H., Zorumski, C.F., and Mennerick, S. (2007). Mechanisms of Neurosteroid Interactions with GABAA Receptors. Pharmacology & Therapeutics 116: 35–57.

Balan, I., Beattie, M.C., O’Buckley, T.K., Aurelian, L., and Morrow, A.L. (2019). Endogenous neurosteroid (3α,5α)3-hydroxypregnan-20-one inhibits toll-like-4 receptor activation and pro-inflammatory signaling in macrophages and brain. Sci. Rep. 9: 1220.

Balan, I., Patterson, R., Boero, G., Krohn, H., O’Buckley, T.K., Meltzer-Brody, S., et al. (2023). Brexanolone therapeutics in post-partum depression involves inhibition of systemic inflammatory pathways. EBioMedicine 89: 104473.

Barrera, N.P., Betts, J., You, H., Henderson, R.M., Martin, I.L., Dunn, S.M.J., et al. (2008). Atomic force microscopy reveals the stoichiometry and subunit arrangement of the alpha4beta3delta GABA(A) receptor. Mol. Pharmacol. 73: 960–967.

Belelli, D., and Lambert, J.J. (2005). Neurosteroids: endogenous regulators of the GABA(A) receptor. Nat. Rev. Neurosci. 6: 565–575.

Borovska, J., Vyklicky, V., Stastna, E., Kapras, V., Slavikova, B., Horak, M., et al. (2012). Access of inhibitory neurosteroids to the NMDA receptor. Br. J. Pharmacol. 166: 1069–1083.

Brickley, S.G., and Mody, I. (2012). Extrasynaptic GABA(A) receptors: their function in the CNS and implications for disease. Neuron 73: 23–34.

Buhr, A., Baur, R., Malherbe, P., and Sigel, E. (1996). Point mutations of the alpha 1 beta 2 gamma 2 gamma-aminobutyric acid(A) receptor affecting modulation of the channel by ligands of the benzodiazepine binding site. Mol. Pharmacol. 49: 1080–1084.

Campo-Soria, C., Chang, Y., and Weiss, D.S. (2006). Mechanism of action of benzodiazepines on GABAA receptors. Br. J. Pharmacol. 148: 984–990.

Chakrabarti, S., Qian, M., Krishnan, K., Covey, D.F., Mennerick, S., and Akk, G. (2016). Comparison of steroid modulation of spontaneous inhibitory postsynaptic currents in cultured hippocampal neurons and steady-state single-channel currents from heterologously expressed α1β2γ2L GABA(A) receptors. Mol. Pharmacol. 89: 399–406.

Chua, H.C., and Chebib, M. (2017). Chapter One - GABAA Receptors and the Diversity in their Structure and Pharmacology. In Advances in Pharmacology, D.P. Geraghty, and L.D. Rash, eds. (Academic Press), pp 1–34.

Covey, D.F., Evers, A.S., Izumi, Y., Maguire, J.L., Mennerick, S.J., and Zorumski, C.F. (2023). Neurosteroid enantiomers as potentially novel neurotherapeutics. Neurosci. Biobehav. Rev. 149: 105191.

Covey, D.F., Evers, A.S., Mennerick, S., Zorumski, C.F., and Purdy, R.H. (2001). Recent developments in structure-activity relationships for steroid modulators of GABA(A) receptors. Brain Res. Brain Res. Rev. 37: 91–97.

Farrant, M., and Nusser, Z. (2005). Variations on an Inhibitory Theme: Phasic and Tonic Activation of GABAA Receptors. Nature Reviews. Neuroscience 6: 215–229.

Gunn, B.G., Cunningham, L., Mitchell, S.G., Swinny, J.D., Lambert, J.J., Belelli, D., et al. (2004). Neuroactive Steroid Pregnenolone Sulphate Inhibits Long-Term Potentiation via Activation of Alpha2-Adrenoreceptors at Excitatory Synapses in Rat Medial Prefrontal Cortex. The International Journal of Neuropsychopharmacology 36: 611–624.

Günther, U., Benson, J., Benke, D., Fritschy, J.M., Reyes, G., Knoflach, F., et al. (1995). Benzodiazepine-insensitive mice generated by targeted disruption of the gamma 2 subunit gene of gamma-aminobutyric acid type A receptors. Proc. Natl. Acad. Sci. U. S. A. 92: 7749–7753.

Gurley, D., Amin, J., Ross, P.C., Weiss, D.S., and White, G. (1995). Point mutations in the M2 region of the alpha, beta, or gamma subunit of the GABAA channel that abolish block by picrotoxin. Receptors Channels 3: 13–20.

Ishikawa, M., Nakazawa, T., Kunikata, H., Sato, K., Yoshitomi, T., Krishnan, K., et al. (2022). The enantiomer of allopregnanolone prevents pressure-mediated retinal degeneration via autophagy. Front. Pharmacol. 13: 855779.

Ishikawa, M., Takaseki, S., Yoshitomi, T., Covey, D.F., Zorumski, C.F., and Izumi, Y. (2021). The neurosteroid allopregnanolone protects retinal neurons by effects on autophagy and GABRs/GABAA receptors in rat glaucoma models. Autophagy 17: 743–760.

Jager, P., Ye, Z., Yu, X., Zagoraiou, L., Prekop, H.-T., Partanen, J., et al. (2016). Tectal-derived interneurons contribute to phasic and tonic inhibition in the visual thalamus. Nat. Commun. 7: 13579.

Jiang, X., Shu, H.-J., Krishnan, K., Qian, M., Taylor, A.A., Covey, D.F., et al. (2016). A clickable neurosteroid photolabel reveals selective Golgi compartmentalization with preferential impact on proximal inhibition. Neuropharmacology 108: 193–206.

Korinek, M., Kapras, V., Vyklicky, V., Adamusova, E., Borovska, J., Vales, K., et al. (2011). Neurosteroid modulation of N-methyl-D-aspartate receptors: molecular mechanism and behavioral effects. Steroids 76: 1409–1418.

Kumar, A., Qian, M., Xu, Y., Benz, A., Covey, D.F., Zorumski, C.F., et al. (2025). Unravelling the multifaceted actions of neurosteroids: Machine learning and in vitro screening for novel target discovery. Br. J. Pharmacol.

Laverty, D., Thomas, P., Field, M., Andersen, O.J., Gold, M.G., Biggin, P.C., et al. (2017). Crystal structures of a GABAA-receptor chimera reveal new endogenous neurosteroid-binding sites. Nat. Struct. Mol. Biol. 24: 977–985.

Maguire, J.L., and Mennerick, S. (2023). Neurosteroids: mechanistic considerations and clinical prospects. Neuropsychopharmacology.

Mangan, P.S., Sun, C., Carpenter, M., Goodkin, H.P., Sieghart, W., and Kapur, J. (2005). Cultured hippocampal pyramidal neurons express two kinds of GABA_A_ receptors. Mol. Pharmacol. 67: 775–788.

Mennerick, S., Que, J., Benz, A., and Zorumski, C.F. (1995). Passive and synaptic properties of hippocampal neurons grown in microcultures and in mass cultures. J. Neurophysiol. 73: 320–332.

Mennerick, S., Zeng, C.M., Benz, A., Shen, W., Izumi, Y., Evers, A.S., et al. (2001). Effects on gamma-aminobutyric acid (GABA)(A) receptors of a neuroactive steroid that negatively modulates glutamate neurotransmission and augments GABA neurotransmission. Mol. Pharmacol. 60: 732–741.

Mortensen, M., Bright, D.P., Fagotti, J., Dorovykh, V., Cerna, B., and Smart, T.G. (2025). Forty years searching for neurosteroid binding sites on GABAA receptors. Neuroscience 578: 6–24.

Otis, T.S., and Mody, I. (1992). Modulation of decay kinetics and frequency of GABAA receptor-mediated spontaneous inhibitory postsynaptic currents in hippocampal neurons. Neuroscience 49: 13–32.

Pritchett, D.B., Sontheimer, H., Shivers, B.D., Ymer, S., Kettenmann, H., Schofield, P.R., et al. (1989). Importance of a novel GABA_A_ receptor subunit for benzodiazepine pharmacology. Nature 338: 582–585.

Rudolph, U., Crestani, F., Benke, D., Brunig, I., Benson, J.A., Fritschy, J.M., et al. (1999). Benzodiazepine actions mediated by specific g-aminobutyric acidA receptor subtypes. Nature 401: 796–800.

Sanchez, J., Bonifazi, A., Groom, S., Sambrook, M.O., Camacho-Hernandez, G.A., Kuijer, E.J., et al. (2025). Targeted inhibition of mu-opioid receptors in neuronal subpopulations by membrane-tethered Naloxo-DART antagonists. Cell Chem. Biol.

Sedelnikova, A., Erkkila, B.E., Harris, H., Zakharkin, S.O., and Weiss, D.S. (2006). Stoichiometry of a pore mutation that abolishes picrotoxin-mediated antagonism of the GABAA receptor. J. Physiol. 577: 569–577.

Sequeira, A., Shen, K., Gottlieb, A., and Limon, A. (2019). Human brain transcriptome analysis finds region- and subject-specific expression signatures of GABAAR subunits. Commun. Biol. 2: 153.

Shields, B.C., Kahuno, E., Kim, C., Apostolides, P.F., Brown, J., Lindo, S., et al. (2017). Deconstructing behavioral neuropharmacology with cellular specificity. Science 356: eaaj2161.

Shields, B.C., Yan, H., Lim, S.S.X., Burwell, S.C.V., Cammarata, C.M., Fleming, E.A., et al. (2024). DART.2: bidirectional synaptic pharmacology with thousandfold cellular specificity. Nat. Methods 21: 1288–1297.

Shu, H.-J., Lu, X., Bracamontes, J., Steinbach, J.H., Zorumski, C.F., and Mennerick, S. (2021). Pharmacological and biophysical characteristics of picrotoxin-resistant, δSubunit-containing GABAA receptors. Front. Synaptic Neurosci. 13: 763411.

Stell, B.M., Brickley, S.G., Tang, C.Y., Farrant, M., and Mody, I. (2003). Neuroactive steroids reduce neuronal excitability by selectively enhancing tonic inhibition mediated by delta subunit-containing GABAA receptors. Proc. Natl. Acad. Sci. U. S. A. 100: 14439–14444.

Sun, M.-Y., Shu, H.-J., Benz, A., Bracamontes, J., Akk, G., Zorumski, C.F., et al. (2018). Chemogenetic isolation reveals synaptic contribution of δ GABAA receptors in mouse dentate granule neurons. J. Neurosci. 38: 8128–8145.

Sun, M.-Y., Ziolkowski, L., and Mennerick, S. (2020). δ subunit-containing GABAA IPSCs are driven by both synaptic and diffusional GABA in mouse dentate granule neurons. J. Physiol. 598: 1205–1221.

Suryanarayanan, A., Liang, J., Meyer, E.M., Lindemeyer, A.K., Chandra, D., Homanics, G.E., et al. (2011). Subunit compensation and plasticity of synaptic GABA(A) receptors induced by ethanol in α4 subunit knockout mice. Front. Neurosci. 5: 110.

Thompson, S.M. (2024). Modulators of GABAA receptor-mediated inhibition in the treatment of neuropsychiatric disorders: past, present, and future. Neuropsychopharmacology 49: 83–95.

Wohlfarth, K.M., Bianchi, M.T., and Macdonald, R.L. (2002). Enhanced neurosteroid potentiation of ternary GABA(A) receptors containing the delta subunit. J. Neurosci. 22: 1541–1549.

Ye, Z., McGee, T.P., Houston, C.M., and Brickley, S.G. (2013). The contribution of δ subunit-containing GABAA receptors to phasic and tonic conductance changes in cerebellum, thalamus and neocortex. Front. Neural Circuits 7: 203.

Zhu, S., Noviello, C.M., Teng, J., Walsh, R.M., Jr, Kim, J.J., and Hibbs, R.E. (2018). Structure of a human synaptic GABAA receptor. Nature 559: 67–72.

Ziolkowski, L., Mordukhovich, I., Chen, D.M., Chisari, M., Shu, H.-J., Lambert, P.M., et al. (2020). A neuroactive steroid with a therapeutically interesting constellation of actions at GABA(A) and NMDA receptors. Neuropharmacology 183: 108358.

Zorumski, C.F., Covey, D.F., Izumi, Y., Evers, A.S., Maguire, J.L., and Mennerick, S.J. (2025). New directions in neurosteroid therapeutics in neuropsychiatry. Neurosci. Biobehav. Rev. 172: 106119.

Zuo, Y., Zhao, Y., Liu, G., and Sun, Q. (2026). Recent advances in GABAA receptor targeting ligands. Eur. J. Med. Chem. 308: 118651.

