## Supplemental file for "Subunit selective modulation of GABA_A_ receptors using pharmacogenetically tethered neurosteroids"

#### Supplementary figures:

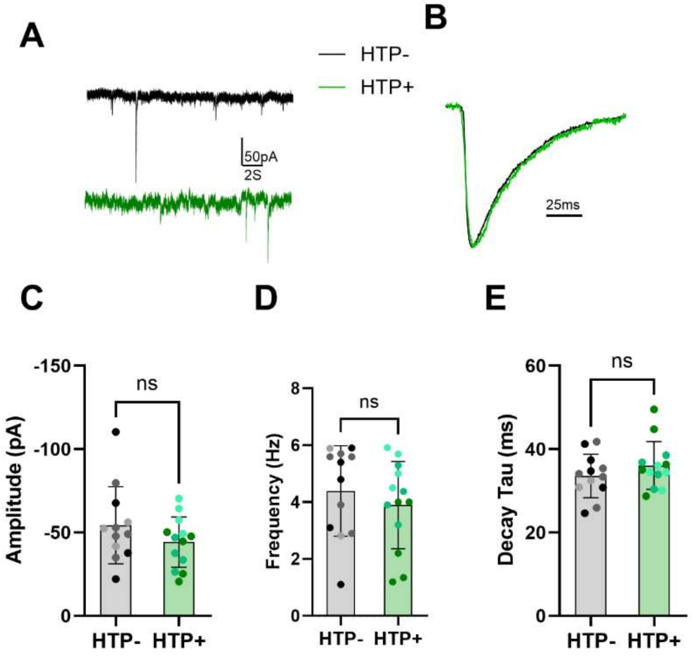


**Figure S1: HTP alone did not modulate IPSCs parameters.** (A) Representative raw traces of spontaneous IPSCs recorded from $HTP^-$ (black trace) and $HTP^+$ (green trace) hippocampal neurons (B) Representative traces of peak-normalized, average IPSC waveform from $HTP^-$ neurons (black trace) and $HTP^+$ neurons (green trace) and (C), (D) and (E) show summaries of amplitude and frequency and decay time constant respectively, derived from analysis of individual events. Student's $t$ test showed no significant difference, $p > 0.05$ for all three. Color shading of data points denote cells from independent platings.

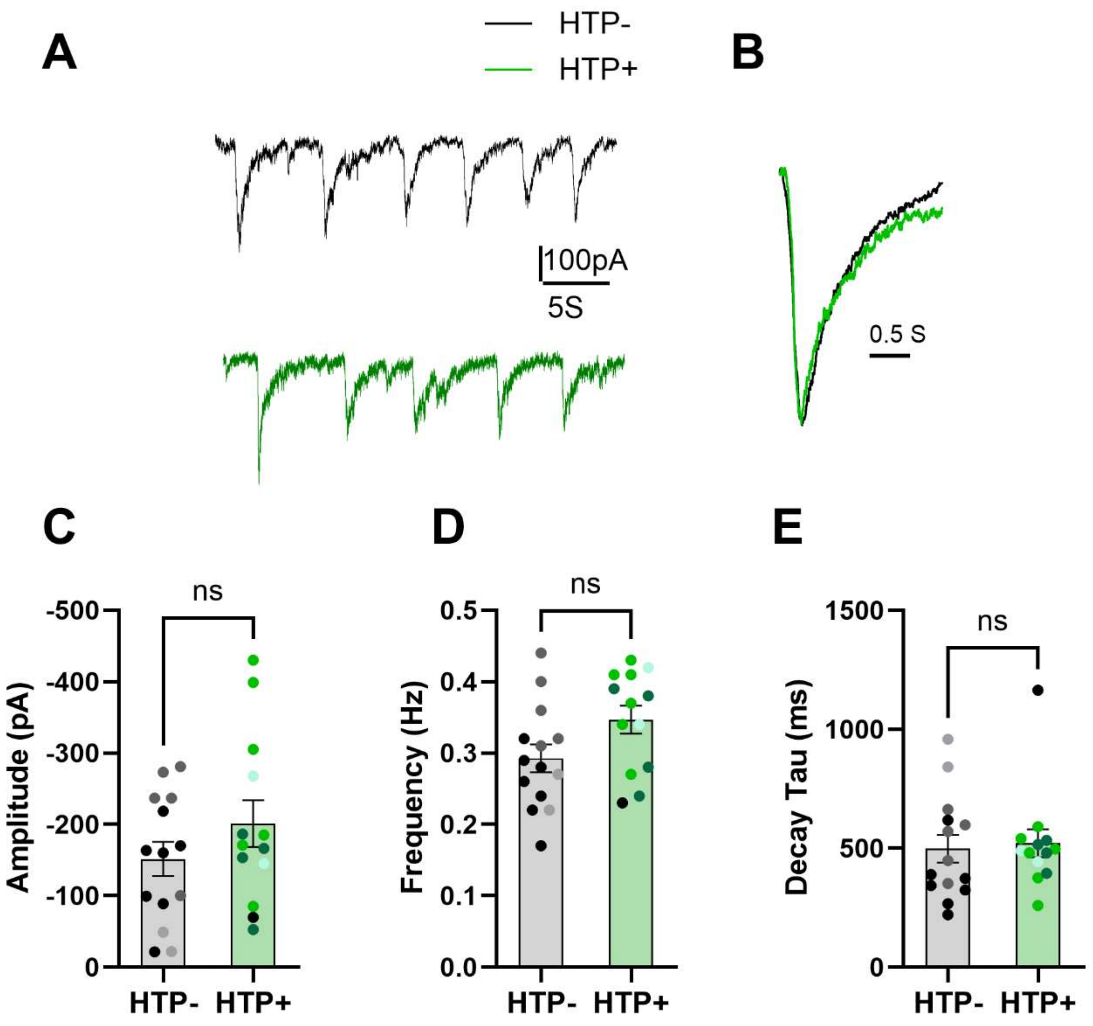


**Figure S2: Off-target effect of HTP transduction on NMDAR sEPSC parameters.** (A) Representative raw traces of spontaneous NMDAR EPSCs recorded from $HTP^-$ (black trace) and $HTP^+$ (green trace) hippocampal neurons (B) Representative traces of peak-normalized, average EPSC waveform from $HTP^-$ neurons (black trace) and $HTP^+$ neurons (green trace) and (C), (D) and (E) show summaries of amplitude, frequency, and decay time constant respectively, derived from analysis of individual events. Student's *t* test showed no significant difference, $p > 0.05$ for all three. Color shading of data points denote cells from independent culture.

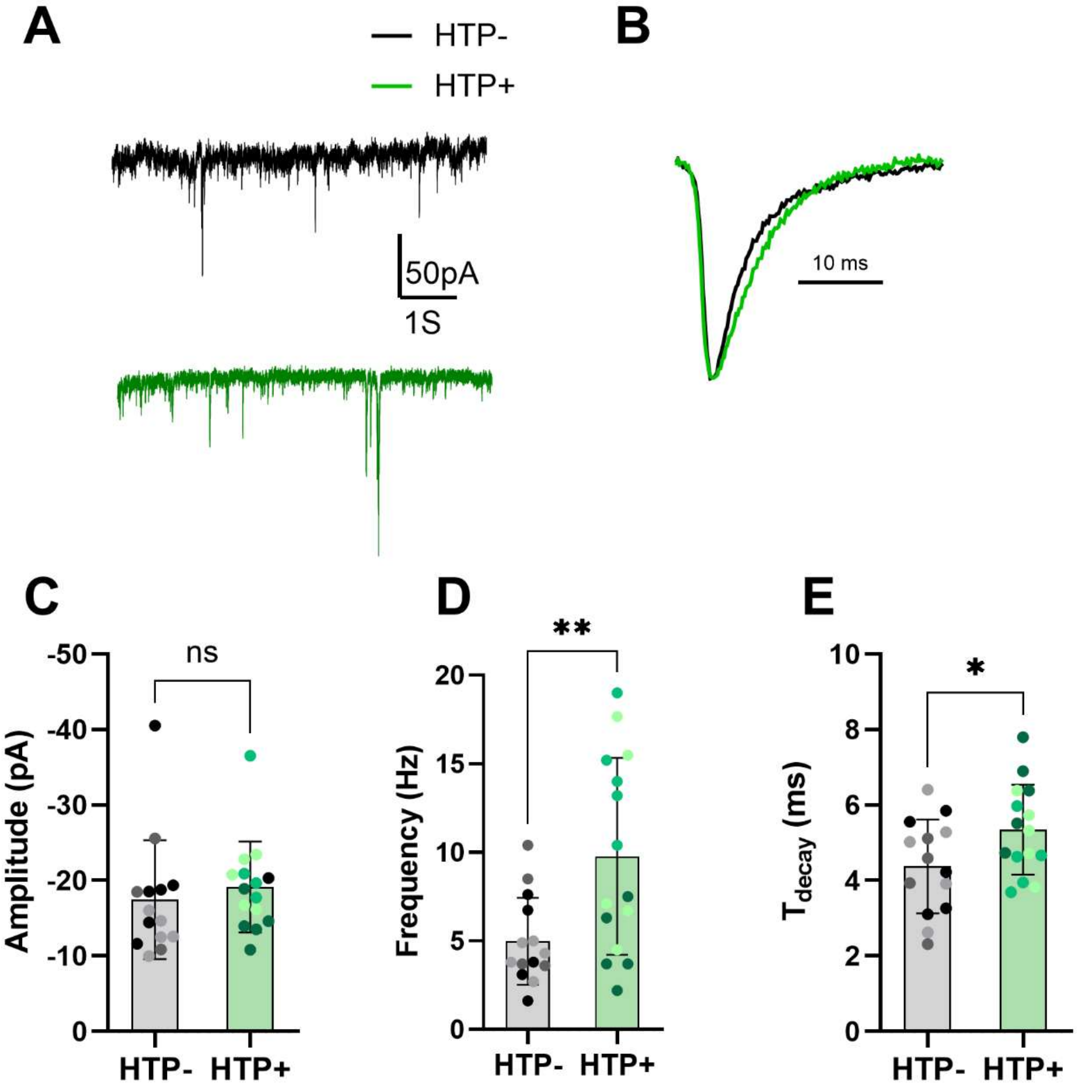


**Figure S3: Off-target effect of HTP transduction on AMPAR sEPSC parameters.** (A) Representative raw traces of spontaneous EPSCs recorded from HTP$^-$ (black trace) and HTP$^+$ (green trace) hippocampal neurons (B) Representative traces of peak-normalized, average EPSC waveform from HTP$^-$ neurons (black trace) and HTP$^+$ neurons (green trace) and (C), (D) and (E) show summaries of amplitude, frequency, and decay time constant respectively, derived from analysis of individual events. Student's *t* test showed $^*p < 0.05$, $^{**}p < 0.01$, for frequency and decay time constant. Color shading of data points denote cells from independent culture.

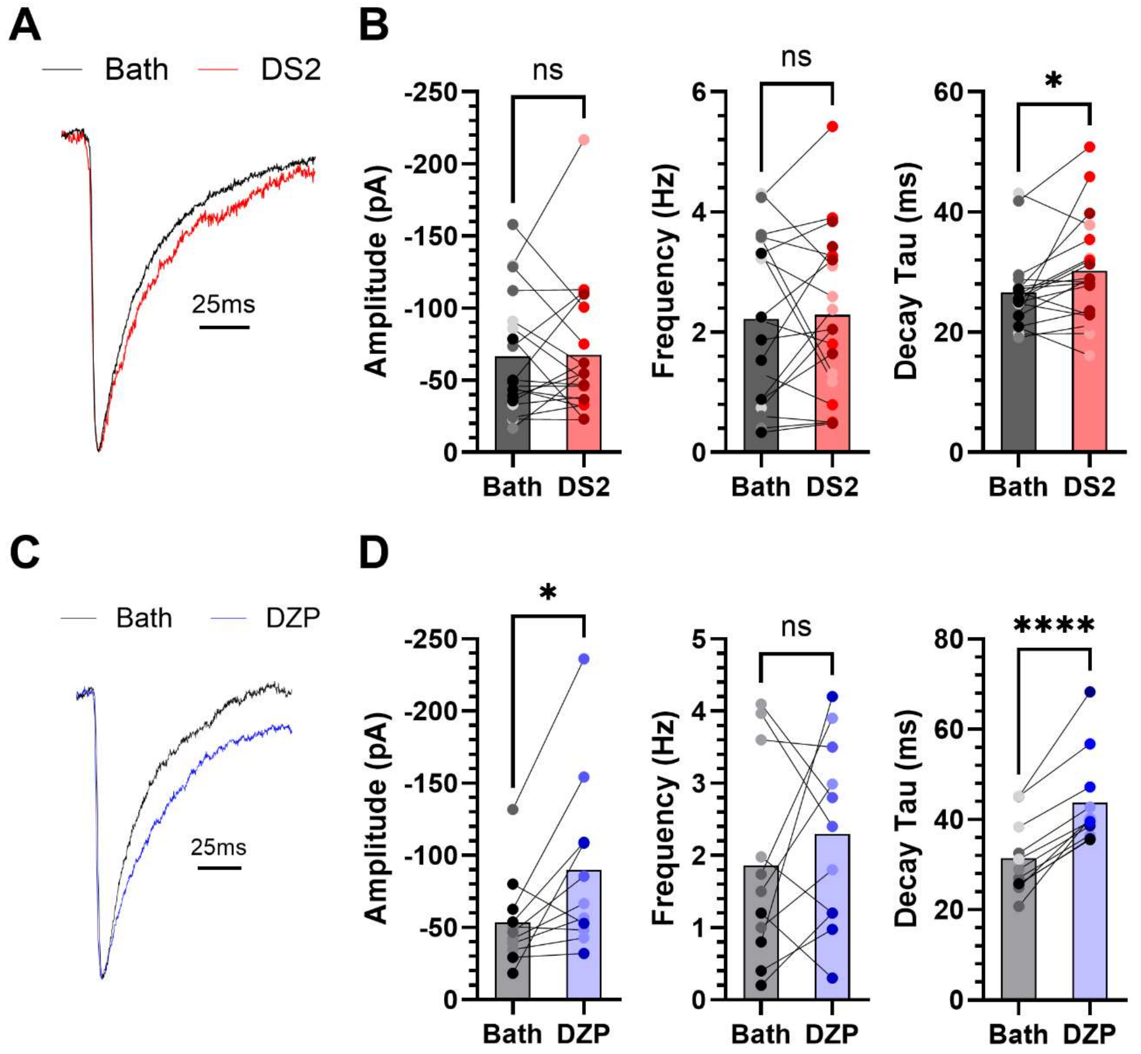


**Figure S4. Effect of a δ-selective and γ-selective PAM on sIPSCs in hippocampal cultures.** (A) Neurons were recorded at baseline and during application of 1 μM DS2. Representative average traces are shown, normalized to peak. (B) The indicated parameters were measured. Only decay time constant exhibited a reliable change with DS2 application (n=19). (C) Neurons were recorded at baseline and during application of 0.5 μM diazepam (DZP). Representative average traces normalize to peak are shown. (D) The indicated parameters were measured; amplitude and Decay tau exhibited a reliable change with DZP application (n=11). The ROUT method was used to identify outliers (two cells from DZP data set). Student's paired *t* test, *p<0.05, ***p<0.0001. Color shading of data points denote neurons from independent culture.

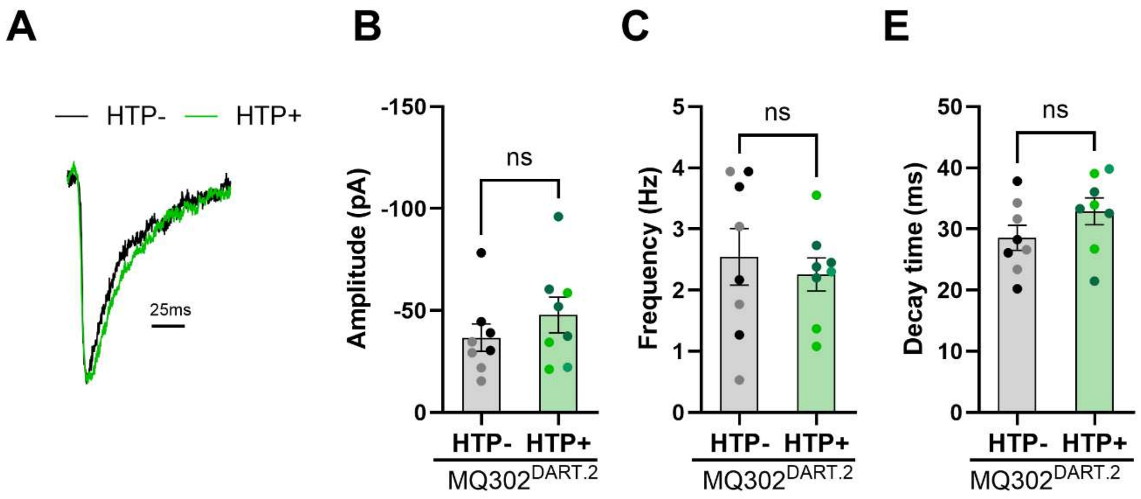


**Figure S5: Effect of MQ302$^{DART.2}$ on modulation of IPSCs in HTP neurons.** (A) Representative traces of peak-normalized, average IPSC waveform from HTP$^-$ neurons (black trace) and HTP$^+$ neurons (green trace) treated with **MQ302$^{DART.2}$**, (B), (C) and (D) show summaries of amplitude, frequency, and decay time constant respectively, derived from analysis of individual events. Student's *t* test showed no significant difference, $p > 0.05$. Color shading of data points denote cells from independent platings.

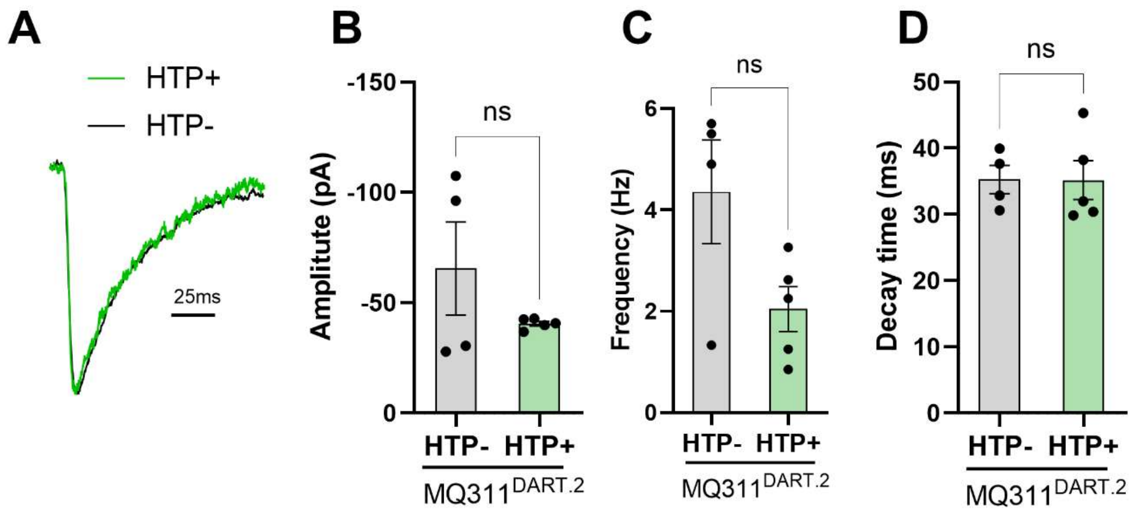


**Figure S6: Effect of MQ311$^{DART.2}$ on modulation of IPSCs in HTP neurons.** (A) Representative traces of peak-normalized, average IPSC waveform from HTP$^-$ neurons (black trace) and HTP$^+$ neurons (green trace) treated with **MQ311$^{DART.2}$**, (B), (C) and (D) show summaries of amplitude, frequency and decay time constant respectively, derived from analysis of individual events. Student's *t*-test no significant difference, $p > 0.05$.

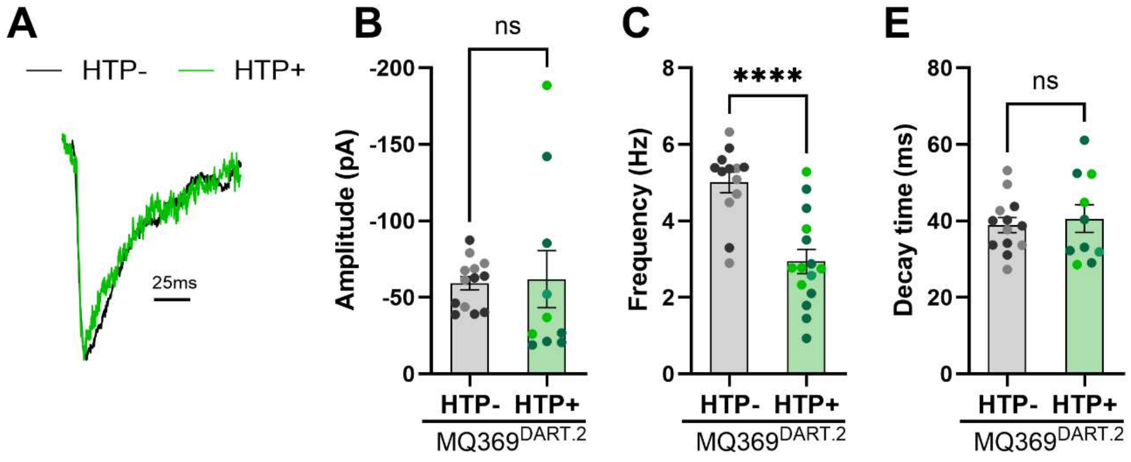


**Figure S7: Effect of MQ369$^{DART.2}$ on modulation of IPSCs in HTP neurons.** (A) Representative traces of peak-normalized, average IPSC waveform from HTP$^{-}$ neurons (black trace) and HTP$^{+}$ neurons (green trace) treated with **MQ369$^{DART.2}$**, (B), (C) and (D) show summaries of amplitude, frequency and decay time constant respectively, derived from analysis of individual events. Student's *t* test showed nonsignificant for amplitude and decay time constant p < 0.05 and frequency significantly changed (***p<0.05). Color shading of data points denote cells from independent platings.

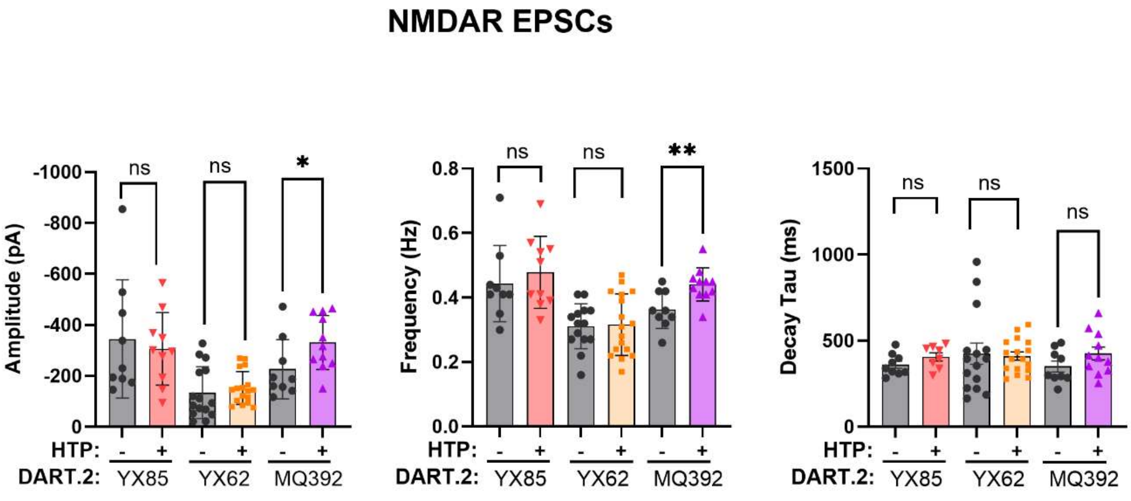


**Figure S8: Off-target activity of active NAS-DART on modulation of NMDAR EPSCs.** NMDAR mediated spontaneous EPSC recordings were performed in HTP$^{+}$ and HTP– neurons following application of NAS-DART compounds (e.g. **YX85.1$^{DART.2}$, YX62$^{DART.2}$, MQ392$^{DART.2}$**), each compound applied separately. Bar graphs summarize the mean ± SEM of EPSC (A) amplitude, (B) event frequency, and (C) decay time constants derived from analysis of individual synaptic events. Statistical comparisons were performed using Student's *t* test. Differences were considered not significant when $p \geq 0.05$ and significant when $*p < 0.05$, $**p < 0.01$.

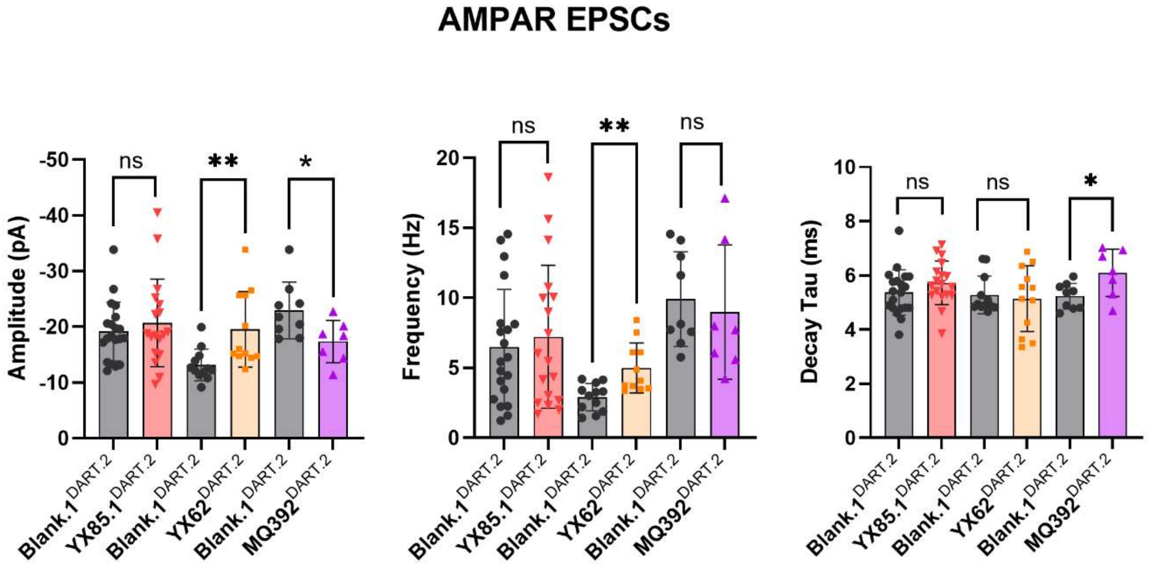


**Figure S9: Off-target activity of active NAS-DART on modulation of AMPAR EPSCs in HTP+ neurons.** AMPAR mediated spontaneous EPSC recordings were performed selectively in $HTP^+$ neurons following application of either **blank.1$^{DART.2}$** or NAS-DART compounds (e.g. **YX85.1$^{DART.2}$, YX62$^{DART.2}$, MQ392$^{DART.2}$**), each compound applied separately. Bar graphs summarize the mean ± SEM of EPSC (A) amplitude, (B) event frequency, and (C) decay time constants derived from analysis of individual synaptic events. Statistical comparisons were performed using Student's *t* test. Differences were considered not significant when $p \geq 0.05$ and significant when $*p < 0.05$, $**p < 0.01$.

**Synthesis of neurosteroid analogues:**

**Synthesis of MQ368 and MQ369**

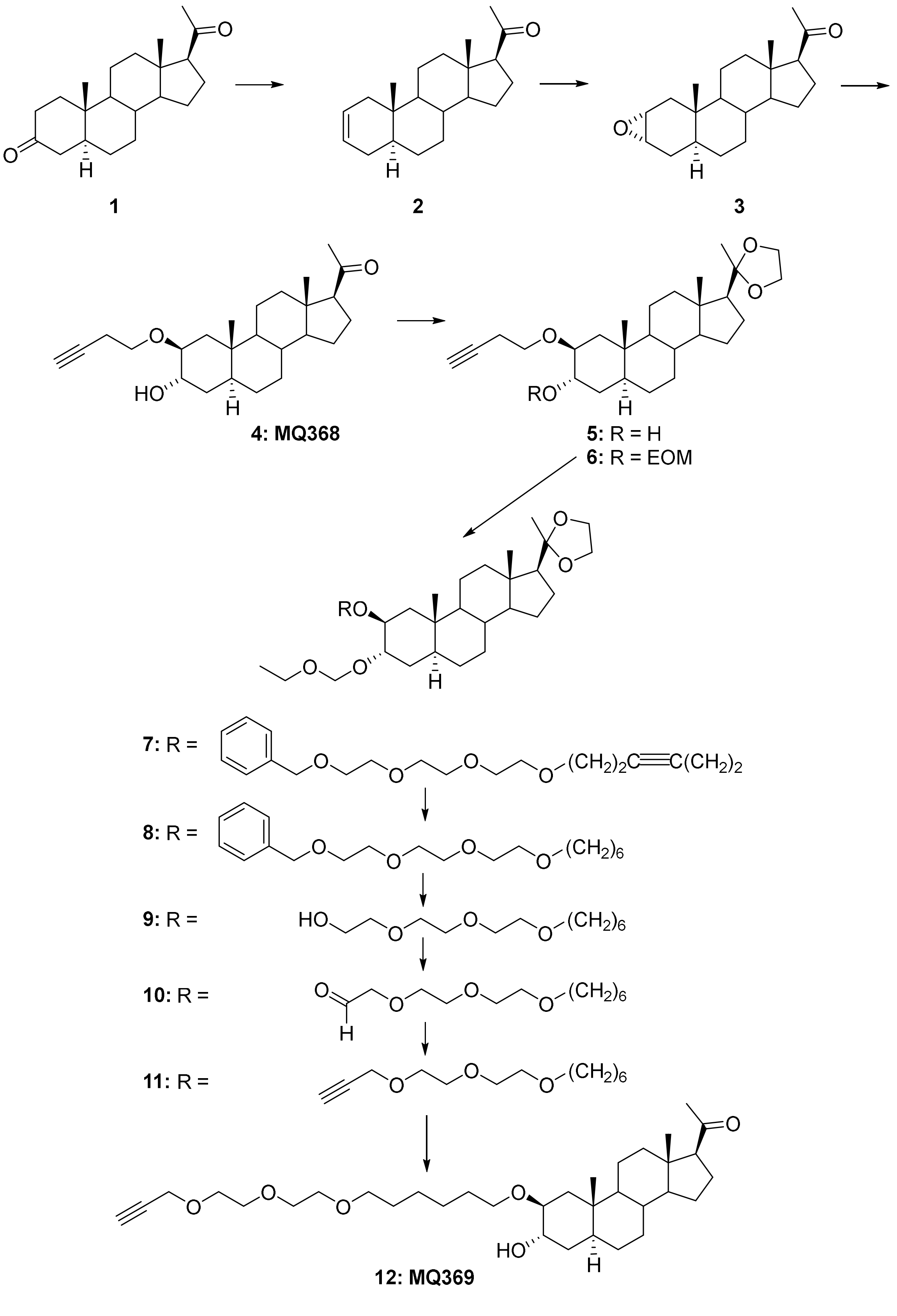

**5α-Pregn-2-en-20-one (2).** To a suspension of zinc dust (8.23 g, 126.6 mmol) and TMSCl (8.03 mL, 63.3 mmol) in THF (100 mL) was added 5α-pregnan-3,20-dione (**1**, 2.0 g, 6.33 mmol ) at 23 °C. The reaction mixture was refluxed for 16 h. After cooling, the mixture was filtered through Celite and washed with THF (50 mL). The filtrate was washed with aqueous $NaHCO_3$, dried over anhydrous $Na_2SO_4$, filtered and the solvent removed. The residue was purified by flash column chromatography (silica gel, eluted with 5% EtOAc in hexanes) to give steroid **2** which also contained the $\Delta^3$ isomer (1.43 g, $\Delta^2/\Delta^3 = 6/1$, 75%), steroid **2** has: $^1$H NMR (400 MHz, $CDCl_3$) δ 5.58-5.56 (m, 2H), 2.53 (t, $J$ = 8.9 Hz, 1H), 2.15-0.75 (m, 20H), 2.10 (s, 3H), 0.73 (s, 3H), 0.59 (s, 3H); $^{13}$C NMR (100 MHz, $CDCl_3$) δ 209.6, 125.7, 125.6, 63.7, 56.6, 53.8, 44.0, 41.3, 39.6, 38.9, 35.4, 34.5, 31.6, 31.5, 30.1, 28.5, 24.3, 22.6, 20.8, 13.3, 11.6.

**2α,3α-Epoxy-5α-pregnan-20-one (3).** To a solution of steroid **2** containing the $\Delta^3$ isomer (1.43 g, 4.83 mmol) in $CH_2Cl_2$ (40 mL) was added formic acid (10 mL) and hydrogen peroxide (10 mL) at 23 °C. After 4 h, the product was extracted into $CH_2Cl_2$ (200 mL). The extract was washed with $NaHCO_3$ (50 mL), dried over anhydrous $Na_2SO_4$, filtered and the solvent removed. The residue was purified by flash column chromatography (silica gel, eluted with 10% EtOAc in hexanes) to give steroid **3** (1.23 g, 81%): $^1$H NMR (400 MHz, $CDCl_3$) δ 3.16-3.10 (m, 2H), 2.54 (t, $J$ = 9.0 Hz, 1H), 2.18-0.59 (m, 20H), 2.10 (s, 3H), 0.75 (s, 3H), 0.58 (s, 3H); $^{13}$C NMR (100 MHz, $CDCl_3$) δ 209.6, 63.7, 56.5, 53.5, 52.4, 51.0, 44.0, 38.9, 38.2, 36.2, 35.6, 33.6, 31.6, 31.5, 29.0, 28.3, 24.4, 22.7, 20.9, 13.3, 12.9.

**2β-(But-3-yn-1-yloxy)-3α-hydroxy-5α-pregnan-20-one (4, MQ368).** To a stirred solution of steroid **3** (1.23 g, 3.94 mmol) in 3-butyn-1-ol (10 mL) was added tetracyanoethylene (1.03 g, 8 mmol) at 23 °C. After 72 h, 3-butyn-1-ol was removed under reduced pressure and the residue was purified by flash column chromatography (silica gel, eluted with 25-35% EtOAc in hexanes) to give steroid **MQ368** (**4**, 720 mg, 47%): $^1$H NMR (400 MHz, $CDCl_3$) δ 3.90 (s, 1H), 3.63-3.57 (m, 1H), 3.47-3.41 (m, 2H), 2.50 (t, $J$ = 8.9 Hz, 1H), 2.40-0.71 (m, 24H), 2.07 (s, 3H), 0.89 (s, 3H), 0.55 (s, 3H); $^{13}$C NMR (100 MHz, $CDCl_3$) δ 209.9, 81.4, 79.0, 69.1, 68.2, 66.9, 63.7, 56.6, 54.7, 44.2, 38.9, 38.8, 36.1, 35.7, 34.7, 31.9, 31.8, 31.4, 27.9, 24.2, 22.6, 20.8, 20.0, 13.3, 13.2.

**2β-(But-3-yn-1-yloxy)-17β-(2-methyl-1,3-dioxolan-2-yl)-5α-androstan-3α-ol (5).** To a stirred solution of steroid **4** (700 mg, 1.83 mmol) in benzene (150 mL) was added ethylene glycol (2 mL) and PTSA (50 mg) at 23 °C. The reaction was refluxed under $N_2$ in a flask equipped with a Dean-Stark apparatus for 16 h. After cooling, solid $NaHCO_3$ (300 mg) was added and stirring was continued for 10 min. Aqueous $NaHCO_3$ (100 mL) was added and the product was extracted into EtOAc (300 mL). The extract was washed with brine (100 mL x3), dried over anhydrous $Na_2SO_4$, filtered and the solvent removed. The residue was purified by flash column chromatography (silica gel, eluted with 25% EtOAc in hexanes) to give steroid **5** (705 mg, 90%): $^1$H NMR (400 MHz, $CDCl_3$) δ 3.99-3.83 (m, 5H), 3.65-3.60 (m, 1H), 3.49-3.43 (m, 2H), 2.43-2.38 (m, 2H), 2.03- 0.58 (m, 23H), 1.27 (s, 3H), 0.91 (s, 3H), 0.73 (s, 3H); $^{13}$C NMR (100 MHz, $CDCl_3$) δ 111.9, 81.5, 79.1, 69.1, 68.5, 67.0, 65.1, 63.1, 58.2, 56.3, 55.0, 42.0, 39.6, 38.9, 36.2, 35.8, 34.3, 32.1, 31.8, 28.1, 24.5, 23.6, 22.8, 20.6, 20.1, 13.3, 13.0.

**2β-(But-3-yn-1-yloxy)-3α-(ethoxymethoxy)-17β-(2-methyl-1,3-dioxolan-2-yl)-5α-androstane (6).** To a stirred solution of steroid **5** (705 mg, 1.66 mmol) in $CH_2Cl_2$ (20 mL) was added chloromethyl ethyl ether (0.31 mL, 3,3 mmol) and (*i*-Pr)$_2$NEt (0.87 mL, 5 mmol) at 23 °C. After 16 h, solvent was removed and the residue was purified by flash column chromatography (silica gel, eluted with 10-20% EtOAc in hexanes) to give steroid **6** (795 mg, 99%): $^1$H NMR (400 MHz, $CDCl_3$) δ 4.69 (s, 2H), 3.98-3.82 (m, 4H), 3.75 (s, 1H), 3.62-3.45 (m 5H), 2.42-2.37 (m, 2H), 2.09-0.68 (m, 25H), 1.26 (s, 3H), 0.91 (s, 3H), 0.72 (s, 3H); $^{13}$C NMR (100 MHz, $CDCl_3$) δ 111.9, 83.9, 81.4, 77.4, 73.8, 69.1, 67.0, 65.1, 63.1 (2 x C), 58.2, 56.3, 54.9, 42.0, 39.6, 39.5, 36.7, 35.5, 34.3, 31.8, 29.7, 28.1, 24.5, 23.6, 22.8, 20.6, 20.1, 15.1, 13.2, 13.0.

**3α-(Ethoxymethoxy)-17β-(2-methyl-1,3-dioxolan-2-yl)-2β-((1-phenyl-2,5,8,11-tetraoxaheptadec-14-yn-17-yl)oxy)-5α-androstane (7).** To a mixture of steroid **6** (795 mg, 1.64 mmol), 13-iodo-1-phenyl-2,5,8,11-tetraoxatridecane (840 mg, 2.13 mmol), bis[(2-dimethylamino)phenyl]amine nickel(II) chloride (57 mg, 0.164 mmol), Cu(I)I (19 mg, 0.1 mmol) and $Cs_2CO_3$ (748 mg, 2.30 mmol) was added anhydrous dioxane (25 mL) at 23 °C. The reaction was heated to 120 °C for 16 h. After cooling, dioxane was removed under reduced pressure and the residue was purified by flash column chromatography (silica gel, eluted with 25-40% EtOAc in hexanes) to give steroid **7** (471 mg, 37%): $^1$H NMR (400 MHz, $CDCl_3$) δ 7.30-7.24 (m, 5H), 4.67 (s, 2H), 4.54 (s, 2H), 3.96-3.38 (m, 26H), 2.42-0.88 (m, 26H), 1.26 (s, 3H), 0.90 (s, 3H), 0.71

(s, 3H); $^{13}C$ NMR (100 MHz, $CDCl_3$) δ 138.2, 128.3 (2 x C), 127.7 (2 x C), 127.6, 111.9, 93.9, 78.1, 77.6, 77.3, 73.9, 73.2, 70.6 (3 x C), 70.5, 70.2, 69.9, 69.4, 67.5, 65.2, 63.2, 58.3, 56.4, 55.0, 42.0, 39.7, 39.6, 36.7, 35.6, 34.4, 31.9, 29.8, 28.2, 24.6, 23.7, 22.9, 20.7, 20.5, 20.0, 15.2, 13.3, 13.1.

**3α-(Ethoxymethoxy)-17β-(2-methyl-1,3-dioxolan-2-yl)-2β-((1-phenyl-2,5,8,11-tetraoxaheptadecan-17-yl)oxy)-5α-androstane (8).** To a solution of steroid **7** (471 mg, 0.604mmol) in EtOAc (50 mL) was added Pd/C (300 mg) at 23 °C in a Parr hydrogenation flask. The flask was evacuated and charged with $H_2$ three times and hydrogenation was carried out under 55 psi $H_2$ overnight. The mixture was filtered through Celite and washed with EtOAc (100 mL). The solvent was removed and the residue was purified by flash column chromatography (silica gel, eluted with 25-50% EtOAc in hexanes) to give steroid **8** (350 mg, 74%): $^{1}H$ NMR (400 MHz, $CDCl_3$) δ 7.31-7.23 (m 5H), 4.67 (s, 2H), 4.53 (s, 2H), 3.95-3.40 (m, 26H), 1.83-0.75 (m, 30H), 1.25 (s, 3H), 0.89 (s, 3H), 0.71 (s, 3H); $^{13}C$ NMR (100 MHz, $CDCl_3$) δ 138.1, 128.2 (2 x C), 127.6 (2 x C), 127.4, 111.8, 93.7, 77.0, 73.7, 73.1, 71.3, 70.5 (2 x C), 70.4, 69.9, 69.3, 68.8, 65.0, 63.0, 62.9, 58.2, 56.3, 54.9, 41.9, 39.6, 39.5, 36.8, 35.5, 34.2, 31.8, 30.0, 29.7, 29.4, 28.1, 26.0, 25.8, 24.4, 23.6, 22.8, 20.5, 15.0, 13.2, 12.9.

**2-(2-(2-((6-((-3α-(Ethoxymethoxy)-17β-(2-methyl-1,3-dioxolan-2-yl)-5α-androstan-2β-yl)oxy)hexyl)oxy)ethoxy)ethoxy)-ethan-1-ol (9).** To a solution of anhydrous liquid ammonia (50 mL) was added sodium metal (230 mg, 10 mmol) at -78 °C. The reaction was stirred for 15 min. Steroid **8** (350 mg, 0.448 mmol) in THF (30 mL) was added over 15 min and the reaction stirred for additional 1 h. Solid $NaHCO_3$ was added and the reaction was warmed to 23 °C and the ammonia allowed to evaporate overnight. Aqueous $NaHCO_3$ was added and the product was extracted into EtOAc (200 mL x2). The combined extracts were dried over anhydrous $Na_2SO_4$, filtered, the solvent removed and the residue purified by flash column chromatography (silica gel, eluted with 25-50% EtOAc in hexanes) to give steroid **9** (278 mg, 93%): $^{1}H$ NMR (400 MHz, $CDCl_3$) δ 4.66 (s, 2H), 3.94-3.24 (m 26H), 2.84 (br, 1H), 1.96-0.60 (m, 30H), 1.23 (s, 3H), 0.87 (s, 3H), 0.68 (s, 3H); $^{13}C$ NMR (100 MHz, $CDCl_3$) δ 111.8, 93.6, 77.0, 73.7, 72.4, 71.3, 70.4, 70.3, 70.2, 69.8, 68.7, 65.0, 63.0, 62.9, 61.5, 58.1, 56.2, 54.8, 41.8, 39.5, 39.4, 36.7, 35.5, 34.2, 31.7, 29.9, 29.6, 29.3, 28.0, 26.0, 25.7, 24.4, 23.6, 22.7, 20.5, 15.0, 13.1, 12.9.

**2-(2-(2-((6-((-3α-(Ethoxymethoxy)-17β-(2-methyl-1,3-dioxolan-2-yl)-5α-androstan-2β-yl)oxy)hexyl)oxy)ethoxy)ethoxy)-acetaldehyde (10).** To a stirred solution of oxalyl chloride (0.25 mL, 3 mmol) in $CH_2Cl_2$ (15 mL) was added DMSO (0.32 mL, 4.5 mmol) at -78 °C. After 15 min, steroid **9** (278 mg, 0.402 mmol) in $CH_2Cl_2$ (8 mL) was added and the reaction was stirred at -78 °C for 1 h. $Et_3N$ (0.84 mL, 6 mmol) was added at -78 °C and stirred for 30 min. The mixture was warmed to 23 °C for an additional 30 min. Water (50 mL) was added and the product was extracted into $CH_2Cl_2$ (150 mL x2). The combined extracts were dried over anhydrous $Na_2SO_4$, filtered, the solvent removed and the residue purified by flash column chromatography (silica gel, eluted with 25-50% EtOAc in hexanes) to give steroid **10** (222 mg, 80%): $^1$H NMR (400 MHz, $CDCl_3$) δ 9.67 (s, 1H), 4.64 (s, 2H), 4.11 (s, 2H), 3.94-3.35 (m, 22H), 1.96-0.64 (m, 30H), 1.22 (s, 3H), 0.86 (s, 3H), 0.68 (s, 3H); $^{13}$C NMR (100 MHz, $CDCl_3$) δ 200.7, 111.8, 93.6, 77.0, 76.7, 73.6, 71.2, 71.0, 70.6, 70.5, 70.4, 69.8, 68.7, 65.0, 63.0, 62.9, 58.1, 56.2, 54.8, 41.8, 39.5, 36.8, 35.5, 34.2, 31.7, 29.9, 28.6, 29.4, 28.0, 26.0, 25.7, 24.4, 23.5, 22.7, 20.5, 15.0, 13.1, 12.9.

**2-(3α-(Ethoxymethoxy)-2β-((6-(2-(2-(prop-2-yn-1-yloxy)ethoxy)ethoxy)hexyl)oxy)-5α-androstan-17β-yl)-2-methyl-1,3-dioxolane (11).** To a stirred solution of steroid **10** (222 mg, 0.322 mmol) in THF (5 mL) and MeOH (5mL) was added dimethyl-1-diazo-2-oxopropyl phosphonate (0.31 mL, 2 mmol) and a fine powder of $K_2CO_3$ (552 mg, 4 mmol) at 23 °C. After 16 h, water was added and the mixture was extracted into EtOAc (100 mL x2). The combined extracts were dried over anhydrous $Na_2SO_4$, filtered, the solvent removed and the residue purified by flash column chromatography (silica gel, eluted with 25-40% EtOAc in hexanes) to give steroid **11** (182 mg, 83%): $^1$H NMR (400 MHz, $CDCl_3$) δ 4.65 (s, 2H), 4.16 (s, 2H), 3.94-0.64 (m, 20H), 2.40 (s, 1H), 1.97-0.64 (m, 32H), 1.23 (s, 3H), 0.87 (s, 3H), 0.67 (s, 3H); $^{13}$C NMR (100 MHz, $CDCl_3$) δ 111.8, 93.7, 79.4, 77.0, 74.4, 73.7, 71.2, 70.5, 70.3, 70.0, 68.9, 68.7, 65.0, 63.0, 62.9, 58.2, 58.1, 56.2, 54.9, 41.9, 39.5, 39.4, 36.8, 35.5, 34.2, 31.7, 29.9, 29.6, 29.4, 28.1, 26.0, 25.8, 24.4, 23.5, 22.7, 20.5, 15.0, 13.1, 12.9.

**3α-(Hydroxy)-2β-((6-(2-(2-(prop-2-yn-1-yloxy)ethoxy)ethoxy)hexyl)oxy)-5α-pregnan-20-one (12, MQ369).** To a stirred solution of steroid **11** in THF (10 mL) was added 6 N HCl (10 mL) at 23 °C. After 2 h, solid $NaHCO_3$ was slowly added to reach pH~8. The product was extracted into EtOAc (150 mL x3). The combined extracts were dried over anhydrous $Na_2SO_4$, filtered, the solvent removed and the residue purified by flash column chromatography (silica gel, eluted with

25-40% EtOAc in hexanes) to give **MQ369** (**12**, 182 mg, 83%): $^{1}$H NMR (400 MHz, $CDCl_3$) δ 4.19 (s, 2H), 3.89 (s, 1H), 3.70-3.28 (m, 14H), 2.52 (t, *J* = 8.9, 1H), 2.42 (s, 1H), 2.13-0.72 (m, 28H), 2.09 (s, 3H), 0.90 (s, 3H), 0.57 (s, 3H); $^{13}$C NMR (100 MHz, $CDCl_3$) δ 209.9, 79.6, 78.9, 74.6, 71.4, 70.6, 70.4, 70.0, 69.1, 68.9, 68.4, 63.8, 58.4, 56.7, 54.9, 44.3, 39.1, 39.0, 36.3, 35.9, 34.9, 32.1, 31.9, 31.5, 30.0, 29.5, 28.0, 26.1, 25.9, 24.3, 22.7, 20.9, 13.5, 13.3.

**Synthesis of MQ297 andMQ311**

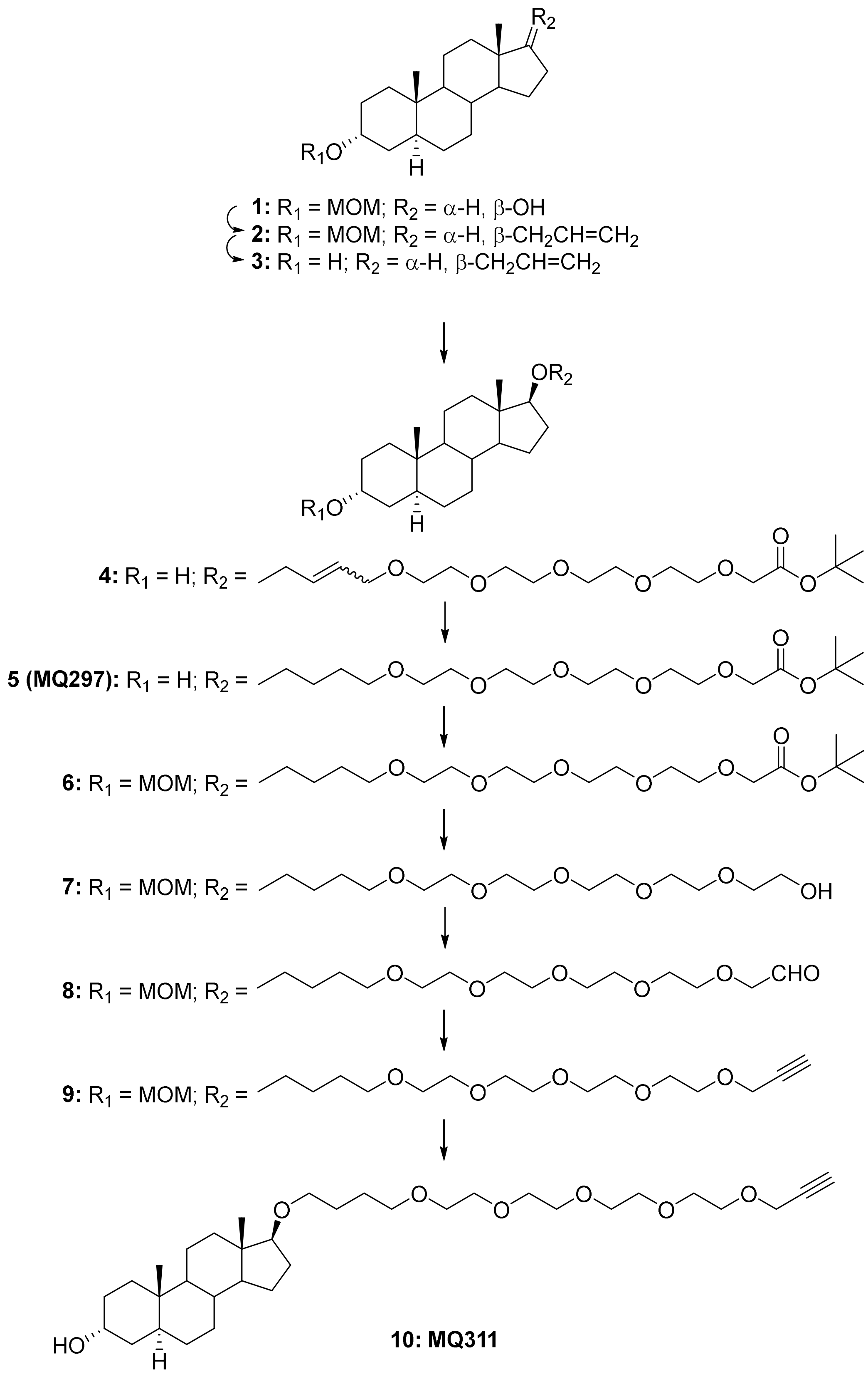

**3α-(Methoxymethoxy)-17β-(2-propenyloxy)-5α-androstane (2).** 3α-(Methoxymethoxy)-5α-androstan-17β-ol (**1**) was prepared according to the literature (Chintala, S.M. et al. *Br. J. Pharmacol.* 2024**, 181**, 4229-4244). To a stirred solution of steroid **1** (755 mg, 2.25 mmol) in THF (40 mL) was added NaH (60% in mineral oil, 680 mg, 17 mmol) at 23 °C. The mixture was refluxed for 1 h. Allyl bromide (1.93 mL, 22.5 mmol) was added and the mixture was refluxed for an additional 16 h. After cooling, aqueous $NH_4Cl$ was added and the product was extracted into EtOAc (150 mL x 2). The combined extracts were dried over anhydrous $Na_2SO_4$, filtered, solvent removed and the residue purified by flash column chromatography (silica gel, eluted with 10% EtOAc in hexanes) to give steroid **2** (811 mg, 96%): $^1$H NMR (400 MHz, $CDCl_3$) δ 5.89-5.82 (m, 1H), 5.23-5.19 (m, 2H), 4.61 (s, 2H), 3.95-3.94 (m, 2H), 3.78 (s, 1H), 3.32 (s, 3H), 3.33-3.30 (m, 1H), 1.96-0.68 (m, 22H), 0.76 (s, 3H), 0.73 (s, 3H); $^{13}$C NMR (100 MHz, $CDCl_3$) δ 135.5, 115.7, 94.3, 88.4, 71.4, 70.6, 54.9, 54.3, 51.2 42.9, 39.6, 37.9, 35.8, 35.2, 33.5, 32.7, 31.4, 28.4, 27.9, 26.2, 23.2, 20.3, 11.6, 11.3.

**17β-(2-Propenyloxy)-5α-androstan-3α-ol (3).** To a stirred solution of steroid **2** (811 mg, 2.16 mmol) in MeOH (30 mL) was added acetyl chloride (4 mL) at 23 °C. After 2 h, the product was extracted into $CH_2Cl_2$ (150 mL x 2). The combined extracts were washed with aqueous $NaHCO_3$ (50 mL), dried over anhydrous $Na_2SO_4$, filtered, solvent removed and the residue was purified by flash column chromatography (silica gel, eluted with 25% EtOAc in hexanes) to give steroid **3** (679 mg, 95%): $^1$H NMR (400 MHz, $CDCl_3$) δ 5.94-5.86 (m, 1H), 5.28-5.11 (m, 2H), 4.04-3.98 (m, 3H), 3.37 (t, $J$ = 8.2 Hz, 1H), 2.00-0.65 (m, 23H), 0.78 (s, 3H), 0.77 (s, 3H); $^{13}$C NMR (100 MHz, $CDCl_3$) δ 135.7, 116.0, 88.5, 70.8, 66.5, 54.5, 51.3, 43.0, 39.1, 38.0, 36.1, 35.9, 35.3, 32.2, 31.5, 29.0, 28.4, 28.0, 23.3, 20.4, 11.8, 11.2.

***tert*-Butyl 19-((3α-hydroxy-5α-androstan-17β-yl)oxy)-3,6,9,12,15-pentaoxanonadec-17-enoate (4).** To a stirred solution of steroid **3** (679 mg, 2.05 mmol) and *tert*-butyl 3,6,9,12,15-pentaoxaoctadec-17-enoate (3.48 g, 10 mmol) in THF (6 mL) was added Hoveyda-Grubbs catalyst M2001 (32 mg, 0.05 ) and Grubbs catalyst second generation (43 mg, 0.05 mmol) at 23 °C. After 16 h, the solvent was removed and the residue was purified by flash column chromatography (silica gel, eluted with 25% EtOAc in hexanes) to give steroid **4** (430 mg, *E* and *Z* mixture, not pure): $^1$H NMR (400 MHz, $CDCl_3$) δ 6.37-5.78 (m, 1H), 5.36-4.72 (m, 1H), 4.52-4.18 (m, 1H), 4.02 (s, 4H), 3.83-3.22 (m, 22H), 2.31-0.63 (m, 29H), 0.77 (s, 3H), 0.73 (s, 3H); $^{13}$C NMR (100 MHz, $CDCl_3$)

δ 169.6, 147.2, 103.3, 89.2, 89.0, 81.5, 71.3, 70.7, 70.6, 70.3, 69.7, 69.4, 69.0, 68.1, 66.5, 54.4, 51.3, 43.0, 39.1, 38.1, 36.1, 35.9, 35.3, 32.2, 31.5, 29.0, 28.8, 28.4, 28.1(3 x C), 25.2, 23.3, 20.4, 11.8, 11.7, 11.4, 11.2, 9.6.

***tert*-Butyl 19-((3α-hydroxy-5α-androstan-17β-yl)oxy)-3,6,9,12,15-pentaoxanonadecanoate (5, MQ297).** To a solution of steroid **4** (430 mg pure) in EtOAc (45 mL) in a Parr hydrogenation flask was added Pd/C (100 mg) at 23 °C. The flask was evacuated and charged with $H_2$ three times. The hydrogenation was carried out at 55 psi $H_2$ overnight. The mixture was filtered through Celite and washed with EtOAc (100 mL). The solvent was removed and the residue was purified by flash column chromatography (silica gel, eluted with 25-50% EtOAc in hexanes) to give steroid **5** (132 mg, 31%): $^1$H NMR (400 MHz, $CDCl_3$) δ 4.01 (s, 3H), 3.71-3.23 (m, 21H), 1.96-0.69 (m, 27H), 1.45 (s, 9H), 0.76 (s, 3H), 0.72 (s, 3H); $^{13}$C NMR (100 MHz, $CDCl_3$) δ 169.6, 89.0, 81.4, 71.2, 70.7, 70.5 (4 x C), 70.0, 69.7, 69.0, 66.4, 54.4, 51.3, 43.0, 39.1, 38.1, 36.1, 35.8, 35.3, 32.1, 31.5, 28.9, 28.4, 28.0 (6 x C), 26.7, 26.4, 23.3, 20.4, 11.7, 11.1.

***tert*-Butyl 19-(((3α-methoxymethoxy)-5α-androstan-17β-yl) oxy)-3,6,9,12,15-pentaoxanonadecanoate (6).** To a stirred solution of steroid **5** (120 mg, 0.183 mmol) in $CH_2Cl_2$ (10 mL) was added chloromethyl methyl ether (2 mmol) and (*i*-Pr)$_2$NEt (0.42 mL, 3 mmol) at 23 °C. After 24 h, the solvent was removed and the residue was purified by flash column chromatography (silica gel, eluted with 30-50% EtOAc in hexanes) to give steroid **6** (118 mg, 92%): $^1$H NMR (400 MHz, $CDCl_3$) δ 4.62 (q, $J$ = 6.6 Hz, 2H), 3.98 (s, 2H), 3.79 (s, 1H), 3.32 (s, 3H), 3.66-3.22 (m, 20H), 2.30-0.72 (m, 27H), 1.43 (s, 9H), 0.74 (s, 3H), 0.68 (s, 3H); $^{13}$C NMR (100 MHz, $CDCl_3$) δ 169.5, 94.3, 89.0, 81.3, 71.4, 71.1, 70.5, 70.43 (3 x C), 70.41 (3 x C), 70.4, 69.9, 69.6, 68.8, 55.0, 54.3, 51.2, 42.9, 39.6, 38.0, 35.8, 35.1, 33.5, 32.7, 31.4, 28.3, 27.9 (3 x C), 26.7, 26.3, 26.1, 23.2, 20.3, 11.6, 11.3.

**19-((3α-(Methoxymethoxy)-5α-androstan-17β-yl)oxy)-3,6,9,12,15-pentaoxanonadecan-1-ol (7).** To a stirred solution of steroid **6** (118 mg, 0.169 mmol) in diethyl ether (30 mL) was added lithium aluminum hydride (2.0 M in diethyl ether, 5 mL, 10 mmol) at 23 °C. After 2 h, water (0.4 mL), 10% NaOH (0.8 mL), and water (1.2 mL) were slowly added sequentially. After stirring for 1 h, the mixture was filtered through Celite and washed with EtOAc (200 mL). The solvent was removed and the residue was purified by flash column chromatography (silica gel, eluted with 50% EtOAc in hexanes) to give steroid **7** (101 mg, 95%): $^1$H NMR (400 MHz, $CDCl_3$) δ 4.63 (q,

$J$ = 6.7 Hz, 2H), 3.78 (s, 1H), 3.32 (s, 3H), 3.66-3.22 (m, 25H), 1.93-0.72 (m, 27H), 0.75 (s, 3H), 0.69 (s, 3H); $^{13}$C NMR (100 MHz, $CDCl_3$) δ 94.3, 89.0, 72.5, 71.5, 71.1, 70.4, 70.4, 70.4, 70.3 (2 x C), 70.3, 70.1, 69.8, 69.7, 61.5, 55.0, 54.3, 51.2, 42.9, 39.6, 38.0, 35.8, 35.2, 33.5, 32.7, 31.4, 28.4, 27.9, 26.7, 26.3, 26.1, 23.2, 20.3, 11.6, 11.3.

**19-((3α-(Methoxymethoxy)-5α-androstan-17β-yl)oxy)-3,6,9,12,15-pentaoxanonadecan-1-al (8).** To a stirred solution of oxalyl chloride (0.25 mL, 3mmol) in $CH_2Cl_2$ (10 mL) was added DMSO (0.32 mL, 4.5 mmol) in $CH_2Cl_2$ (4 mL) at -78 °C. After 10 min, steroid **7** (101 mg, 0.161 mmol) in $CH_2Cl_2$ (8 mL) was added and stirred at -78 °C for 2 h. $Et_3N$ (0.84 mL, 6 mmol) was added at -78 °C and stirred for 30 min. The reaction was warmed to 23 °C for 30 min. Water was added and the product was extracted into $CH_2Cl_2$ (150 mL x 2). The combined extracts were dried over $Na_2SO_4$, filtered, the solvent removed and the residue was purified by flash column chromatography (silica gel, eluted with 50% EtOAc in hexanes) to give steroid **8** (98 mg, 97%): $^1$H NMR (400 MHz, $CDCl_3$) δ 9.72 (s, 1H), 4.65-4.64 (m, 2H), 4.16 (s, 1H),3.82 (s, 1H), 3.36 (s, 3H), 3.72-3.36 (m, 21H), 2.20-0.86 (m, 27H), 0.78 (s, 3H), 0.72 (s, 3H); $^{13}$C NMR (100 MHz, $CDCl_3$) δ 200.9, 94.4, 89.0, 76.7, 71.5, 71.2, 71.1, 70.6, 70.5, 70.47 (3 x C), 70.4, 69.9, 69.7, 55.0, 54.3, 51.2, 42.9, 39.7, 38.0, 35.8, 35.2, 33.5, 32.7, 31.4, 28.4, 28.0, 26.7, 26.4, 26.2, 23.2, 20.3, 11.6, 11.3.

**20-((3α-(Methoxymethoxy)-5α-androstan-17β-yl)oxy)-4,7,10,13,16-pentaoxaicos-1-yne (9).** To a stirred solution of steroid **8** (98 mg, 0.157 mmol) in MeOH/ THF (3 mL/3 mL) was added dimethyl-1-diazo-2-oxopropyl-phosphonate (0.23 mL, 1.5 mmol) and $K_2CO_3$ (414 mg, 3 mmol) at 23 °C. After 16 h, water (20 mL) was added and the product was extracted into EtOAc (100 mL x 2). The combined extracts were dried over $Na_2SO_4$, filtered, the solvent removed and the residue purified by flash column chromatography (silica gel, eluted with 40% EtOAc in hexanes) to give steroid **9** (68 mg, 70%): $^1$H NMR (400 MHz, $CDCl_3$) δ 4.67-4.64 (q, $J$ = 6.6 Hz, 2H), 4.20 (s, 2H), 3.82 (s, 1H), 3.68-3.23 (m, 20H), 3.36 (s, 3H), 2.19-2.15 (m, 2H), 2.43-0.75 (m, 26H), 0.78 (s, 3H), 0.72 (s, 3H); $^{13}$C NMR (100 MHz, $CDCl_3$) δ 94.4, 89.1, 79.6, 74.5, 71.5, 71.2, 70.5 (3 x C), 70.4, 70.3, 69.9, 69.7, 69.0, 58.3, 55.1, 54.4, 51.3, 43.0, 39.7, 38.1, 35.8, 35.2, 33.6, 32.8, 31.5, 29.6, 28.4, 28.0, 26.7, 26.4, 26.2, 23.2, 20.4, 11.7, 11.3.

**17β-((4,7,10,13,16-Pentaoxaicos-1-yn-20-yl)oxy)-5α-androstan-3α-ol (10, MQ311).** To a stirred solution of compound **9** (68 mg, 0.11 mmol) in MeOH (6 mL) was added acetyl chloride (2 mL) at 23 °C. After 2 h, water was added and the product was extracted into $CH_2Cl_2$ (60 mL x 2). The combined extracts were washed with aqueous $NaHCO_3$ (50 mL), dried over $Na_2SO_4$, filtered, the solvent removed and the residue purified by flash column chromatography (silica gel, eluted with 40% EtOAc in hexanes) to give **MQ311** (**10**, 32 mg, 51%): $^1$H NMR (400 MHz, $CDCl_3$) δ 4.20 (s, 2H), 4.03 (s, 1H), 3.71-3.24 (m, 23H), 2.44 (s, 1H), 1.97-0.75 (m, 25H), 0.77 (s, 3H), 0.72 (s, 3H); $^{13}$C NMR (100 MHz, $CDCl_3$) δ 89.1, 79.6, 74.5, 71.2, 70.5, 70.5 (4 x C), 70.3, 70.0, 69.7, 69.0, 66.5, 58.3, 54.4, 51.2, 43.0, 39.1, 38.1, 36.1, 35.8, 35.3, 32.1, 31.5, 28.9, 28.4, 28.0, 26.7, 26.4, 23.3, 20.4, 11.7, 11.1.

**Synthesis of MQ302**

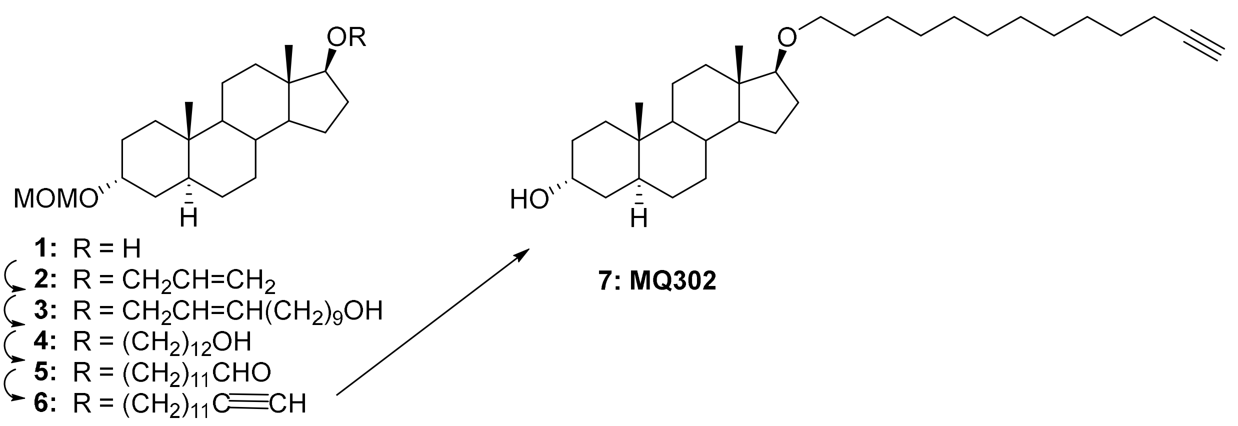


**3α-(Methoxymethoxy)-17β-(2-propenyloxy)-5α-androstane (2).** 3α-(Methoxymethoxy)-5α-androstan-17β-ol (**1**) was prepared according to the literature (Chintala, S.M. et al. *Br. J. Pharmacol.* 2024**, 181**, 4229-4244). To a stirred solution of steroid **1** (1.8 g, 5,4 mmol) in THF (90 mL) was added NaH (2.17 g, 54.2 mmol, 60% in mineral oil) and allyl bromide (6.95 mL, 81 mmol) at 23 °C. The reaction was refluxed for 16 h, cooled to 23 °C, water was slowly added and the product was extracted into EtOAc (200 mL x 2). The combined extracts were washed with brine (80 mL), dried over anhydrous $Na_2SO_4$, filtered and the solvent removed. The residue was purified on a CombiFlash chromatography apparatus equipped with a silica gel cartridge eluted with 0-10% EtOAc in hexanes to give steroid **2** (1.9 g, 93%): $^{1}$H NMR (400 MHz, $CDCl_3$) δ 5.95-5.85 (m, 1H), 528 (dd, *J* = 1.6 Hz, 7.5, 1H), 5.14 (dd, *J* = 1.9 Hz, 10.5, 1H), 4.68 (q, *J* = 7.0 Hz, 2H), 400-3.98 (m, 1H), 384-3.82 (m, 1H), 3.37 (s, 3H), 3.36-3.33 (m, 1H), 2.05-0.66 (m, 22H), 0.80 (s, 3H), 0.77 (s, 3H); $^{13}$C NMR (100 MHz, $CDCl_3$) δ 135.7, 115.9, 94.5, 88.5, 71.6, 70.8, 55.1, 54.4, 51.3, 43.0, 39.7, 38.0, 35.9, 35.3, 33.6, 32.8, 31.5, 28.5, 28.0, 26.3, 23.3, 20.4, 11.8, 11.4.

**((3α-(Methoxymethoxy)-5α-androstane-17β-yl)oxy)dodec-10-en-1-ol (3).** To a stirred solution of steroid **2** (460 mg, 1.22 mmol) in THF (6 mL) was added 10-undecen-1-ol and Hoveyda-Grubbs catalysts M2001 (19.3 mg, 0.031 mmol) in THF (2 mL) at 23 °C. After 16 h, THF was removed and the residue was purified by flash column chromatography (silica gel, eluted with 10-15% EtOAc in hexanes) to give steroid **3** (171 mg, 33%): $^{1}$H NMR (400 MHz, $CDCl_3$) δ 5.65-5.36 (m, 2H), 4.72-4.62 (m, 2H), 3.93-3.91 (m, 1H), 3.81 (s, 1H), 3.62 (t, *J* = 6.6 Hz, 2H), 3.35 (s, 3H), 3.33-3.30 (m, 1H), 2.03-0.70 (m, 42H), 0.78 (s, 3H), 0.75 (s, 3H); $^{13}$C NMR (100 MHz, $CDCl_3$) δ 133.7, 127.0, 94.4, 88.1, 71.6, 62.8, 55.0, 54.4, 51.3, 42.9, 39.7, 38.0, 35.8, 35.2, 33.5, 32.8, 32.7, 32.2, 31.5, 29.5, 29.3, 29.3, 29.0, 28.4, 28.0, 26.2, 25.7, 23.3, 20.4, 20.4, 11.7, 11.3.

**((3α-(Methoxymethoxy)-5α-androstane-17β-yl)oxy)dodecan-1-ol (4).** To a stirred solution of steroid **3** (171 mg, 0.33 mmol) in EtOAc (45 mL) was added Pd/C (100 mg) at 23 °C in a Parr hydrogenation flask. The flask was evacuated and charged with $H_2$ three times. The hydrogenation was carried out overnight at 55 psi $H_2$. The reaction mixture was filtered through Celite and washed with EtOAc (100 mL). The solvent was removed and the residue was purified by flash column chromatography (silica gel, eluted with 20% EtOAc in hexanes) to give steroid **4** (150 mg, 88%): $^{1}$H NMR (400 MHz, $CDCl_3$) δ 4.63-4.61 (m, 2H), 3.78 (s, 1H), 3.58 (t, *J* = 6.6 Hz, 2H), 3.32 (s, 3H), 3.41-3.33 (m, 1H), 3.24 (t, *J* = 8.2 Hz, 1H), 2.08-0.83 (m, 44H), 0.75 (s, 3H), 0.70 (s, 3H); $^{13}$C NMR (100 MHz, $CDCl_3$) δ 94.4, 69.1, 72.0, 70.2, 62.8, 55.1, 54.4, 51.4, 43.0, 39.7, 38.2, 35.9, 35.3, 33.6, 32.8, 32.7, 31.5, 30.2, 29.6 (3 x C), 29.5, 29.4, 28.5, 28.1, 26.3, 26.2, 25.8, 23.3, 20.4, 11.7, 11.4.

**((3α-(Methoxymethoxy)-5α-androstane-17β-yl)oxy)dodecan-1-al (5).** To a stirred solution of oxalyl chloride (0.17 mL, 2 mmol) in $CH_2Cl_2$ (10 mL) was added DMSO (0.17 mL, 2.4 mmol) in $CH_2Cl_2$ (2 mL) at -78 °C. After 10 min, steroid **4** (150 mg, 0.3 mmol) in $CH_2Cl_2$ (6 mL) was added.

After 2 h, $Et_3N$ (0.56 mL, 4 mmol) was added and stirring continued for 1 h. Water (30 mL) was added and the product was extracted into $CH_2Cl_2$ (100 mL x 2). The combined extracts were dried over anhydrous $Na_2SO_4$, filtered, the solvent removed and the residue purified by flash column chromatography (silica gel, eluted with 15% EtOAc in hexanes) to give steroid **5** (142 mg, 95%): $^1$H NMR (400 MHz, $CDCl_3$) δ 9.74 (s, 1H), 4.65-4.61 (m, 2H), 3.80 (s, 1H), 3.43-3.35 (m, 2H), 3.34 (s, 3H), 3.26 (t, $J$ = 8.2 Hz, 1H), 2.41-2.37 (m, 2H), 1.96-0.70 (m, 40H), 0.77 (s, 3H), 0.72 (s, 3H); $^{13}$C NMR (100 MHz, $CDCl_3$) δ 202.7, 94.4, 89.0, 71.5, 70.1, 55.0, 54.4, 51.3, 43.8, 43.0, 39.7, 38.1, 35.8, 35.3, 33.6, 32.8, 31.5, 30.1, 29.5, 29.4, 29.4, 29.3, 29.3, 29.1, 28.4, 28.1, 26.2, 26.1, 23.2, 22.0, 20.4, 11.6, 11.3.

**3α-(Methoxymethoxy)-17β-(tridec-12-yn-1-yloxy)-5α-androstane (6).** To a stirred solution of steroid **5** (142 mg, 0.27 mmol) in MeOH/THF (3 mL/3 mL) was added dimethyl-1-diazo-2-oxopropyl-phosphonate (0.23 mL, 1.5 mmol) and $K_2CO_3$ (414 mg, 3 mmol) at 23 °C. After 3 days, water (20 mL) was added and the product was extracted into EtOAc (100 mL x 2). The combined extracts were dried over anhydrous $Na_2SO_4$, filtered, the solvent removed and the residue purified by flash column chromatography (silica gel, eluted with 20% EtOAc in hexanes) to give steroid **6** (110 mg, 78%): $^1$H NMR (400 MHz, $CDCl_3$) δ 4.67-4.64 (m, 2H), 3.82 (s, 1H), 3.44-3.37 (m, 2H), 3.36 (s, 3H), 3.28 (t, $J$ = 8.6 Hz, 1H), 2.19-2.15 (m, 2H), 1.97-0.70 (m, 41H), 0.77 (s, 3H), 0.72 (s, 3H); $^{13}$C NMR (100 MHz, $CDCl_3$) δ 94.4, 89.1, 84.7, 71.6, 70.1, 68.0, 55.1, 54.4, 51.4, 43.0, 39.7, 38.2, 35.9, 35.3, 33.6, 32.8, 31.5, 30.2, 29.5, 29.5, 29.4 (2 x C), 29.1, 28.7, 28.5, 28.5, 28.1, 26.3, 26.2, 23.3, 20.4, 18.3, 11.7, 11.4.

**17β-(Tridec-12-yn-1-yloxy)-5α-androstan-3a-ol (7, MQ302).** To a stirred solution of steroid **6** (110 mg, 0.214 mmol) in THF (4 mL) was added 6 N HCl (4 mL) at 23°C. After 4 h, the product was extracted into EtOAc (100 mL x 2). The combined extracts were washed with aqueous $NaHCO_3$ (50 mL), dried over anhydrous $Na_2SO_4$, filtered, the solvent removed and the residue purified by flash column chromatography (silica gel, eluted with 25% EtOAc in hexanes) to give steroid **MQ302** (**7**, 22 mg, 22%): $^1$H NMR (400 MHz, $CDCl_3$) δ 4.05-4.04 (m, 1H), 3.48-3.36 (m, 2H), 3.30 (t, $J$ = 8.5 Hz, 1H), 2.20-2.16 (m, 2H), 2.00-0.71 (m, 42H), 0.79 (s, 3H), 0.75 (s, 3H); $^{13}$C NMR (100 MHz, $CDCl_3$) δ 89.1, 84.8, 70.2, 68.0, 66.5, 54.5, 51.4, 43.0, 39.2, 38.2, 36.2, 35.9,

35.3, 32.2, 31.6, 30.2, 29.6, 29.5, 29.5 (2 x C), 29.1, 29.0, 28.7, 28.5, 28.4, 28.1, 26.2, 23.3, 20.5, 18.4, 11.7, 11.2.

**Synthesis of MQ303**

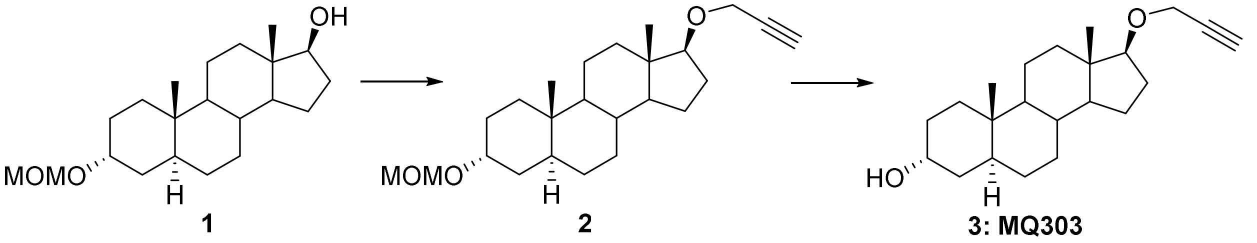


**3α-(Methoxymethoxy)-17β-(2-propynyloxy)-5α-androstane (2).** 3α-(Methoxymethoxy)-5α-androstan-17β-ol (**1**) was prepared according to the literature (Chintala, S.M. et al., *Br. J. Pharmacol.* 2024,**181**, 4229-4244). To a stirred solution of steroid **1** (250 mg, 0.74 mmol) in THF (30 mL) was added NaH (600 mg, 15 mmol, 60% in mineral oil) and propargyl bromide (80% in toluene, 3 mL) at 23 °C. The mixture was refluxed for 16 h. After cooling to 23 °C, water was slowly added and the product was extracted into EtOAc (150 mL x 2). The combined extracts were dried over anhydrous $Na_2SO_4$, filtered, solvent removed and the residue purified by flash column chromatography (silica gel, eluted with 10% EtOAc in hexanes) to give steroid **2** (217 mg, 79%): $^1$H NMR (400 MHz, $CDCl_3$) δ 4.64 ( q, *J* = 7.0 Hz, 2H), 4.11 (t, *J* = 2.0 Hz, 2H), 3.80-3.79 (m, 1H), 3.51 (t, *J* = 8.2 Hz, 1H), 3.33 (s, 3H), 2.38 (t, *J* = 2.4 Hz, 1H), 2.20-0.69 (m, 22H), 0.77 (s, 3H), 0.73 (s, 3H); $^{13}$C NMR (100 MHz, $CDCl_3$) δ 94.4, 88.2, 80.5, 73.6, 71.4, 57.0, 55.0, 54.3, 51.1, 42.8, 39.6, 37.6, 35.8, 35.2, 33.5, 32.7, 31.4, 28.4, 27.5, 26.2, 23.2, 20.3, 11.6, 11.3.

**17β-(2-Propynyloxy)-5α-androstan-3α-ol (3, MQ303).** To a stirred solution of steroid **2** in MeOH (20 mL) was slowly added acetyl chloride (2 mL) at 23 °C. After 2 h, water (30 mL) was added and the product was extracted into $CH_2Cl_2$ (100 mL x 2). The combined extracts were washed with aqueous $NaHCO_3$ (50 mL), dried over anhydrous $Na_2SO_4$, filtered and the solvent removed. The residue was purified by flash column chromatography (silica gel, eluted with 20% EtOAc in hexanes) to give **MQ302** (**3**, 162 mg, 85%): $^{1}$H NMR (400 MHz, $CDCl_3$) δ 4.11 (m, 4.14-4.12, 2H), 4.02-4.01 (m, 1H), 3.53 (t, $J$ = 8.2 Hz, 1H), 2.39 (t, $J$ = 2.3 Hz, 1H), 2.03-0.70 (m, 23H), 0.77 (s, 3H), 0.74 (s, 3H); $^{13}$C NMR (100 MHz, $CD_3OD$) δ 88.2, 80.5, 73.6, 66.3, 57.1, 54.3, 51.1, 42.8, 39.0, 37.6, 36.0, 35.8, 35.2, 32.1, 31.5, 28.9, 28.3, 27.5, 23.2, 20.3, 11.6, 11.1.

**Synthesis of YX46 and MQ342**

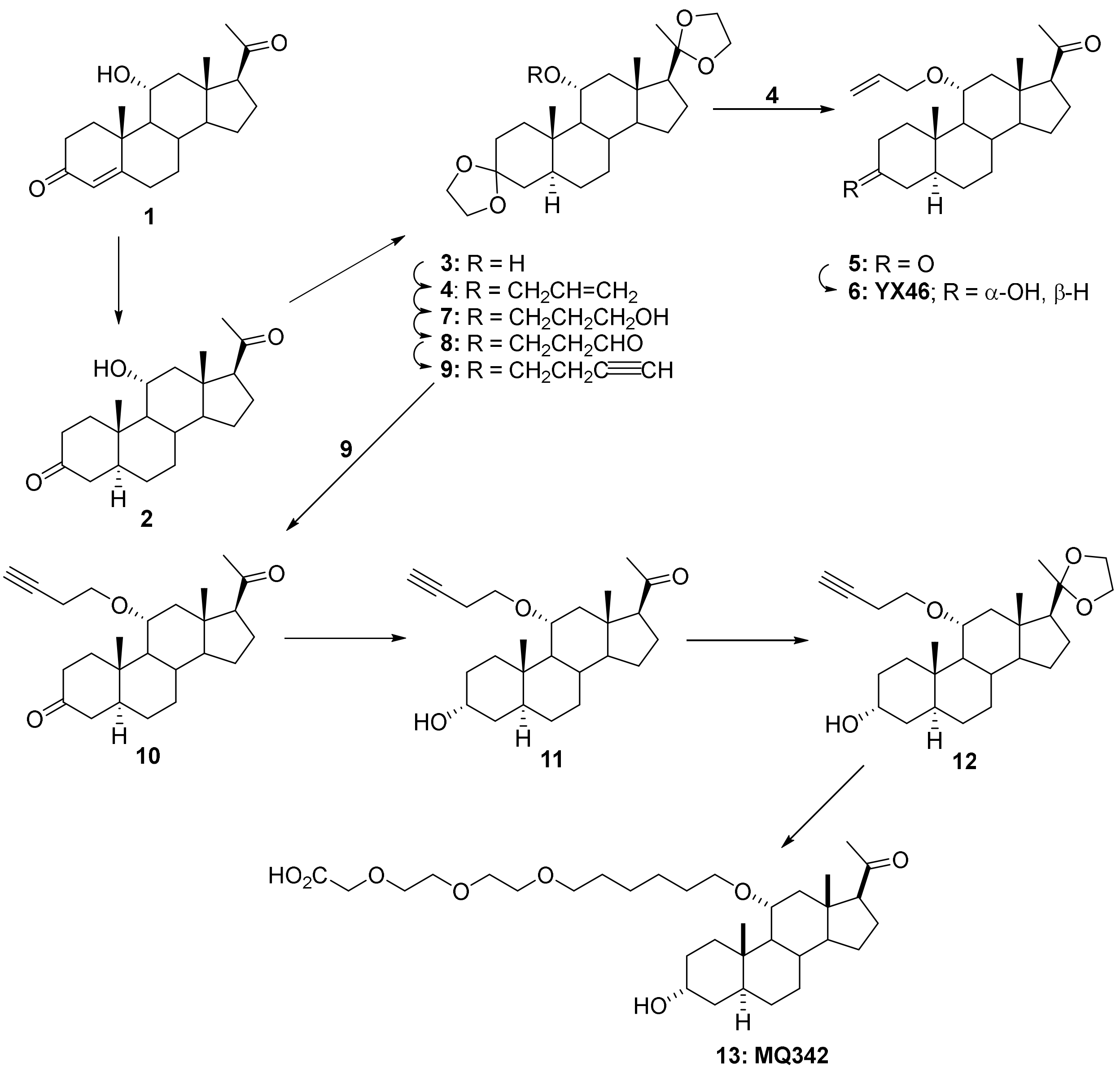

**11α-Hydroxy-5α-pregnane-3,20-dione (2).** A mixture of 17α-hydroxyprogesterone (**1**, 1.0 g, 3.0 mmol) dissolved in THF (20 mL) and 5% palladium on $CaCO_3$ (200 mg) was placed in a Parr hydrogenation flask and hydrogenation was carried at 50 psi for 16 h at 23 °C. The contents of the flask were filtered through celite and the filter cake was washed with $CH_2Cl_2$. Solvent was removed from the filtrate and the residue was purified by flash column chromatography (silica gel, eluted with 50% EtOAc in hexanes) to give steroid **2** (470 mg, 47%): $^1$H NMR (400 MHz, $CDCl_3$) δ 3.86-3.85 (m, 1H), 2.73-2.69 (m, 1H), 2.47-0.79 (m, 21H), 2.01 (s, 3H), 1.02 (s, 3H), 0.53 (s, 3H); $^{13}$C NMR (100 MHz, $CDCl_3$) δ 212.2, 209.0, 68.3, 63.0, 59.1, 55.2, 50.0, 47.2, 44.8, 43.8, 39.8, 38.1, 36.9, 34.1, 31.2, 31.1, 29.0, 24.0, 22.5, 14.1, 11.4.

**17β-(2-Methyl-1,3-dioxolan-2-yl)-spiro[5α-androstan-3,2'-[1,3]dioxolan]-11α-ol (3).** To a stirred solution of steroid **2** (470 mg, 1.42 mmol) in benzene (100 mL) was added ethylene glycol (2.0 mL) and PTSA (100 mg) at 23 °C. The reaction was refluxed in a flask equipped with a Dean-Stark apparatus for 16 h. The flask was cooled to 23 °C, aqueous $NaHCO_3$ was added and the product was extracted into EtOAc (100 mL × 2). The combined extracts were washed with brine (100 mL × 5), dried over anhydrous $Na_2SO_4$, filtered and the solvent was removed. The residue was dried under high vacuum to give steroid **3** (550 mg, 92%): $^1$H NMR (400 MHz, $CDCl_3$) δ 3.92-3.78 (m, 9H), 2.23-0.53 (m, 22H), 1.20 (s, 3H), 0.86 (s, 3H), 0.68 (s, 3H); $^{13}$C NMR (100 MHz, $CDCl_3$) δ 111.4, 108.8, 68.7, 64.9, 63.8, 63.8, 63.0, 60.0, 57.9, 55.2, 51.1, 43.7, 42.2, 38.2, 37.4, 36.9, 33.9, 31.6, 31.0, 28.9, 24.3, 23.6, 22.7, 13.9, 11.6.

**17β-(2-Methyl-1,3-dioxolan-2-yl)-11α-(2-propenyloxy)spiro[5α-androstane-3,2'-[1,3]dioxolane] (4).** To a stirred solution of steroid **3** (550 mg, 1.31 mmol) in THF (30 mL) was added KH (400 mg, 10 mmol) at 23 °C. The mixture was refluxed for 30 min. Allyl bromide (2 mL) was added and the reaction was stirred for 16 h. The reaction was cooled to 23 °C and MeOH (2 mL) was added. After 10 min, water (20 mL) was added and the product was extracted into EtOAc (20 mL × 3). The combined extracts were dried over anhydrous $Na_2SO_4$, filtered and the solvent was removed. The residue was purified by flash column chromatography (silica gel, eluted with 10% EtOAc in hexanes) to give steroid **4** (390 mg, 65%): $^1$H NMR (400 MHz, $CDCl_3$) δ 5.86-5.82 (m, 1H), 5.22-5.17 (m, 1H), 5.07-5.04 (m, 1H), 4.05-3.75 (m, 10H), 3.52-3.48 (m, 1H), 2.47-2.43 (m, 1H), 2.06-2.02 (m, 1H), 1.81-0.85 (m, 17H), 1.26 (s, 3H), 0.87 (s, 3H), 0.70 (s, 3H); $^{13}$C

NMR (100 MHz, $CDCl_3$) δ 135.3, 115.9, 111.5, 108.8, 76.2, 68.7, 65.0, 64.0, 63.9, 63.1, 57.9, 57.3, 55.1, 45.0, 43.7, 41.9, 38.4, 37.1, 37.0, 34.2, 31.8, 31.2, 29.0, 24.3, 23.8, 22.9, 13.9, 12.0.

**11α-(2-Propenyloxy)-5α-pregnan-3,20-dione (5).** To a stirred solution of steroid **4** (390 mg, 0.85 mmol) in acetone (20 mL) was added PTSA (100 mg in 5 mL acetone) at 23 °C. The reaction was stirred for 16 h. Solid $NaHCO_3$ was added and after 10 min the acetone was removed. Water (20 mL) was added to the residue and the product was extracted into EtOAc (100 mL). The EtOAc was dried over anhydrous $Na_2SO_4$, filtered and the solvent removed. The residue was purified by flash column chromatography (silica gel, eluted with 10% EtOAc in hexanes) to give steroid **5** (300 mg, 94%): $^1$H NMR (400 MHz, $CDCl_3$) δ 5.89-5.80 (m, 1H), 5.23-5.19 (m, 1H), 5.11-5.08 (m, 1H), 4.09-4.05 (m, 1H), 3.84-3.80 (m, 1H), 3.59-3.53 (m, 1H), 2.53-0.90 (m, 21H), 2.09 (s, 3H), 1.06 (s, 3H), 0.59 (s, 3H); $^{13}$C NMR (100 MHz, $CDCl_3$) δ 211.8, 208.9, 134.7, 116.5, 76.1, 69.0, 63.1, 57.3, 55.2, 47.1, 45.0, 44.2, 43.5, 39.7, 38.3, 37.1, 34.5, 31.5, 31.3, 29.2, 24.3, 22.9, 14.2, 12.0.

**3α-(Hydroxy)-11α-(2-propenyloxy)-5α-pregnan-20-one (6, YX46).** To a solution of steroid **5** (300 mg, 0.8 mmol) in THF (50 mL) was added dropwise K-seletride (1M in THF, 1.0 mL, 1.0 mmol) over 20 min at -78 °C. The reaction was stirred for 2 h, acetone (3 mL) was added and stirring was continued for 15 min at -78 °C. Aqueous $NH_4Cl$ was added and after 1 h at 23 °C the product was extracted into EtOAc (50 mL × 3). The combined extracts were dried over anhydrous $Na_2SO_4$, filtered and the solvent removed. The residue was purified by flash column chromatography (silica gel, eluted with 25% EtOAc and 25% $CH_2Cl_2$ in hexanes) to give steroid **YX46** (**6**, 220 mg, 73%): $^1$H NMR (400 MHz, $CDCl_3$) δ 5.86-5.79 (m, 1H), 5.20-5.16 (m, 1H), 5.06-5.03 (m, 1H), 4.03-3.99 (m, 1H), 3.91 (s, 1H), 3.80-3.76 (m, 1H), 3.50-3.44 (m, 1H), 2.50-2.37 (m, 2H), 2.06-0.88 (m, 20H), 2.06 (s, 3H), 0.81 (s, 3H), 0.52 (s, 3H); $^{13}$C NMR (100 MHz, $CDCl_3$) δ 209.2, 135.0, 116.2, 76.2, 68.9, 65.8, 63.2, 57.4, 55.3, 44.4, 43.6, 38.7, 37.7, 36.1, 34.5, 33.3, 31.9, 31.3, 29.2, 28.8, 24.2, 22.8, 14.1, 11.7.

**17β-(2-Methyl-1,3-dioxolan-2-yl)-11α-(1-hydroxy-3-propoxy)spiro[5α-androstane-3,2'-[1,3]dioxolane] (7).** To a solution of steroid **4** (1.13 g, 2.45 mmol) in THF (30 mL) was added 9-BBN (0.5 M in THF, 10 mL, 5 mmol) at 23 °C. After 16 h, 3 N NaOH (30 mL, 90 mmol) and $H_2O_2$ (10 mL) were added and stirred at 23 °C for 1 h. The product was extracted into EtAOc (150 mL x 3). The combined extracts were washed with brine (100 mL x 3), dried over anhydrous $Na_2SO_4$, filtered, solvent removed and the residue was purified by flash column chromatography (silica gel, eluted with 25-50% EtOAc in hexanes) to give steroid **7** (950 mg, 81%): $^1$H NMR (400 MHz, $CDCl_3$) δ 3.99-3.85 (m, 8H), 3.72-3.65 (m, 3H), 3.47-3.40 (m 2H), 2.52-2.48 (m, 2H), 2.03-0.92 (m, 22H), 1.28 (s, 3H), 0.89 (s, 3H), 0.71 (s, 3H); $^{13}$C NMR (100 MHz, $CDCl_3$) δ 111.6, 108.8, 76.9, 67.0, 65.0, 64.03, 64.01, 63.1, 62.2, 57.8, 57.5, 55.0, 45.1, 43.7, 41.9, 38.5, 37.2, 37.1, 34.2, 32.4, 31.8, 31.3, 29.0, 24.4, 23.8, 23.0, 14.0, 12.1.

**17β-(2-Methyl-1,3-dioxolan-2-yl)-11α-(1-oxo-3-propoxy)spiro[5α-androstane-3,2'-[1,3]dioxolane] (8).** To a stirred solution of oxalyl chloride (1 mL, 12 mmol) in $CH_2Cl_2$ (25 mL) was added DMSO (1 mL, 14 mmol) in $CH_2Cl_2$ (3 mL) at -78 °C and stirred for 15 min. Steroid **7** (950 mg, 1.98 mmol) in $CH_2Cl_2$ (10 mL) was added and stirred for 1 h at -78 °C. $Et_3N$ (2.8 mL, 20 mmol) was added and stirred for 30 min. The reaction was warmed to 23 °C and water was added. The product was extracted into $CH_2Cl_2$ (100 mL x 3). The combined extracts were dried over anhydrous $Na_2SO_4$, filtered, solvent removed and the residue was purified by flash column chromatography (silica gel, eluted with 25-40% EtOAc in hexanes) to give steroid **8** (760 mg, 80%): $^1$H NMR (400 MHz, $CDCl_3$) δ 9.75 (s, 1H), 4.01-3.85 (m, 9H), 3.59-3.45 (m, 2H), 2.61-2.48 (m, 3H), 1.91-0.93 (m, 20H), 1.29 (s, 3H), 0.89 (s, 3H), 0.72 (s, 3H); $^{13}$C NMR (100 MHz, $CDCl_3$) δ 201.8, 111.6, 108.8, 65.0, 64.1, 64.0, 63.1, 61.1, 57.8, 57.3, 55.0, 44.9, 44.3, 43.7, 41.9, 38.4, 37.1, 37.0, 34.2, 31.8, 31.2, 29.0, 24.4, 23.8, 23.1, 14.0, 12.1.

**17β-(2-Methyl-1,3-dioxolan-2-yl)-11α-(but-3-yn-1-yloxy)spiro[5α-androstane-3,2'-[1,3]dioxolane (9).** To a stirred solution of steroid **8** (740 mg, 1.55 mmol) in THF/MeOH (10 mL/10 mL) was added dimethyl-1-diazo-oxopropyl phosphonate (0.62 mL, 4 mmol) and $K_2CO_3$ (1.1 g, 8 mmol, fine powder) at 23 °C. After 16 h, water was added and the product was extracted into EtOAc (100 mL x 3). The combined extracts were dried over anhydrous $Na_2SO_4$, filtered, solvent removed and the residue was purified by flash column chromatography (silica gel, eluted with 25-35% EtOAc in hexanes) to give steroid **9** (580 mg, 79%): $^1$H NMR (400 MHz, $CDCl_3$) δ 3.98-3.82 (m, 8H), 3.66-3.61 (m, 1H), 3.50-3.44 (m, 1H), 3.38-3.32 (m, 1H), 2.47-2.34 (m, 3H), 2.10-0.91 (m, 21H), 1.27 (s, 3H), 0.88 (s, 3H), 0.70 (s, 3H); $^{13}$C NMR (100 MHz, $CDCl_3$) δ 111.6, 108.9, 81.7, 76.7, 69.1, 65.8, 65.0, 64.0, 63.9, 63.1, 57.9, 57.3, 55.1, 45.0, 43.7, 41.9, 38.4, 37.02, 37.01, 34.2, 31.8, 31.2, 29.0, 24.4, 23.8, 23.0, 20.2, 14.0, 12.0.

**11α-(But-3-yn-1-yloxy)-5α-pregnane-3,20-dione (10).** To a stirred solution of steroid **9** (580 mg, 1.23 mmol) in acetone (40 mL) was added PTSA (100 mg) at 23 °C. After 16 h, solid $NaHCO_3$ (300 mg) was added and stirred for 15 min. The solvent was removed and the residue was purified by flash column chromatography (silica gel, eluted with 30-40% EtOAc in hexanes) to give steroid **10** (450 mg, 95%): $^1$H NMR (400 MHz, $CDCl_3$) δ 3.83 (s, 1H), 3.65-3.31 (m, 3H), 2.51-0.84 (m, 23H), 2.05 (s, 3H), 1.02 (s, 3H), 0.55 (s, 3H); $^{13}$C NMR (100 MHz, $CDCl_3$) δ 211.5, 208.7, 81.5, 76.5, 69.2, 66.0, 62.9, 57.1, 55.1, 47.0, 44.9, 43.9, 43.3, 39.5, 38.1, 36.9, 34.3, 31.4, 31.2, 29.0, 24.2, 22.8, 20.1, 14.1, 11.8.

**11α-(But-3-yn-1-yloxy)-3α-hydroxy-5α-pregnane-20-one (11).** To a stirred solution of steroid **10** (450 mg, 1.17 mmol) in THF (60 mL) was added K-selectride (1.0 M in THF, 2 ml, 2 mmol) at -78 °C. After 2 h, 1 mL of acetone was added and stirred for 15 min. Aqueous $NH_4Cl$ was added and stirring was continued at 23 °C for 2 h. The product was extracted into EtOAc (100 mL x 3). The combined extracts were dried over anhydrous $Na_2SO_4$, filtered, solvent removed and the residue was purified by flash column chromatography (silica gel, eluted with 45% EtOAc in hexanes) to give steroid **11** (330 mg, 73%): $^1$H NMR (400 MHz, $CDCl_3$) δ 3.99 (s, 1H), 3.67-3.40 (m, 3H), 2.56-2.41 (m, 4H), 2.12 (s, 3H), 2.08-0.92 (m, 21H), 0.87 (s, 3H), 0.59 (s, 3H); $^{13}$C NMR (100 MHz, $CDCl_3$) δ 209.2, 81.6, 76.7, 69.2, 66.2, 66.1, 63.3, 57.5, 55.4, 44.4, 43.7, 38.9, 37.8, 36.3, 34.6, 33.3, 32.0, 31.5, 29.3, 28.9, 24.4, 23.0, 20.3, 14.3, 11.8.

**11α-(But-3-yn-1-yloxy)-17β-(2-methyl-1,3-dioxolan-2-yl)-5α-androstan-3α-ol (12).** To a stirred solution of steroid **11** (330 mg, 0.85 mmol) in benzene (150 mL) was added ethylene glycol (1 mL) and PTSA (50 mg) at 23 °C. The reaction was refluxed in a flask equipped with a Dean-Stark apparatus under $N_2$ for 16 h. After cooling, solid $NaHCO_3$ (300 mg) was added and stirred for 10 min. Aqueous $NaHCO_3$ (100 mL) was added and the product was extracted into EtAOc (300 mL). The extract was washed with brine (100 mL x 3), dried over anhydrous $Na_2SO_4$, filtered, solvent removed and the residue was purified by flash column chromatography (silica gel, eluted with 45% EtOAc in hexanes) to give steroid **12** (280 mg, 76%): $^1$H NMR (400 MHz, $CDCl_3$) δ 3.99-3.83 (m, 4H), 3.67-3.35 (m, 3H), 2.47-2.35 (m, 3H), 2.02-0.90 (m, 23H), 1.27 (s, 3H), 0.86 (s, 3H), 0.71 (s, 3H); $^{13}$C NMR (100 MHz, $CDCl_3$) δ 111.6, 81.7, 76.8, 69.1, 66.2, 65.8, 65.0, 63.1, 57.9, 57.5, 55.1, 45.0, 41.9, 38.9, 37.7, 36.3, 34.2, 33.3, 32.0, 29.2, 29.0, 24.4, 23.7, 23.0, 20.2, 14.0, 11.8.

**2-(2-(2-((6-((3α-Hydroxy-5α-pregnan-20-on-11α-yl)oxy)hexyl)oxy)ethoxy)ethoxy)acetic acid (12, MQ342).** To a mixture of steroid **12** (70 mg, 0.21 mmol), methyl 3-(2-(2-(2-iodoethoxy)ethoxy)ethoxy)propanoate (96 mg, 0.2 mmol), bis[(2-dimethylamino)phenyl]amine nickel(ii) chloride (7.2 mg, 0.15 mmol), CuI (2.5 mg, 0.012 mmol), $Cs_2CO_3$ (163 mg, 0.5 mmol) was added anhydrous dioxane (25 mL) at room temperature. The stirred reaction was heated to 140 °C for 16 h. After cooling, water was added, and the crude product was extracted into EtOAc. The crude alkyne product was a mixture of the 17-ketone and 17-ketal which was not separable by column chromatography.

To a solution of the crude alkyne product in EtOAc (60 mL) was added Pd/C (300 mg) in a Parr hydrogenation flask at 23 °C. The flask was evacuated and charged with $H_2$ three times. Hydrogenation was carried out at 55 psi overnight. The product was filtered through Celite and washed with EtOAc (100 mL). LC-MS (ESI) m/z: calcd for C33H56O8Na [M +Na]+, 603.4; found, 603.4 (17-ketone)and calcd for C35H60O9Na [M +Na]+, 647.4; found, 647.5 (17-ketal).

The EtOAc was removed and the 17-ketone/17-ketal mixture was dissolved in acetone (40 mL), PTSA (100 mg) was added and the reaction was stirred at 23 °C for 16 h. Solid $NaHCO_3$ (300 mg) was added and stirred for 15 min. The solvent was removed and the residue was purified by flash column chromatography (silica gel, eluted with 10-25% MeOH in dichloromethane) to **MQ342** (**13**, 18 mg, 15%, 3 steps): $^{1}$H NMR (400 MHz, $CDCl_3$) δ 4.00-3.98 (m, 3H), 3.71-3.20 (m, 13H), 3.0 (br, s, 1H), 2.49-2.45 (m, 2H), 2.19-2.10 (m, 1H), 2.15 (s, 3H), 1.85-1.00 (m, 27H), 0.88 (s, 3H), 061 (s, 3H); $^{13}$C NMR (100 MHz, $CDCl_3$) δ 209.4, 174.1, 77.2, 76.4, 71.4, 67.9, 66.0, 63.4, 57.6, 55.5, 44.5, 43.8 (2 x C), 39.0, 37.8 (2 x C), 36.3, 34.7, 33.4, 32.1, 30.4, 29.7, 29.4, 29.3, 29.0, 26.3, 26.0, 24.4, 23.0, 14.3 (2 x C), 11.9 (2 x C).

**Synthesis of YX32 andYX62**

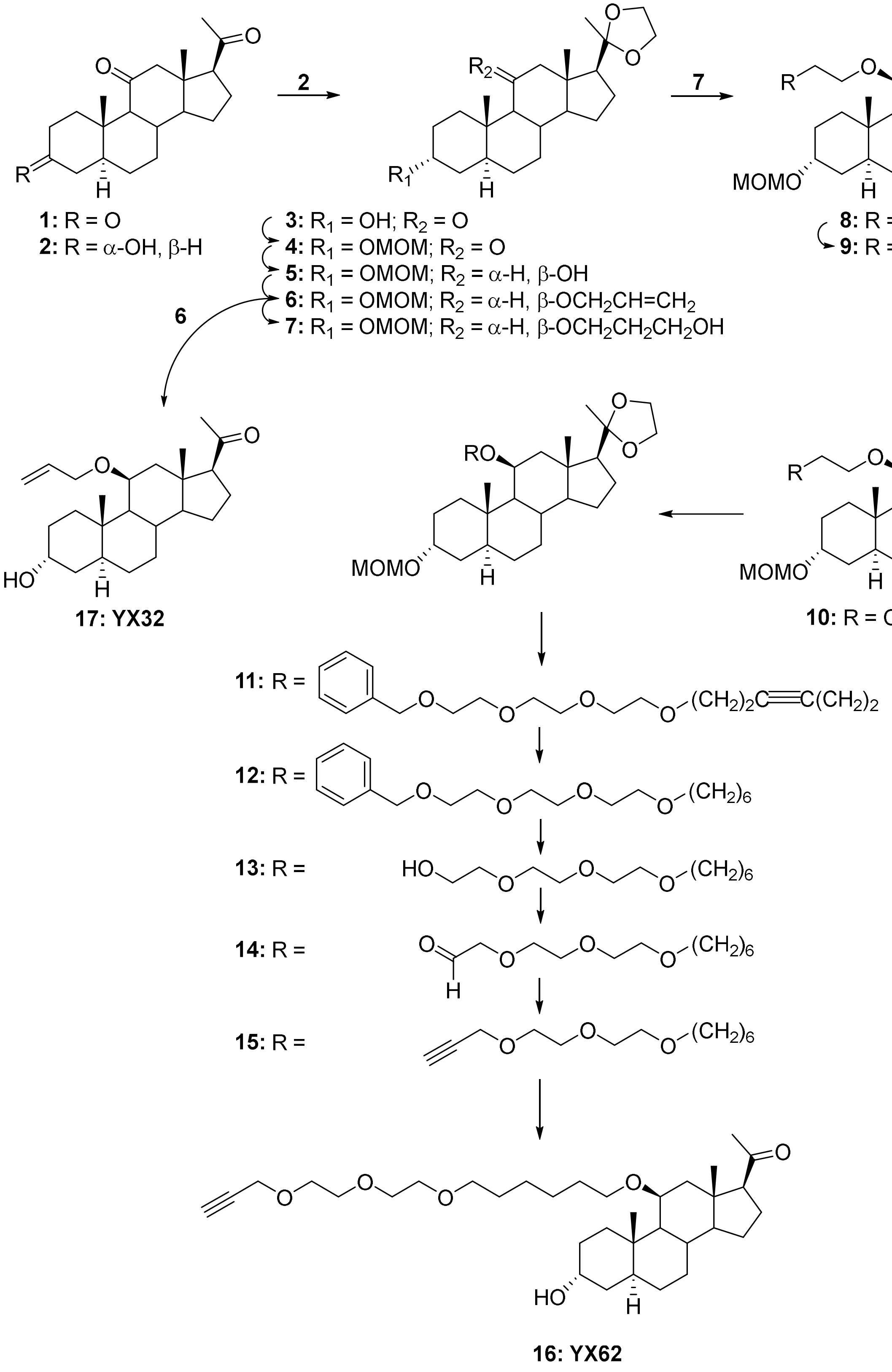

**3α-Hydroxy-5α-pregnane-11,20-dione (2).** To a stirred solution of 5α-pregnan-3,11,20-trione (**1**, 5 g, 15 mmol) in dry THF (150 mL) K-selectride (18 mL, 18 mmol, 1.0M in THF) was slowly added dropwise at -78 °C over ~30 min. After 2 h, 3 N NaOH (15 mL) and $H_2O_2$ (5 mL) were added dropwise. The reaction was allowed to warm to 23 °C and stirring continued for 1 h. The product was extracted into EtOAc (100 mL x 3). The combined extracts were washed with brine (100 mL x 5), dried over anhydrous $Na_2SO_4$, filtered and the solvent removed. The residue was purified by flash column chromatography (silica gel, eluted with 25% EtOAc and 25% $CH_2Cl_2$ in hexanes) to give steroid **2** (4.3 g, 86%): $^1$H NMR (400 MHz, $CD_3OD$) δ 4.11-4.03 (m, 1H), 2.75-2.70 (m, 1H), 2.56-0.84 (m, 21H), 2.08 (s, 3H), 0.99 (s, 3H), 0.56 (s, 3H); $^{13}$C NMR (100 MHz, $CD_3OD$) δ 209.8, 208.2, 66.2, 64.2, 62.1, 56.7, 55.8, 47.2, 38.8, 36.6, 35.7, 35.2, 32.6, 31.3, 30.8, 28.8, 27.7, 23.8, 23.2, 14.2, 10.8.

**3α-Hydroxy-17β-(2-methyl-1,3-dioxolan-2-yl)-5α-androstan-11-one (3).** To a stirred solution of steroid **2** (4.3 g, 13 mmol) in benzene (150 mL) was added ethylene glycol (12 mL) and PPTS (300 mg) at 23 °C. The reaction was refluxed in a flask equipped with a Dean-Stark apparatus for 16 h. The flask was cooled to 23 °C and aqueous $NaHCO_3$ was added. The product was extracted into EtOAc (200 mL × 2). The combine extracts were washed with brine (100 mL × 5), dried over anhydrous $Na_2SO_4$, filtered and the solvent removed. The residue was dried under high vacuum to give steroid **3** (4.5 g, 92%): $^1$H NMR (400 MHz, $CDCl_3$) δ 4.19-4.04 (m, 1H), 4.0-3.74 (m, 4H), 2.59-2.56 (m, 1H), 2.28-1.09 (m, 21H), 1.25 (s, 3H), 1.00 (s, 3H), 0.71 (s, 3H); $^{13}$C NMR (100 MHz, $CDCl_3$) δ 211.7, 111.3, 66.3, 64.9, 64.2, 63.2, 57.8, 56.9, 55.6, 46.0, 39.0, 36.3, 35.7, 35.3, 32.5, 30.9, 28.9, 27.9, 24.3, 23.5, 23.3, 14.1, 10.9.

**3α-(Methoxymethoxy)-17β-(2-methyl-1,3-dioxolan-2-yl)-5α-androstan-11-one (4).** To a stirred solution of steroid **3** (2 g, 5.3 mmol) in $CH_2Cl_2$ (100 mL) was added chloromethyl methyl ether (1.27 g, 15.9 mmol), (*i*-Pr)$_2$NEt (3.4 g, 26.5 mmol) and DMAP (5 mg) at 0 °C. The reaction was warmed to 23 °C and stirring continued for 16 h. The solvent was removed and the residue was purified by flash column chromatography (silica gel, eluted with 20%-40% EtOAc in hexanes) to give steroid **4** (2.2 g , 98%): $^1$H NMR (400 MHz, $CDCl_3$) δ 4.65-4.60 (m, 2H), 3.97-3.79 (m, 5H), 3.33 (s, 3H), 2.56-2.53 (m, 1H), 2.24-1.13 (m, 20H), 1.23 (s, 3H), 0.99 (s, 3H), 0.69 (s, 3H); $^{13}$C NMR (100 MHz, $CDCl_3$) δ 211.6, 111.2, 94.4, 71.4, 64.8, 64.2, 63.1, 57.8, 56.8, 55.6, 55.1, 46.0, 39.5, 36.3, 35.4, 33.2, 32.4, 31.4, 27.9, 25.9, 24.3, 23.4, 23.2, 14.0, 11.0.

**3α-(Methoxymethoxy)-17β-(2-methyl-1,3-dioxolan-2-yl)-5α-androstan-11β-ol (5).** Steroid **4** (2.2 g, 5.2 mmol) was dissolved in stirred dry $Et_2O$ and added dropwise to a suspension of LAH (600 mg, 15.7 mmol) in $Et_2O$ at 0 °C. The mixture was warmed to 23 °C and stirring continued for 2 h. Water (0.6 mL) was added, followed by aqueous 15% NaOH (0.6 mL) and stirring continued for 0.5 h. Additional water (1.8mL) was then added and stirring continued for 15 min. Anhydrous solid $MgSO_4$ was added and after 15 min the mixture was filtered through celite. Solvent was removed from the filtrate and the residue was purified by flash column chromatography (silica gel, eluted with 20-30% EtOAc in hexanes) to give steroid **5** (1.87 g, 85%): $^1$H NMR (400 MHz, $CDCl_3$) δ 4.68-4.66 (m, 2H), 4.30-4.29 (m, 1H), 3.99-3.84 (m, 4H), 3.74 (s, 1H), 3.37 (s, 3H), 2.56-2.53 (m, 1H), 2.21-2.17 (m, 2H), 1.80-0.82 (m, 19H), 1.29 (s, 3H), 1.04 (s, 3H), 0.98 (s, 3H); $^{13}$C NMR (100 MHz, $CDCl_3$) δ 111.9, 94.5, 71.4, 68.3, 65.0, 63.2, 58.6, 58.1, 57.9, 55.2, 48.5, 41.3, 40.6, 40.0, 33.0, 32.5, 32.2, 30.7, 28.0, 26.0, 24.5, 23.7, 22.7, 15.8, 14.5.

**3α-(Methoxymethoxy)-11β-(2-propenyloxy)-17β-(2-methyl-1,3-dioxolan-2-yl)-5α-androstane (6).** To a stirred solution of steroid **5** (1.75 g, 4.13 mmol) in THF (50 mL) KH (1 g, 25 mmol) was added at 23 °C and then the reaction was refluxed for 50 min. Ally bromide (5 mL) was added and stirring was continued for 2 h. After cooling to 23 °C, MeOH (5 mL) was added and stirring was continued for 10 min. Water (50 mL) was added and the product was extracted into EtOAc (50 mL× 3). The combined extracts were dried over anhydrous $Na_2SO_4$, filtered and the solvent removed. The residue was purified by flash column chromatography (silica gel, eluted with 5% EtOAc in hexanes) to give steroid **6** (1.58 g, 83%): $^1$H NMR (400 MHz, $CDCl_3$) δ 5.94-5.81 (m, 1H), 5.26-5.21 (m, 1H), 5.08-5.05 (m, 1H), 4.66-4.62 (m, 2H), 4.10-3.68 (m, 8H), 3.36 (s, 3H), 2.50-2.47 (m, 1H), 1.81-0.63 (m, 20H), 1.30 (s, 3H), 1.00 (s, 3H), 0.92 (s, 3H); $^{13}$C NMR (100 MHz, $CDCl_3$) δ 135.7, 115.2, 112.0, 94.4, 75.1, 71.5, 69.0, 65.1, 63.2, 58.5, 58.2, 58.2, 55.1, 41.5, 40.6, 40.6, 36.0, 33.0, 32.6, 32.4, 31.3, 28.0, 26.0, 24.5, 23.6, 22.7, 14.4, 14.4.

**3-(((3α-Methoxymethoxy)-17β-(2-methyl-1,3-dioxolan-2-yl)-5α-androstan-11β-yl)oxy)-propan-1-ol (7).** To a stirred solution of steroid **6** (1.42 g, 3.1 mmol) in THF (10mL) was added 9-BBN (12 mL, 6 mmol, 0.5 M in THF) at 0 °C. After 4 h, the reaction was allowed to warm to 23 °C and stirring was continued for 16 h. 3 N NaOH (5 mL, 15 mmol) and $H_2O_2$ (5mL) were added and the reaction was stirred for another 1 h. The product was extracted into EtOAc (100mL× 2) and the combined extracts were washed with brine (50 mL × 3), dried over anhydrous $Na_2SO_4$, filtered and the solvent removed. The residue was purified by flash column chromatography (silica gel, eluted with 10-20% EtOAc in hexanes) to give steroid **7** (1.11 g, 74%): $^1$H NMR (400 MHz, $CDCl_3$) δ 4.66-4.64 (m, 2H), 4.00-3.71 (m, 9H), 3.35 (s, 3H), 3.30-3.27 (m, 1H), 2.51-2.48 (m, 1H), 2.23-0.80 (m, 23H), 1.29 (s, 3H), 0.98 (s, 3H), 0.91 (s, 3H); $^{13}$C NMR (100 MHz, $CDCl_3$) δ 111.9, 94.4, 76.4, 71.5, 67.1, 65.1, 63.6, 63.2, 61.9, 58.5, 58.1, 55.1, 41.4, 40.6, 40.5, 35.9, 33.0, 32.7 (2 x C), 32.3, 31.2, 27.9, 26.0, 24.4, 23.5, 22.6, 14.6, 14.5.

**3α-(Methoxymethoxy)-11β-(3-oxopropoxy)-5α-pregnan-20-one (8).** To a stirred solution of oxalyl chloride (2 mL) in $CH_2Cl_2$ (30 mL) was added DMSO (2.4 mL) in $CH_2Cl_2$ (10 mL) at -78 °C. After 10 min, steroid **7** (1.11 g, 2.3 mmol) in $CH_2Cl_2$ (10 mL + 3 mL + 3 mL) was added. After 2 h, $Et_3N$ (4.8mL) was added and the reaction was allowed to warm to 23 °C and stirring was continued for 30 min. The product was extracted into $CH_2Cl_2$ (150 mL). The extract was dried over anhydrous $Na_2SO_4$, filtered and the solvent removed. The residue was purified by flash column chromatography (silica gel, eluted with 20-50% EtOAc in hexanes) to give steroid **8** (700 mg, 70%): $^1$H NMR (400 MHz, $CDCl_3$) δ 9.74-9.73 (m, 1H), 4.65-4.61 (m, 2H), 3.92-3.81 (m, 3H), 3.48-3.46 (m, 1H), 3.34 (s, 3H), 2.60-2.57 (m, 2H), 2.47-2.38 (m, 2H), 2.10 (s, 3H), 2.08-0.74 (m, 19H), 0.89 (s, 3H), 0.72 (s, 3H); $^{13}$C NMR (100 MHz, $CDCl_3$) δ 209.6, 201.6, 94.5, 75.8, 71.4, 63.9, 62.0, 58.2, 58.0, 55.1, 44.2, 43.5, 40.6, 39.9, 36.0, 33.0, 32.7, 32.4, 31.6, 31.4, 27.9 26.0, 24.2, 22.6, 14.7, 14.3.

**11β-(But-3-yn-1-yloxy)-3α-(methoxymethoxy)-5α-pregnan-20-one (9).** To a solution of steroid **8** (700 mg, 1.6 mmol) in MeOH/THF (10 mL/10 mL) was added dimethyl (1-diazo-2-oxopropyl) phosphonate (340 mg, 1.76 mmol) and $K_2CO_3$ (1 g, fine powder) at 0 °C. After 4 h, water was added and the product was extracted into EtOAc (100 mL × 3). The combined extracts were dried over anhydrous $Na_2SO_4$, filtered and the solvent removed. The residue was purified by flash column chromatography (silica gel, eluted with 20% EtOAc in hexanes) to give steroid **9** (580 mg, 81%): $^1$H NMR (400 MHz, $CDCl_3$) δ 4.59 (s, 2H), 3.78 (s, 2H), 3.64-3.58 (m, 1H), 3.31 (s, 3H), 3.25-3.20 (m, 1H), 2.40-2.33 (m, 4H), 2.11-0.78 (m, 20H), 2.06 (s, 3H), 0.94 (s, 3H), 0.72 (s, 3H); $^{13}$C NMR (100 MHz, $CDCl_3$) δ 209.36, 94.4, 81.9, 75.5, 71.4, 69.2, 66.6, 64.0, 58.3, 58.0, 55.1, 43.5, 40.5, 40.1, 36.0, 33.0, 32.5, 32.4, 31.6, 31.4, 27.9, 26.0, 24.2, 22.5, 20.1, 14.6, 14.2.

**11β-(But-3-yn-1-yloxy)-3α-(methoxymethoxy)-17β-(2-methyl-1,3-dioxolan-2-yl)-5α-androstane (10).** To a stirred solution of steroid **9** (580 mg, 1.3 mmol) in benzene (80 mL) was added ethylene glycol (1 mL) and PTSA (30 mg) at 23 °C. The reaction was refluxed in a flask equipped with a Dean-Stark apparatus for 16 h. The flask was cooled to 23 °C and aqueous $NaHCO_3$ was added. The product was extracted into EtOAc (50 mL × 2). The combined extracts were washed with brine (50 mL × 5), dried over anhydrous $Na_2SO_4$, filtered and the solvent was removed. The residue was purified by flash column chromatography (silica gel, eluted with 20% EtOAc in hexanes) to give steroid **10** (280 mg, 45%): $^1$H NMR (400 MHz, $CDCl_3$) δ 4.65-4.61 (m, 2H), 3.98-3.64 (m, 7H), 3.34 (s, 3H), 3.26-3.21 (m, 1H), 2.47-0.77 (m, 24H), 1.28 (s, 3H), 0.97 (s, 3H), 0.89 (s, 3H); $^{13}$C NMR (100 MHz, $CDCl_3$) δ 112.0, 94.4, 82.1, 75.8, 71.5, 69.0, 66.5, 65.1, 63.3, 58.5, 58.1, 58.1, 55.1, 41.5, 40.7, 40.6, 36.0, 33.1, 32.5, 32.4, 31.3, 28.0, 26.0, 24.5, 23.6, 22.7, 20.2, 14.3, 14.2.

**3α-(Methoxymethoxy)-17β-(2-methyl-1,3-dioxolan-2-yl)-11β-((1-phenyl-2,5,8,11-tetraoxaheptadec-14-yn-17-yl)oxy)-5α-androstane (11).** To a mixture of steroid **10** (280 mg, 0.59 mmol), 13-iodo-1-phenyl-2,5,8,11-tetraoxatridecane (400 mg, 1.02 mmol), Cu(I)I (7.7 mg, 0.04 mmol), $CsCO_3$ (332 mg, 1.02 mmol) and bis[(2-dimethylamino)phenyl]amine nickel(II) chloride (48 mg, 0.14 mmol) was added dioxane (12 mL). The stirred reaction was heated up to 120 °C for 16 h. The reaction was cooled to 23 °C and the solvent was removed. The residue was purified by flash column chromatography (silica gel, eluted with 20-50% EtOAc in hexanes) to give steroid **11** (370 mg, 85%): $^1$H NMR (400 MHz, $CDCl_3$) δ 7.32-7.26 (m, 5H), 4.65-4.63 (m, 2H), 4.61-4.55 (m, 2H), 3.98-3.71 (m, 20H), 3.34 (s, 3H), 3.18-3.16 (m, 1H), 2.46-2.33 (m, 5H), 1.75-0.05 (m, 21H), 1.28 (s, 3H), 0.97 (s, 3H), 0.89 (s, 3H); $^{13}$C NMR (100 MHz, $CDCl_3$) δ 138.1, 128.3 (2 x C), 127.6 (2 x C), 127.5, 111.9, 94.4, 78.8, 75.5, 73.1, 71.5, 70.5 (4 x C), 70.4, 70.1, 69.8, 69.3, 66.9, 65.0, 63.2, 58.4, 58.1, 58.0, 55.0, 41.5, 40.6, 40.5, 35.9, 33.0, 32.4, 32.3, 31.2, 28.0, 26.1, 24.4, 23.5, 22.6, 20.4, 19.9, 14.2, 14.1.

**3α-(Methoxymethoxy)-17β-(2-methyl-1,3-dioxolan-2-yl)-11β-((1-phenyl-2,5,8,11-tetraoxaheptadecan-17-yl)oxy)-5α-androstane (12).** To a stirred solution of steroid **11** (370 mg, 0.5 mmol) in EtOAc (30 mL) was added Pd/C (10%, 70 mg) in a Parr hydrogenation flask and hydrogenation ($H_2$, 55 psi) was carried out overnight. The mixture was filtered through celite and the celite was washed with EtOAc. The solvent was removed and the residue was purified by flash column chromatography (silica gel, eluted with 20-50% EtOAc in hexanes) to give steroid **12** (290 mg, 78%): $^{1}$H NMR (400 MHz, $CDCl_3$) δ 7.27-7.21 (m, 5H), 4.58-4.59 (m, 2H), 4.49-4.50 (m, 2H), 3.93-3.36 (m, 21H), 3.29 (s, 3H), 3.02-3.00 (m, 1H), 2.43-2.40 (m, 1H), 1.71-0.71 (m, 28H), 1.24 (s, 3H), 0.93 (s, 3H), 0.85 (s, 3H); $^{13}$C NMR (100 MHz, $CDCl_3$) δ 138.2, 128.3 (2 x C), 127.7 (2 x C), 127.5, 111.9, 94.4, 75.4, 73.1, 71.5, 71.4, 70.6 (5 x C), 70.0, 69.4, 68.1, 65.1, 63.2, 58.5, 58.2, 55.0, 41.5, 40.6, 40.6, 36.0, 33.0, 32.5, 32.5, 31.3, 30.3, 29.6, 28.1, 26.4, 26.1, 25.9, 24.5, 23.6, 22.6, 14.3, 14.2.

**2-(2-(2-((6-((-3α-(Methoxymethoxy)-17β-(2-methyl-1,3-dioxolan-2-yl)-5α-androstan-11β-yl)oxy)hexyl)oxy)ethoxy)ethoxy)-ethan-1-ol (13).** To a solution of anhydrous liquid ammonia (30 mL) was added sodium metal (100 mg) at -78 °C. The mixture was stirred 15 min and then steroid **12** (290 mg, 0.39 mmol) in THF (10 mL + 2 mL + 2 mL) was added dropwise. After 1 h, solid $NH_4Cl$ (2.0 g) was added. The mixture was allowed to warm to 23 °C and the ammonia allowed to evaporate. Water was added and the product was extracted into EtOAc (100 mL × 3). The combined extracts were dried over anhydrous $Na_2SO_4$, filtered, and the solvent removed. The residue was purified by flash column chromatography (silica gel, eluted with 3-5% MeOH in $CH_2Cl_2$) to give steroid **13** (260 mg, 100%): $^{1}$H NMR (400 MHz, $CDCl_3$) δ 4.61-4.57 (m, 2H), 3.94-3.36 (m, 21H), 3.30 (s, 3H), 3.02-3.00 (m, 1H), 2.42-2.39 (m, 1H), 1.70-0.70 (m, 29H), 1.24 (s, 3H), 0.92 (s, 3H), 0.84 (s, 3H); $^{13}$C NMR (100 MHz, $CDCl_3$) δ 112.0, 94.4, 75.4, 72.5, 71.6, 71.4, 70.6, 70.5, 70.3, 70.0, 68.1, 65.1, 63.2, 61.6, 58.5, 58.1 (2 x C), 55.1, 41.5, 40.6, 40.5, 35.9, 33.1, 32.5, 32.4, 31.3, 30.3, 29.5, 28.1, 26.4, 26.0, 25.9, 24.5, 23.6, 22.6, 14.3, 14.2.

**2-(2-(2-((6-((-3α-(Methoxymethoxy)-17β-(2-methyl-1,3-dioxolan-2-yl)-5α-androstan-11β-yl)oxy)hexyl)oxy)ethoxy)ethoxy)-acetaldehyde (14).** To a stirred solution of oxalyl chloride (0.5 mL) in $CH_2Cl_2$ (10 mL) was added DMSO (0.6 mL) in $CH_2Cl_2$ (10 mL) at -78°C. After 10 min, steroid **13** (260 mg, 0.4 mmol) in $CH_2Cl_2$ (3 mL + 2 mL + 1 mL) was added. After 2 h, $Et_3N$ (4 mL) was added and the reaction was allowed to warm to 23 °C and stirring was continued for 30 min. The product was extracted into $CH_2Cl_2$ (50 mL). The extract was dried over anhydrous $Na_2SO_4$, filtered and the solvent removed. The residue was purified by flash column chromatography (silica gel, eluted with 20-50% EtOAc in hexanes) to give steroid **14** (190 mg, 73%): $^{1}$H NMR (400 MHz, $CDCl_3$) δ 9.6 (s, 1H), 4.56 (s, 2H), 4.08-3.34 (m, 19H), 3.27 (s, 3H), 3.00-2.98 (m, 1H), 2.40-2.37 (m, 1H), 1.66-0.69 (m, 28H), 1.21 (s, 3H), 0.90 (s, 3H), 0.82 (s, 3H);

$^{13}C$ NMR (100 MHz, $CDCl_3$) δ 200.8, 111.9, 94.4, 76.8, 75.4, 71.5, 71.4, 71.1, 70.7, 70.6, 69.9, 68.1, 65.1, 63.2, 58.5, 58.1 (2 x C), 55.0, 41.5, 40.6, 40.5, 35.9, 33.0, 32.5, 32.4, 31.3, 30.2, 29.5, 28.0, 26.4, 26.0, 25.9, 24.5, 23.6, 22.6, 14.3, 14.2.

**2-(3α-(Methoxymethoxy)-11β-((6-(2-(2-(prop-2-yn-1-yloxy)ethoxy)ethoxy)hexyl)oxy)-5α-androstan-17β-yl)-2-methyl-1,3-dioxolane (15).** To a stirred solution of steroid **14** (190 mg, 0.29 mmol) in MeOH/THF (5 mL/5 mL) was added dimethyl (1-diazo-2-oxopropyl) phosphonate (340 mg, 1.76 mmol) and $K_2CO_3$ (1 g, fine powder) at 23 °C. After 16 h, water was added and the product was extracted into EtOAc (50 mL × 3). The combined extracts were dried over anhydrous $Na_2SO_4$, filtered and the solvent removed. The residue was purified by flash column chromatography (silica gel, eluted with 20-50% EtOAc in hexanes) to give steroid **15** (150 mg, 79%): $^1H$ NMR (400 MHz, $CDCl_3$) δ 4.61 (s, 2H), 4.16 (s, 2H), 3.96-3.32 (m, 17H), 3.31 (s, 3H), 3.04-3.01 (m, 1H), 2.44-2.40 (m, 2H), 1.70-0.73 (m, 28H), 1.25 (s, 3H), 0.94 (s, 3H), 0.86 (s, 3H); $^{13}C$ NMR (100 MHz, $CDCl_3$) δ 112.0, 94.4, 79.6, 75.5, 74.5, 71.6, 71.4, 70.6, 70.4, 70.0, 69.1, 68.1, 65.2, 63.2, 58.6, 58.4, 58.2 (2 x C), 55.1, 41.6, 40.6, 40.6, 36.0, 33.1, 32.5, 32.4, 31.4, 30.3, 29.5, 28.1, 26.4, 26.1, 26.0, 24.5, 23.6, 22.6, 14.3, 14.2.

**3α-Hydroxy-11β-((6-(2-(2-(prop-2-yn-1-yloxy)ethoxy)ethoxy)hexyl)oxy)-5α-pregnan-20-one (16, YX62).** To a stirred solution of steroid **15** (150 mg, 0.23 mmol) in THF (5 mL) was added 6 N HCl (5 mL) at 23 °C. After 4 h, the product was extracted into EtOAc (20 mL × 3). The combined extracts were dried over anhydrous $Na_2SO_4$, filtered and the solvent removed. The residue was purified by flash column chromatography (silica gel, eluted with 50-100% EtOAc in hexanes) to give **YX62** (**16**,115 mg, 89%): $^1H$ NMR (400 MHz, $CDCl_3$) δ 4.17-4.16 (m, 2H), 4.00 (s, 1H), 3.74-3.38 (m, 12H), 3.07-3.05 (m, 1H), 2.42-2.38 (m, 3H), 2.11-0.77 (m, 28H), 2.08 (s, 3H), 0.93 (s, 3H), 0.73 (s, 3H); $^{13}C$ NMR (100 MHz, $CDCl_3$) δ 209.6, 79.5, 75.1, 74.4, 71.3, 70.5, 70.2, 69.9, 68.9, 68.2, 66.2, 63.9, 58.3, 58.2, 58.0, 43.5, 40.0, 39.7, 36.1, 35.2, 32.4, 31.8, 31.6, 31.3, 30.1, 29.4, 28.5, 27.8, 26.2, 25.8, 24.2, 22.4, 14.6, 13.8.

**3α-Hydroxy-11β-(2-propenyloxy)-5α-pregnan-20-one (17, YX32).** To a solution of steroid **6** (93 mg, 0.2 mmol) in MeOH (10mL) added 6 N HCl (2 mL) at 23 °C. The reaction was stirred for 24 h. After solvent removal, the product was extracted into EtOAc (20 mL), dried over anhydrous $Na_2SO_4$, filtered and the solvent removed. The residue was purified by flash column chromatography (silica gel, eluted with 20% EtOAc in hexanes) to give steroid **YX32** (**17,** 60 mg,

80%): $^1$H NMR (400 MHz, $CDCl_3$) δ 5.92-5.85 (m, 1H), 5.25-5.20 (m, 1H), 5.09-5.06 (m, 1H), 4.08-4.04 (m, 2H), 3.86-3.85 (m, 1H), 3.74-3.70 (m, 1H), 2.47-2.41 (m, 2H), 2.18-0.83 (m, 20H), 2.12 (s, 3H), 0.98 (s, 3H), 0.78(s, 3H); $^{13}$C NMR (100 MHz, $CDCl_3$) δ 209.6, 135.2, 115.5, 74.8, 69.1, 66.4, 64.1, 58.4, 58.2, 43.5, 40.1, 39.9, 36.2, 35.3, 32.5, 32.0, 31.6, 31.4, 28.7, 27.9, 24.2, 22.5, 14.7, 14.1.

### Synthesis of YX34

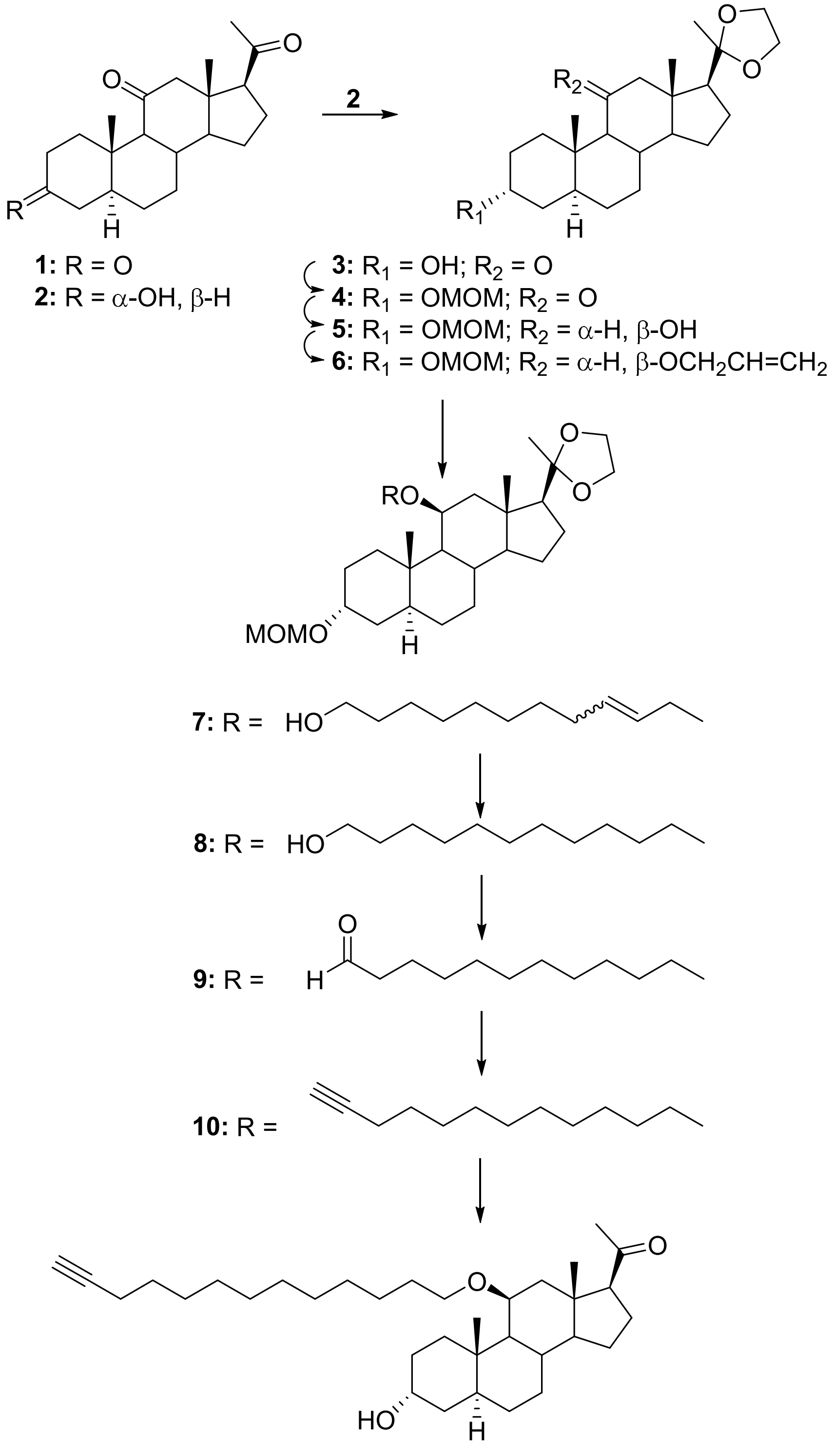


**11: YX34**

**3α-Hydroxy-5α-pregnane-11,20-dione (2).** To a solution of 5α-pregnane-3,11,20-trione (**1**, 5 g, 15 mmol) in THF (150 mL) was added dropwise K-seletride over ~20 minutes at -78 °C and stirring was continued for 2 h. After the mixture was stirred for 2 h acetone (3 mL) was added and the reaction was stirred an additional 15 min at -78 °C. After warming to 23 °C, aqueous $NH_4Cl$ was added and the mixture was stirred for 1 h. The product was extracted with EtOAc (100 mL x 3). The combined extracts were dried over anhydrous $Na_2SO_4$, filtered, the solvent removed and the residue was purified by flash column chromatography (silica gel, eluted with 25% EtOAc and 25% $CH_2Cl_2$ in hexanes) to give steroid **2** (4.3 g, 86%): $^1$H NMR (400 MHz, $CDCl_3$) δ 4.03 (s, 1H), 2.75-2.70 (m, 1H), 2.56-2.45 (m, 2H), 2.23-2.18 (m, 2H), 2.08 (s, 3H), 1.81-0.84 (m, 17H), 0.99 (s, 3H), 0.56 (s, 3H); $^{13}$C NMR (100 MHz, $CDCl_3$) δ 209.8, 208.2, 66.2, 64.2, 62.1, 56.7, 55.8, 47.2, 38.8, 36.6, 35.7, 35.2, 32.5, 31.3, 30.8, 28.8, 27.7, 23.8, 23.2, 14.2, 10.8.

**3α-Hydroxy-17β-(2-methyl-1,3-dioxolan-2-yl)-5α-androstan-11-one (3).** To a solution of steroid **2** (4.3 g, 13 mmol) in benzene (150 mL) was added ethylene glycol (12 mL) and PPTS (300 mg) at 23 °C. The reaction was refluxed in a flask equipped with a Dean-Stark apparatus for 16 h. After cooling to 23 °C, aqueous $NaHCO_3$ was added and the product was extracted into EtOAc (200 mL x 2). The combined extracts were washed with brine (100mL× 5), dried over anhydrous $Na_2SO_4$, filtered and the residue was dried under high vacuum to give steroid **3** (4.5 g, 92%): $^1$H NMR (400 MHz, $CDCl_3$) δ 4.04 (s, 1H), 3.99-2.84 (m, 4H), 2.59-2.25 (m, 2H), 2.10-1.09 (m, 20H),1.25 (s, 3H), 1.00 (s, 3H), 0.71 (s, 3H); $^{13}$C NMR (100 MHz, $CDCl_3$) δ 211.7, 111.3, 66.3, 64.9, 64.2, 63.2, 57.8, 56.9, 55.6, 46.0, 40.0, 36.3, 35.7, 35.3, 32.5, 30.9, 28.9, 27.9, 24.3, 23.5, 23.3, 14.1, 10.9.

**17β-(2-Methyl-1,3-dioxolan-2-yl)-3α-(methoxymethoxy)-5α-androstan-11-one (4).** To a solution of steroid **3** (2.0 g, 5.3 mmol) in $CH_2Cl_2$ (100 mL) was added (*i*-Pr)$_2$EtN (4.6 mL) at 23 °C. After cooling to 0 °C, MOMCl (1.3 mL, 15.9 mmol) was added and the reaction was stirred at 23 °C for 16 h. The mixture was warmed to 23 °C and was stirred for 16 h. Aqueous $NaHCO_3$ was added to consume excess MOMCl and the product was extracted into $CH_2Cl_2$ (100 mL x 2). The combined extracts were dried over anhydrous $Na_2SO_4$, filtered, the solvent removed and the residue was purified by flash column chromatography (silica gel, eluted with 25%-35% EtOAc in hexanes) to give steroid **4** (2.2 g, 98%): $^{1}$H NMR (400 MHz, $CDCl_3$) δ 4.65-4.60 (m, 2H), 3.97-3.79 (m, 5H), 3.33 (s, 3H), 2.56-2.53 (m, 1H), 2.24-1.11 (m, 20H), 1.23 (s, 3H), 0.99 (s, 3H), 0.69 (s, 3H); $^{13}$C NMR (100 MHz, $CDCl_3$) δ 211.57, 111.2, 94.4, 71.4, 64.8, 64.2, 63.1, 57.8, 56.8, 55.6, 55.1, 46.0, 39.5, 36.3, 35.4, 33.2, 32.4, 31.4, 27.9, 25.9, 24.3, 23.4, 23.2, 14.0, 11.0.

**17β-(2-Methyl-1,3-dioxolan-2-yl)-3α-(methoxymethoxy)-5α-androstan-11β-ol (5).** Steroid **4** (2.2 g, 5.2 mmol) was dissolved in anhydrous $Et_2O$ and added dropwise to a suspension of $LiAlH_4$ (600 mg, 15.7 mmol) in anhydrous $Et_2O$ at °C. The mixture was warmed to 23 °C and stirred for 2 h. Water (0.6 mL) was added followed by aqueous 15% NaOH (0.6 mL) and stirring continued for 0.5 h. Water (1.8mL) was then added and stirring continued for 15 min. Anhydrous $MgSO_4$ was added and after 15 min the mixture was filtered through celite. Solvent was removed from the filtrate and the residue was purified by flash column chromatography (silica gel, eluted with 20%-30% EtOAc in hexanes) to give steroid **5** (1.87 g, 85%): $^{1}$H NMR (400 MHz, $CDCl_3$) δ 4.68-4.66 (m, 2H), 4.30-4.29 (m, 1H), 3.99-3.84 (m, 4H), 3.74 (s, 1H), 3.37(s, 3H), 2.56-2.53 (m, 1H), 2.21-2.17 (m, 2H), 1.80-0.82 (m, 19H), 1.29 (s, 3H), 1.04 (s, 3H), 0.98 (s, 3H); $^{13}$C NMR (100 MHz, $CDCl_3$) δ 111.9, 94.5, 71.4, 68.3, 65.0, 63.2, 58.6, 58.1, 57.9, 55.2, 48.5, 41.3, 40.6, 40.0, 33.0, 32.5, 32.2, 30.7, 28.0, 26.0, 24.5, 23.7, 22.7, 15.8, 14.5.

**17β-(2-Methyl-1,3-dioxolan-2-yl)-3α-(methoxymethoxy)-11β-(2-propenyloxy)-5α-androstane (6).** To a solution of steroid **5** (1.54 g, 3.6 mmol) in THF (50mL) was added KH (1 g, 25 mmol) at 23 °C. The mixture was refluxed for 50 min., allyl bromide (5mL) was added and stirring continued for 2 h. After cooling to 23 °C, MeOH (5mL) was added and stirring was continued for 10 min. Water (50 mL) was added and the product was extracted into EtOAc (50 mL× 3). The combined extracts were dried over anhydrous $Na_2SO_4$, filtered, the solvent removed and the residue was purified by flash column chromatography (silica gel, eluted with 5% EtOAc in hexanes) to give steroid **6** (1.2 g, 71%): $^{1}$H NMR (400 MHz, $CDCl_3$) δ 5.94-5.81 (m, 1H), 5.26-5.21 (m, 1H), 5.08-5.05 (m, 1H), 4.65 (s, 2H), 4.10-3.68 (m, 8H), 3.36 (s, 3H), 2.50-2.47 (m, 1H), 1.81-0.63 (m, 20H), 1.30 (s, 3H), 1.00 (s, 3H), 0.92 (s, 3H); $^{13}$C NMR (100 MHz, $CDCl_3$) δ 135.7, 115.2, 112.0, 94.4, 75.1, 71.5, 69.0, 65.1, 63.2, 58.5, 58.2, 58.2, 55.1, 41.5, 40.6, 40.6, 36.0, 33.0, 32.6, 32.4, 31.3, 28.0, 26.0, 24.5, 23.6, 22.7, 14.4, 14.4.

**((3α-(Methoxymethoxy)-17β-(2-methyl-1,3-dioxolan-2-yl)-5α-androstan-11β-yl)oxy)-undec-9-en-1-ol (7).** To a solution of steroid **6** (1.2 g, 2.6 mmol) in THF (15 mL) was added 10-undecen-1-ol (2.2 g, 13 mmol) and Grubbs catalyst M102 (30 mg, 0.05 mmol) in THF (2 mL) at 23 °C. After 16 h, THF was removed, and the residue was purified by flash column chromatography (silica gel, eluted with 10-25% EtOAc in hexanes) to give steroid **7** (E/Z olefin mixture, 510 mg, 33%): $^1$H NMR (400 MHz, $CDCl_3$) δ 5.60-5.43 (m, 2H), 4.63 (q, $J$ = 6.6 Hz, 2H), 4.05-3.55 (m, 10H), 3.32 (s, 3H), 2.48-2.41 (m, 1H), 2.11-0.74 (m, 35H), 1.26 (s, 3H), 0.95 (s, 3H), 0.88 (s, 3H); $^{13}$C NMR (100 MHz, $CDCl_3$) δ 132.6, 127.1, 111.9, 94.3, 74.6, 71.4, 68.7, 65.0, 63.1, 62.7, 58.5, 58.1, 55.0, 41.4, 40.5, 35.9, 32.9, 32.7, 32.4, 32.3, 32.1, 31.2, 29.5, 29.4, 29.3, 29.1, 29.0, 28.9, 27.9, 25.9, 25.7, 24.4, 23.5, 22.5, 14.4, 14.2.

**((3α-(Methoxymethoxy)-17β-(2-methyl-1,3-dioxolan-2-yl)-5α-androstan-11β-yl)oxy)-undecan-1-ol (8).** To a solution of steroid **7** (510 mg, 0.86 mmol) in EtOAc (50 mL) was added 10% Pd/C (100 mg) at 23 °C in a Parr hydrogenation flask. The flask was evacuated and charged with $H_2$ three times. Hydrogenation was carried out at 55 psi overnight. The mixture was filtered through Celite and washed with EtOAc (100 mL). The solvent was removed to give steroid **8** (510 mg, 100%): $^1$H NMR (400 MHz, $CDCl_3$) δ 4.65 (q, $J$ = 7.0 Hz, 2H), 4.00-3.60 (m, 8H), 3.52-3.50 (m, 1H), 3.35 (s, 3H), 3.06-3.04 (m, 1H), 2.48-2.44 (m, 1H), 1.88-0.76 (m, 39H), 1.24 (s, 3H), 0.98 (s, 3H), 0.90 (s, 3H); $^{13}$C NMR (100 MHz, $CDCl_3$) δ 112.0, 94.4, 75.4, 71.6, 68.2, 65.1, 63.2, 62.9, 58.5, 58.1, 55.1, 41.5, 40.59, 40.56, 35.9, 33.1, 32.7, 32.5, 32.4, 31.3, 30.3, 29.6, 29.5, 29.48, 29.4, 29.38, 28.0, 26.5, 26.0, 25.7, 24.5, 23.6, 22.6, 14.6, 14.3, 14.1.

**((3α-(Methoxymethoxy)-17β-(2-methyl-1,3-dioxolan-2-yl)-5α-androstan-11β-yl)oxy)-undecan-1-al (9).** To a solution of steroid **8** (510 mg, 0.86 mmol) in $CH_2Cl_2$ (50 mL) was added Dess-Martin reagent (1.3 g, 3.1 mmol) at 23 °C. After 2 h, aqueous $NaHCO_3$ was added. The product was extracted into $CH_2Cl_2$ (50 mL x 2). The combined extracts were dried over anhydrous $Na_2SO_4$, filtered, the solvent removed and the residue was purified by flash column chromatography (silica gel, eluted with 10-25% EtOAc in hexanes) to give steroid **8** (340 mg, 67%): $^1$H NMR (400 MHz, CDCl3) δ 9.77 (s, 1H), 4.67 (s, 2H), 4.00-3.71 (m, 5H), 3.53-3.51 (m, 1H), 3.37 (s, 3H), 3.06-3.01 (m, 1H), 2.43-2.38 (m, 4H), 2.01 (s, 3H), 1.77-0.73 (m, 33H), 1.26 (s, 3H), 1.00 (s, 3H), 0.92 (s, 3H); $^{13}$C NMR (100 MHz, $CDCl_3$) δ 112.0, 94.5, 75.5, 71.6, 68.2, 65.2,

63.3, 58.6, 58.2, 55.1, 43.9, 41.6, 40.6, 36.0, 33.1, 32.6, 32.54, 32.47, 31.4, 30.4, 29.6, 29.5, 29.49, 29.4, 29.3, 29.1, 28.1, 26.6, 26.5, 26.1, 24.5, 23.6, 22.1, 14.7, 14.3, 14.2.

**17β-(2-Methyl-1,3-dioxolan-2-yl)-3α-(methoxymethoxy)-11β-(tridec-12-yn-1-yloxy)-5α-androstane (10).** To a solution of steroid **9** (340 mg, 0.54 mmol) in MeOH/THF (5 mL/5 mL) was added dimethyl (1-diazo-2-oxopropyl) phosphonate (520 mg, 2.7 mmol) and $K_2CO_3$ (0.7 g, fine powder) at 23 °C. After 16 h, water was added, and the product was extracted into EtOAc (50mL x 3). The combined extracts were dried over anhydrous $Na_2SO_4$, filtered, the solvent removed and the residue was purified by flash column chromatography (silica gel, eluted with 10 % EtOAc in hexanes) to give steroid **10** (280 mg, 86%): $^1$H NMR (400 MHz, $CDCl_3$) δ 4.66 (q, $J$ = 7.0 Hz, 2H), 4.00-3.81 (m, 5H), 3.68-3.67 (m, 1H), 3.52-3.50 (m, 1H), 3.35 (s, 3H), 3.07-3.05 (m, 1H), 2.48-2.45 (m, 1H), 2.19-2.15 (m, 2H), 1.93-1.92 (m, 1H), 1.77-0.79 (m, 38H), 1.26 (s, 3H), 0.99 (s, 3H), 0.90 (s, 3H); $^{13}$C NMR (100 MHz, $CDCl_3$) δ 112.0, 94.4, 84.7, 75.4, 71.6, 68.2, 68.0 65.2, 63.2, 58.6, 58.2, 55.1, 41.5, 40.6, 40.57, 35.9, 33.1, 32.5, 32.4, 31.3, 30.3, 29.53, 29.51, 29.48, 29.4, 29.1, 28.7, 28.4, 28.1, 26.5, 26.0, 24.5, 23.6, 22.6, 18.3, 14.3, 14.1.

**3α-Hydroxy-11β-(tridec-12-yn-1-yloxy)-5α-pregnan-20-one (11, YX34).** To a solution of steroid **10** (280 mg, 0.47 mmol) in MeOH (20 mL) was added 6 N HCl (4mL). After 16 h, the product was extracted into EtOAc (20 mL x 3). The combined extracts were dried over anhydrous $Na_2SO_4$, filtered, the solvent removed and the residue was purified by flash column chromatography (silica gel, eluted with 20% EtOAc in hexanes) to give **YX34** (**11**, 210 mg, 88%): $^1$H NMR (400 MHz, $CDCl_3$) δ 4.02 (s, 1H), 3.753-3.747 (m, 1H), 3.49-3.47 (m, 1H), 3.10-3.06 (m, 1H), 2.44-2.40 (m, 2H), 2.17-2.10 (m, 6H), 1.92-1.91 (m, 1H), 1.77-0.78 (m, 34H), 1.24 (s, 3H), 0.96 (s, 3H), 0.75 (s, 3H); $^{13}$C NMR (100 MHz, $CDCl_3$) δ 209.6, 84.7, 75.1, 68.3, 68.0, 66.3, 64.0, 58.4, 58.1, 43.6, 40.0, 39.8, 36.2, 35.3, 32.5, 31.9, 31.6, 31.4, 30.2, 29.5, 29.45, 29.41, 29.4, 29.0, 28.7, 28.6, 28.4, 27.9, 26.4, 24.2, 22.4, 18.3, 14.6, 13.9.

**Synthesis of YX47 and YX85**

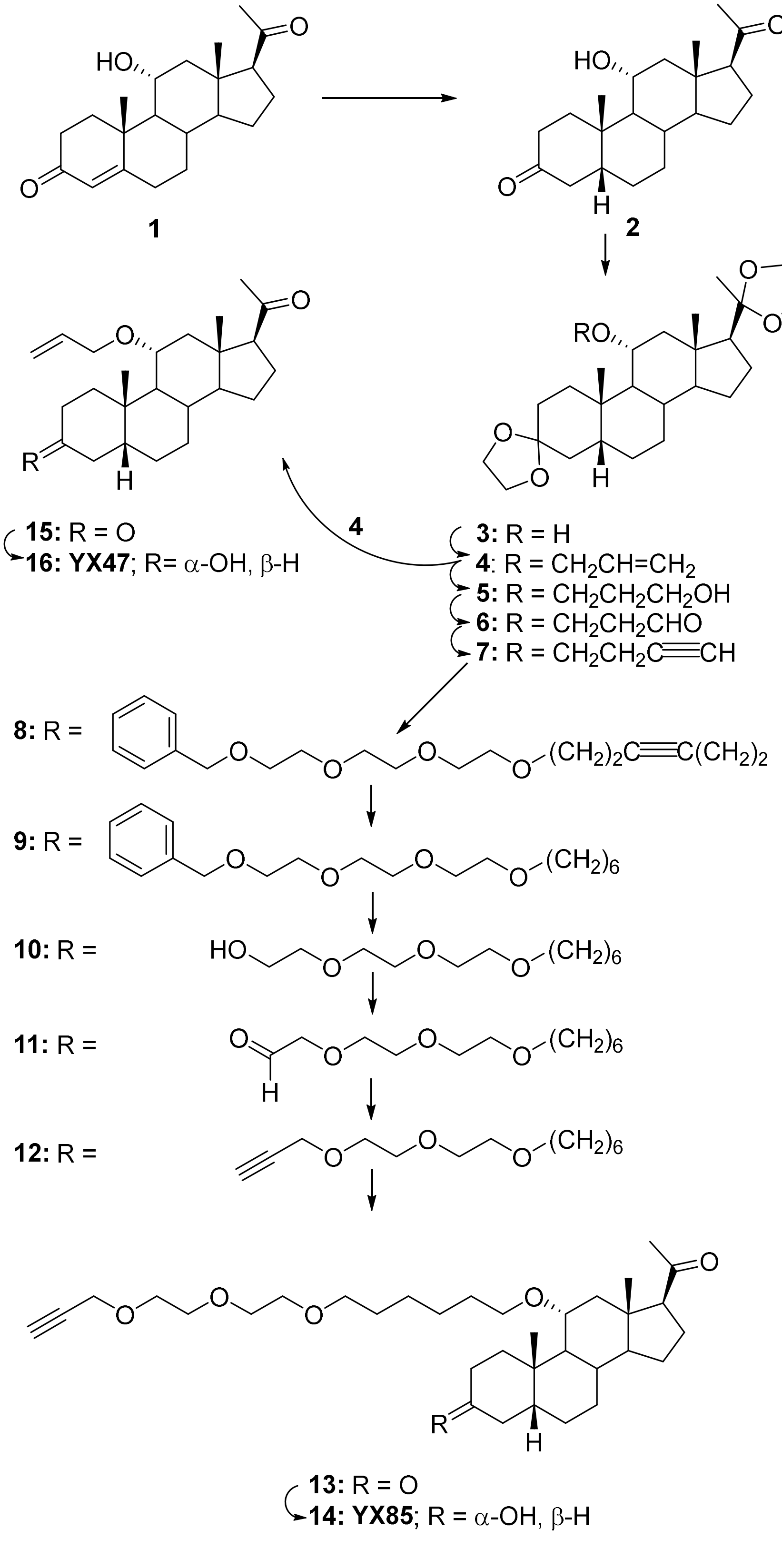

**11α-Hydroxy-5β-pregnane-3,20-dione (2).** To a stirred solution of 11α-hydroxy-pregn-4-ene-3,20-dione (**1**, 4.0 g, 12.1 mmol) in pyridine (50 mL) was added Pd/$CaCO_3$ (5%, 500 mg) in a Parr hydrogenation flask and hydrogenation was carried out overnight at 55 psi $H_2$. The mixture was filtered through celite and washed with $CH_2Cl_2$ (200 mL). The solvent was removed and the residue was purified by flash column chromatography (silica gel, eluted with 50% EtOAc in hexanes) to give steroid **2** (2.75 g, 69%): $^1$H NMR (400 MHz, $CDCl_3$) δ 3.91-3.85 (m, 1H), 2.74-1.08 (m, 22H), 2.04 (s, 3H), 1.04 (s, 3H), 0.55 (s, 3H); $^{13}$C NMR (100 MHz, $CDCl_3$) δ 214.5, 209.2, 68.1, 63.1, 55.3, 50.0, 46.6, 45.7, 43.9, 42.4, 39.7, 38.2, 35.7, 34.1, 31.2, 26.6, 25.3, 24.1, 22.8, 22.6, 14.2.

**17β-(2-Methyl-1,3-dioxolan-2-yl)-spiro[5β-androstane-3,2'-[1,3]dioxolan]-11α-ol (3).** To a stirred solution of steroid **2** (2.2 g, 6.6 mmol) in benzene (150 mL) was added ethylene glycol (3 mL) and PTSA (100 mg) at 23 °C. The reaction was refluxed in a flask equipped with a Dean-Stark apparatus for 4 h. After cooling to 23 °C, aqueous $NaHCO_3$ was added and the product was extracted into EtOAc (200 mL x 2). The combined extracts were washed with brine (100 mL × 5), dried over anhydrous $Na_2SO_4$, filtered, solvent removed and the residue was dried under high vacuum to give steroid **3** (2.6 g, 94%): $^1$H NMR (400 MHz, $CDCl_3$) δ 4.00-3.84 (m, 9H), 2.47-2.44 (m, 1H), 2.36-2.32 (m, 1H), 2.03-0.74 (m, 20H), 1.28 (s, 3H), 1.06 (s, 3H), 0.74 (s, 3H); $^{13}$C NMR (100 MHz, $CDCl_3$) δ 111.6, 110.3, 69.0, 65.1, 64.2, 64.0, 63.2, 58.0, 55.4, 51.6, 46.8, 42.53, 42.48, 37.1, 36.2, 35.8, 34.1, 31.2, 27.1, 26.0, 24.5, 23.8, 23.6, 23.0, 14.1.

**17β-(2-Methyl-1,3-dioxolan-2-yl)-11α-(2-propenyloxy)-spiro[5β-androstane-3,2'-[1,3]dioxolane] (4).** To a stirred solution of steroid **3** (2.6 g, 6.2 mmol) in THF (50 mL) was added KH (1 g, 25 mmol) at 23 °C. The mixture was refluxed for 50 min, allyl bromide (5 mL) was added and the reaction was stirred for 2 h at reflux. After cooling to 23 °C, MeOH (5 mL) was added and stirring continued for 10 min. Water (50 mL) was added and the product was extracted into EtOAc (50 mL x 3). The combined extracts were dried over anhydrous $Na_2SO_4$, filtered, the solvent removed and the residue purified by flash column chromatography (silica gel, eluted with 5% EtOAc in hexanes) to give steroid **4** (2.3 g, 80%): $^1$H NMR (400 MHz, $CDCl_3$) δ 5.87-5.80 (m, 1H), 5.21-5.16 (m, 1H), 5.06-5.03 (m, 1H), 4.10-4.07 (m, 1H), 4.05-3.78 (m, 8H), 3.77-3.74 (m, 1H), 3.48-3.41 (m, 1H), 2.52-2.47 (m, 1H), 2.18-2.15 (m, 1H), 2.04-1.07 (m, 19H), 1.29 (s, 3H), 1.03 (s, 3H), 0.71 (s, 3H); $^{13}$C NMR (100 MHz, $CDCl_3$) δ 135.4, 115.4, 111.7, 110.4, 76.4, 68.9,

65.2, 64.1, 64.0, 63.2, 58.0, 55.2, 45.2, 44.6, 42.6, 42.0, 37.0, 36.2, 35.9, 34.4, 31.2, 27.2, 26.2, 24.5, 24.0, 23.9, 23.0, 14.1.

**3-((17β-(2-Methyl-1,3-dioxolan-2-yl)-spiro[5β-androstane-3,2'-[1,3]dioxolan]-11α-yl)oxy)propan-1-ol (5).** To a stirred solution of steroid **4** (2.3 g, 5 mmol) in THF (20 mL) was added 9-BBN (20 mL, 20 mmol, 0.5 M in THF) at 0 °C. After 4 h, the reaction was allowed to warm to 23 °C and stirring was continued for 16 h. 3 N NaOH (10 mL) and $H_2O_2$ (10 mL) were added and the reaction was stirred for 1 h. The product was extracted into EtOAc (100 mL × 2). The combined extracts were washed with brine (50 mL x 3), dried over anhydrous $Na_2SO_4$, filtered, the solvent removed and the residue purified by flash column chromatography (silica gel, eluted with 10-30% EtOAc in hexanes) to give steroid **5** (1.85 g, 77%): $^1$H NMR (400 MHz, $CDCl_3$) δ 3.91-3.76 (m, 8H), 3.58-3.57 (m, 3H), 3.33-3.28 (m, 2H), 2.56 (s, 1H), 2.46-2.42 (m, 1H), 2.03-1.01 (m, 22H), 1.20 (s, 3H), 0.94 (s, 3H), 0.62 (s, 3H); $^{13}$C NMR (100 MHz, $CDCl_3$) δ 111.6, 110.2, 76.8, 66.6, 65.0, 64.1, 63.9, 63.1, 61.5, 57.8, 55.1, 45.2, 44.6, 42.5, 42.0, 36.9, 36.2, 35.8, 34.3, 32.7, 31.0, 27.1, 26.1, 24.4, 23.9, 23.8, 23.0, 14.0.

**3-((17β-(2-Methyl-1,3-dioxolan-2-yl)-spiro[5β-androstane-3,2'-[1,3]dioxolan]-11α-yl)oxy)-propanal (6).** To a stirred solution of oxalyl chloride (0.50 mL, 6 mmol) in $CH_2Cl_2$ (30 mL) was added DMSO (0.50 mL, 7.0 mmol) in $CH_2Cl_2$ (4 mL) at -78 °C and the reaction was stirred for 15 min. Steroid **5** (1.85 g, 3.9 mmol) in $CH_2Cl_2$ (10 mL) was added and stirring continued for 1 h at -78 °C. $Et_3N$ (1.4 mL, 10 mmol) was added and stirring continued for 45 min. The reaction was allowed to warm to 23 °C and water was added. The product was extracted into $CH_2Cl_2$ (100 mL x 3). The combined extracts were dried over anhydrous $Na_2SO_4$, filtered, the solvent removed and the residue purified by flash column chromatography (silica gel, eluted with 25-40% EtOAc in hexanes) to give steroid **6** (1.48 g, 80%): $^1$H NMR (400 MHz, $CDCl_3$) δ 9.68 (s, 1H), 3.97-3.81 (m, 9H), 3.52-3.38 (m, 2H), 2.53-2.51 (m, 3H), 2.00-1.01 (m, 20H), 1.25 (s, 3H), 0.99 (s, 3H), 0.67 (s, 3H); $^{13}$C NMR (100 MHz, $CDCl_3$) δ 201.6, 111.6, 110.2, 76.7, 65.0, 64.1, 64.0, 63.2, 61.4, 57.9, 55.0, 45.0, 44.5, 44.2, 42.4, 42.0, 37.1, 36.2, 35.9, 34.3, 30.9, 27.1, 26.3, 24.4, 23.9, 23.8, 23.1, 14.0.

**11α-(But-3-yn-1-yloxy)-17β-(2-methyl-1,3-dioxolan-2-yl)-spiro[5β-androstane-3,2'-[1,3]dioxolane] (7).** To a stirred solution of steroid **6** (1.48 g, 3.1 mmol) in MeOH/THF (10 mL/10 mL) was added dimethyl (1-diazo-2-oxopropyl) phosphonate (2.4g, 12.4 mmol) and $K_2CO_3$ (2.5 g, fine powder) at 23 °C. After 16 h, water was added and the product was extracted into EtOAc (100 mL x 3). The combined extracts were dried over anhydrous $Na_2SO_4$, filtered, the solvent removed and the residue purified by flash column chromatography (silica gel, eluted with 10-20% EtOAc in hexanes) to give steroid **7** (1.24 g, 84%): $^1H$ NMR (400 MHz, $CDCl_3$) δ 4.00-3.85 (m, 8H), 3.69-3.63 (m, 1H), 3.46-3.40 (m, 1H), 3.37-3.31 (m, 1H), 2.53-2.49 (m, 1H), 2.34-2.31 (m, 2H), 2.17- 2.13 (m, 1H), 2.04-1.05 (m, 20H), 1.29 (s, 3H), 1.03 (s, 3H), 0.71 (s, 3H); $^{13}C$ NMR (100 MHz, $CDCl_3$) δ 111.7, 110.4, 81.6, 76.6, 69.1, 66.1, 65.1, 64.1, 64.0, 63.2, 58.0, 55.1, 45.22, 44.5, 42.5, 42.0, 37.2, 36.2, 35.9, 34.4, 31.1, 27.2, 26.3, 24.5, 23.9, 23.1, 20.2, 14.1.

**17β-(2-Methyl-1,3-dioxolan-2-yl)-11α-((1-phenyl-2,5,8,11-tetraoxaheptadec-14-yn-17-yl)oxy)-spiro[5β-androstane-3,2'-[1,3]dioxolane] (8).** To a mixture of steroid **7** (1.24 g, 2.6 mmol), 13-iodo-1-phenyl-2,5,8,11-tetraoxatridecane (1.2 g, 3 mmol), Cu(I)I (100 mg, 0.53 mmol), $CsCO_3$ (1.5 g, 4.6 mmol) and bis[(2-dimethylamino)phenyl]amine nickel(II) chloride (100 mg, 0.29 mmol) was added dioxane (20 mL). The stirred mixture was heated to 120 °C for 16 h. The flask was cooled to 23 °C, the solvent was removed and the residue was purified by flash column chromatography (silica gel, eluted with 20-50% EtOAc in hexanes) to give steroid **8** (1.57 g, 82%): $^1H$ NMR (400 MHz, $CDCl_3$) δ 7.32-7.25 (m, 5H), 4.54 (s, 2H), 3.98-3.82 (m, 9H), 3.64-3.50 (m, 14H), 3.39-3.38 (m, 1H), 3.26-3.24 (m, 1H), 2.49-1.04 (m, 25H), 1.27 (s, 3H), 1.00 (s, 3H), 0.68 (s, 3H); $^{13}C$ NMR (100 MHz, $CDCl_3$) δ 138.2, 128.3 (2 x C), 127.7 (2 x C), 127.6, 111.7, 110.4, 78.2, 77.5, 76.5, 73.2, 70.6 (4 x C), 70.5, 70.2, 69.9, 69.4, 66.7, 65.1, 64.1, 64.0, 63.2, 58.0, 55.1, 45.2, 44.6, 42.5, 42.0, 37.1, 36.2, 35.9, 34.3, 31.1, 27.1, 26.3, 24.5, 23.9, 23.1, 20.4, 20.0, 14.0.s

**17β-(2-Methyl-1,3-dioxolan-2-yl)-11α-((1-phenyl-2,5,8,11-tetraoxaheptadecan-17-yl)oxy)-spiro[5β-androstane-3,2'-[1,3]dioxolane] (9).** To a stirred solution of steroid **8** (1.57 g, 2.1 mmol) in EtOAc (30 mL) was added Pd/C (10%, 150 mg) in a Parr hydrogenation flask and hydrogenaton was carried out overnight at 55 psi $H_2$. The mixture was filtered through celite and washed with EtOAc. The solvent was removed to give steroid **9** (1.57 g, 100%): $^1H$ NMR (400 MHz, $CDCl_3$) δ 7.33-7.32 (m, 5H), 4.55 (s, 2H), 4.00-3.83 (m, 9H), 3.65-3.33 (m, 17H), 3.16-3.14 (m,1H), 2.49-2.47 (m, 1H), 2.16-2.13 (m, 1H), 2.00-1.03 (m, 25H), 1.28 (s, 3H), 1.01 (s, 3H), 0.69 (s, 3H); $^{13}C$

NMR (100 MHz, $CDCl_3$) δ 138.2, 128.3 (2 x C), 127.7 (2 x C), 127.6, 111.7, 110.4, 76.2, 73.2, 71.5, 70.6 (4 x C), 70.0, 69.4, 68.0, 65.1, 64.1, 64.0, 63.2, 58.0, 55.1, 45.3, 44.6, 42.6, 42.0, 37.0, 36.2, 35.9, 34.4, 31.1, 30.3, 29.6, 27.2, 26.3, 26.2, 26.0, 24.5, 24.0, 23.9, 23.0, 14.1

**2-(2-(2-((6-((17β-(2-Methyl-1,3-dioxolan-2-yl)-spiro[5β-androstane-3,2'-[1,3]dioxolan]-11-yl)oxy)hexyl)oxy)ethoxy)ethoxy)ethan-1-ol (10).** To a solution of anhydrous liquid ammonium (50 mL) was added sodium metal (230 mg) at -78 °C. After stirring for 15 min, steroid **9** (1.57 g, 2.12 mmol) in THF (10 mL + 2 mL + 2 mL) was added dropwise. After 1 h, solid $NH_4Cl$ (2.0 g) was added. The ammonia was allowed to evaporate at 23 °C. Water was added and the product was extracted into EtOAc (100mL x 3). The combined organic layers were dried over anhydrous $Na_2SO_4$, filtered, the solvent removed and the residue purified by flash column chromatography (silica gel, eluted with 3-5% MeOH in $CH_2Cl_2$) to give steroid **10** (1.3 g, 94%): $^1H$ NMR (400 MHz, $CDCl_3$) δ 4.11-3.91 (m, 8H), 3.88-3.32 (m, 16H), 3.17-3.13 (m, 1H), 2.49-2.45 (m, 1H), 2.16-2.13 (m, 1H), 2.02-1.03 (m, 28H), 1.28 (s, 3H), 1.00 (s, 3H), 0.69 (s, 3H); $^{13}C$ NMR (100 MHz, $CDCl_3$) δ 111.7, 110.4, 76.2, 72.5, 71.5, 70.62, 70.59, 70.4, 70.0, 67.9, 65.1, 64.1, 64.0, 63.2, 61.7, 58.0, 55.1, 45.3, 44.6, 42.5, 42.0, 37.0, 36.2, 35.9, 34.4, 31.2, 30.3, 29.5, 27.2, 26.3, 26.1, 26.0, 24.5, 23.94, 23.89, 23.0, 14.1.

**2-(2-(2-((6-((17β-(2-Methyl-1,3-dioxolan-2-yl)-spiro[5β-androstane-3,2'-[1,3]dioxolan]-11α-yl)oxy)hexyl)oxy)ethoxy)ethoxy)-acetaldehyde (11).** To a stirred solution of the oxalyl chloride (2 mL) in $CH_2Cl_2$ (10 mL) was added DMSO (2.6 mL) in $CH_2Cl_2$ (10 mL) at -78° C. After 10 min, steroid **10** (1.3 mg, 2 mmol) in $CH_2Cl_2$ (3 mL + 2 mL + 1 mL) was added. After 2 h, $Et_3N$ (10 mL) was added and the reaction was allowed to warm to 23 °C and stirring continued for 30 min. The product was extracted into $CH_2Cl_2$ (100 mL). The combined extracts were dried over anhydrous $Na_2SO_4$, filtered, the solvent removed and the residue purified by flash column chromatography (silica gel, eluted with 20-50% EtOAc in hexanes) to give steroid **11** (980 mg, 75%): $^1H$ NMR (400 MHz, $CDCl_3$) δ 9.7 (s, 1H), 4.15 (s, 2H), 4.00-3.32 (m, 20H), 3.16-3.14 (m, 1H), 2.50-2.46 (m, 1H), 2.16-2.13 (m, 1H), 2.02-1.03 (m, 27H), 1.28 (s, 3H), 1.01 (s, 3H), 0.69 (s, 3H); $^{13}C$ NMR (100 MHz, $CDCl_3$) δ 200.9, 111.7, 110.4, 76.8, 76.2, 71.4, 71.2, 70.74, 70.69, 70.0, 67.9, 65.1, 64.1, 64.0, 63.2, 58.0, 55.1, 45.3, 44.6, 42.5, 42.0, 37.0, 36.2, 35.9, 34.4, 31.2, 30.3, 29.6, 27.2, 26.3, 26.1, 26.0, 24.5, 23.94, 23.88, 23.0, 14.1.

**17β-(2-Methyl-1,3-dioxolan-2-yl)-11α-((6-(2-(2-(prop-2-yn-1-yloxy)ethoxy)ethoxy)hexyl)oxy)-spiro[5β-androstane-3,2'-[1,3]dioxolane] (12).** To a stirred solution of steroid **11** (980 mg, 1.5 mmol) in MeOH/THF (10 mL/10 mL) was added dimethyl (1-diazo-2-oxopropyl) phosphonate (600 mg, 3 mmol) and $K_2CO_3$ (2 g, fine powder) at 23 °C. After 16 h, water was added and the product was extracted into EtOAc (100mL x 3). The combined extracts were dried over anhydrous $Na_2SO_4$, filtered, the solvent removed and the residue purified by flash column chromatography (silica gel, eluted with 20-50% EtOAc in hexanes) to give steroid **12** (750 mg, 77%): $^1$H NMR (400 MHz, $CDCl_3$) δ 4.15 (s, 2H), 4.07-3.30 (m, 18H), 3.13-3.11 (m, 1H), 2.47-2.40 (m, 2H), 2.13-2.09 (m, 1H), 1.99-0.99 (m, 29H), 1.25 (s, 3H), 0.98 (s, 3H), 0.66 (s, 3H); $^{13}$C NMR (100 MHz, $CDCl_3$) δ 111.7, 110.4, 79.6, 76.2, 74.5, 71.4, 70.6, 70.4, 70.0, 69.1, 67.9, 65.1, 64.1, 64.0, 63.1, 58.4, 58.0, 55.1, 45.3, 44.6, 42.5, 42.0, 37.0, 36.2, 35.9, 34.4, 31.1, 30.3, 29.5, 27.2, 26.3, 26.1, 26.0, 24.4, 23.92, 23.87, 23.0, 14.0.

**11α-((6-(2-(2-(Prop-2-yn-1-yloxy)ethoxy)ethoxy)hexyl)oxy)-5β-pregnane-3,20-dione (13).** To a stirred solution of steroid **12** (750 mg, 1.16 mmol) in THF (20 mL) was added 6 N HCl (5 mL). After 20 h, the product was extracted into EtOAc (20 mL x 3). The combined extracts were dried over anhydrous $Na_2SO_4$, filtered, the solvent removed and the residue purified by flash column chromatography (silica gel, eluted with 50-100% EtOAc in hexanes) to give steroid **13** (600 mg, 92%): $^1$H NMR (400 MHz, $CDCl_3$) δ 4.15 (s, 2H), 3.64-3.37 (m, 12H), 3.21-3.17 (m, 1H), 2.69-1.12 (m, 30H), 2.09 (s, 3H), 1.05 (s, 3H), 0.56 (s, 3H); $^{13}$C NMR (100 MHz, $CDCl_3$) δ 214.0, 209.1, 79.6, 76.0, 74.5, 71.3, 70.6, 70.4, 70.0, 69.1, 68.3, 63.3, 58.4, 55.3, 45.8, 45.2, 44.4, 43.7, 42.6, 39.8, 38.6, 36.0, 34.6, 31.6, 30.2, 29.5, 26.8, 26.1, 26.0, 25.9, 24.4, 23.4, 23.0, 14.3.

**3α-Hydroxy-11α-((6-(2-(2-(prop-2-yn-1-yloxy)ethoxy)ethoxy)hexyl)oxy)-5β-pregnan-20-one (14, YX85).** To a stirred solution of steroid **13** (600 mg, 1.07 mmol) in THF (20 mL) was added dropwise lithium tri-*tert*-butoxyaluminum hydride (1.5 mL, 1.0 M in THF, 1.5 mmol) at - 40 °C. After 4 h, 6 N HCl (2 mL) was added, the reaction allowed to warm to 23 °C and stirring was continued overnight. The product was extracted into EtOAc (50 mL x 3). The combined extracts were washed with water (100 mL), dried over anhydrous $Na_2SO_4$, filtered, the solvent removed and the residue purified by flash column chromatography (silica gel, eluted with 50-100% EtOAc in hexanes) to give steroid **YX85** (**14**, 500 mg, 80%): $^1$H NMR (400 MHz, $CDCl_3$) δ 4.13-4.12 (m, 2H), 3.62-3.27 (m, 13H), 3.16-3.10 (m, 1H), 2.50-2.38 (m, 3H), 2.18-0.83 (m, 28H), 2.06 (s, 3H),

0.92 (s, 3H), 0.49 (s, 3H); $^{13}C$ NMR (100 MHz, $CDCl_3$) δ 209.4, 79.6, 76.3, 74.6, 71.8, 71.3, 70.6, 70.3, 70.0, 69.0, 68.1, 63.4, 58.3, 55.4, 45.2, 44.6, 43.8, 43.6, 38.1, 36.8, 35.8, 34.9, 31.6, 31.5, 30.2, 29.5, 27.5, 26.5, 26.1, 25.9, 24.5, 24.1, 23.0, 14.3.

**11α-(2-Propenyloxy)-5β-pregnan-3,20-dione (15).** To a stirred solution of steroid **4** (300 mg, 0.65 mmol) in acetone (20 mL) was added PTSA (50 mg in 5 mL acetone) at 23 °C. The reaction was stirred for 16 h. Solid $NaHCO_3$ was added and after 10 min the acetone was removed. Water (20 mL) was added to the residue and the product was extracted into EtOAc (100 mL). The extract was dried over anhydrous $Na_2SO_4$, filtered and the solvent removed. The residue was purified by flash column chromatography (silica gel, eluted with 10% EtOAc in hexanes) to give steroid **15** (200 mg, 83%): $^1H$ NMR (400 MHz, $CDCl_3$) δ 5.84-5.77 (m, 1H), 5.18-5.14 (m, 1H), 5.07-5.04 (m, 1H), 4.09-4.05 (m, 1H), 3.81-3.77 (m, 1H), 3.56-3.49 (m, 1H), 2.72-1.10 (m, 21H), 2.09 (s, 3H), 1.06 (s, 3H), 0.58 (s, 3H); $^{13}C$ NMR (100 MHz, $CDCl_3$) δ 213.7, 208.9, 134.6, 116.1, 75.9, 69.2, 63.1, 55.2, 45.7, 45.1, 44.2, 43.6, 42.5, 39.6, 38.4, 35.8, 34.4, 31.4, 26.6, 25.7, 24.3, 23.3, 22.9, 14.2.

**3α-Hydroxy-11α-(2-propenyloxy)-5β-pregnan-20-one (16, YX47).** To a stirred solution of steroid **5** (200 mg, 0.54 mmol) in THF (20 mL) was added dropwise lithium tri-*tert*-butoxyaluminum hydride (1M in THF, 1.1 mL, 1.1 mmol) at -40 °C. After 6 h acetone (1 mL) was added and stirring continued for 15 min at -40 °C. Aqueous $NH_4Cl$ was added and stirring was continued over the weekend at 23 °C. The product was extracted into $CH_2Cl_2$ (50 mL × 3). The combined extracts were dried over anhydrous $Na_2SO_4$, filtered, solvent removed and the residue purified by flash column chromatography (silica gel, eluted with 25% EtOAc in hexanes) to give **YX47** (**16**,170 mg, 84%): $^1H$ NMR (400 MHz, $CDCl_3$) δ 5.86-5.79 (m, 1H), 5.21-5.17 (m, 1H), 5.06-5.03 (m, 1H), 4.08-4.04 (m, 1H), 3.80-3.76 (m, 1H), 3.75-3.46 (m, 1H), 3.45-3.40 (m, 1H), 2.53-2.42 (m, 2H), 2.25-0.81 (m, 20H), 2.08 (s, 3H), 0.97 (s, 3H), 0.55 (s, 3H); $^{13}C$ NMR (100 MHz, $CDCl_3$) δ 209.1, 135.0, 115.6, 76.3, 71.7, 69.0, 63.3, 55.3, 45.2, 44.5, 43.7, 43.6, 38.0, 36.8, 35.7, 34.9, 31.5, 31.4, 27.4, 26.4, 24.3, 24.0, 22.9, 14.2.

**Synthesis of MQ385, MQ390 and MQ392**

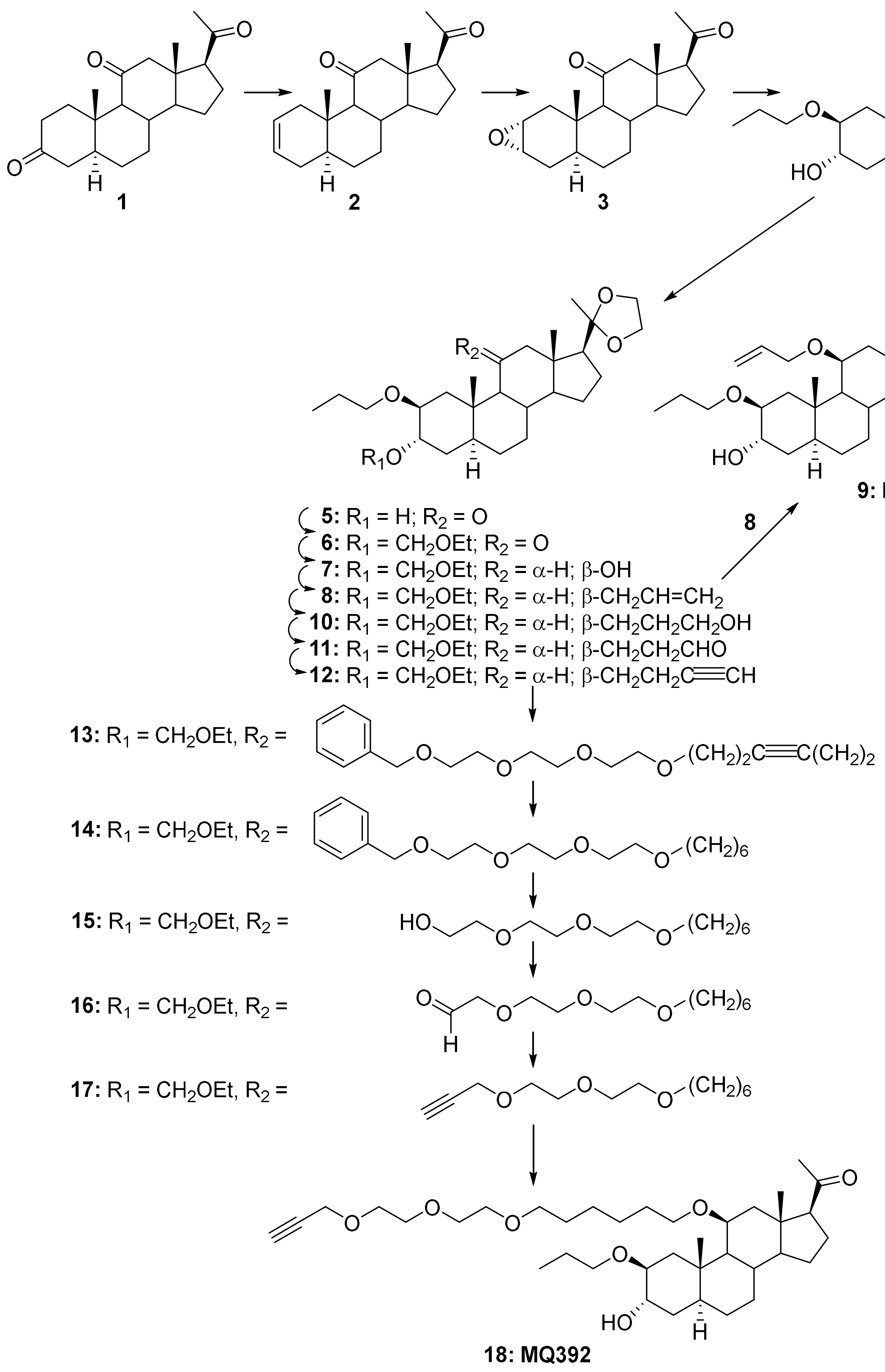

**5α-Pregn-2-en-11,20-dione (2).** To a stirred suspension of zinc dust (12.0 g, 184 mmol) and TMSCl (11.7 mL, 92 mmol) in THF (180 mL) was added 5α-pregnane-3,11,20-trione (**1**, 3.03 g, 9.2 mmol) at 23 °C. The mixture was refluxed for 16 h. After cooling, the mixture was filtered through Celite and washed with THF (50 mL). The filtrate was washed with aqueous $NaHCO_3$, dried over anhydrous $Na_2SO_4$, filtered and the solvent removed. The residue was purified by flash column chromatography (silica gel, eluted with 5% EtOAc in hexanes) to give steroid **2** containing the isomeric $\Delta^3$ steroid (1.67 g, $\Delta^2/\Delta^3$ = 7/1, 58%). Steroid **2** has: $^1$H NMR (400 MHz, $CDCl_3$) δ 5.56-5.55 (m, 2H), 2.79-2.18 (m, 5H), 2.08 (s, 3H), 1.84-0.55 (m, 14H), 0.98 (s, 3H), 0.56 (s, 3H); $^{13}$C NMR (100 MHz, $CDCl_3$) δ 209.6, 208.0, 125.9, 125.0, 64.0, 62.1, 56.6, 55.4, 46.9, 41.1, 38.8, 36.6, 34.2, 32.4, 31.2, 29.5, 28.0, 23.8, 23.1, 14.1, 11.4.

**2α,3α-Epoxy-5α-pregnane-11,20-dione (3).** To a stirred solution of the $\Delta^2$ steroid **2** containing the isomeric $\Delta^3$ steroid (1.67 g, 5.32 mmol) in $CH_2Cl_2$ (40 mL) was added formic acid (10 mL) and hydrogen peroxide (10 mL) at 23 °C. After 4 h, the product was extracted into $CH_2Cl_2$ (200 mL). The extract was washed with $NaHCO_3$ (50 mL), dried over anhydrous $Na_2SO_4$, filtered and the solvent removed. The residue was purified by flash column chromatography (silica gel, eluted with 10% EtOAc in hexanes) to give steroid **3** (1.20 g, 68%): $^1$H NMR (400 MHz, $CDCl_3$) δ 3.14-3.08 (m, 2H), 2.83-2.19 (m, 5H), 2.08 (s, 3H), 1.87-0.98 (m, 14H), 0.96 (s, 3H), 0.55 (s, 3H); $^{13}$C NMR (100 MHz, $CDCl_3$) δ 209.5, 208.0, 63.9, 62.1, 56.6, 55.3, 52.1, 50.8, 46.9, 37.2, 36.7, 36.1, 33.3, 32.2, 31.3, 28.5, 27.7, 23.9, 23.2, 14.2, 12.6.

**3α-Hydroxy-2β-propoxy-5α-pregnane-11,20-dione (4).** To a stirred solution of steroid **3** (1.23 g, 3.94 mmol) in 1-propanol (30 mL) was added concentrated $H_2SO_4$ (3 mL) at 0° C. After addition, the reaction was warmed up to 23 °C for 1 h. Water was added and then solid $NaHCO_3$ was slowly added until $CO_2$ gas evolution stopped. Most of the 1-propanol was removed, water was added and the product was extracted into EtOAc (200 mL x2). The combined extracts were dried over anhydrous $Na_2SO_4$, filtered and the solvent removed. The residue was purified by flash column chromatography (silica gel, eluted with 50% EtOAc in hexanes to 100% EtOAc) to give steroid **4** (901mg, 63%): $^1$H NMR (400 MHz, $CDCl_3$) δ 3.88-3.87 (m, 1H), 3.60-3.54 (m, 1H), 3.26 (s, 1H), 3.24-3.20 (m, 1H), 2.69-2.17 (m, 5H), 2.07 (s, 3H), 1.85-1.12 (m, 17H), 1.10 (s, 3H), 0.91 (t, $J$ = 7.6 Hz, 3H), 0.53 (s, 3H); $^{13}$C NMR (100 MHz, $CDCl_3$) δ 209.8, 208.3, 78.6, 70.4, 68.6, 64.8, 62.1, 56.6, 55.6, 47.1, 38.7, 35.9, 35.6, 34.4, 32.5, 31.6, 31.3, 27.4, 23.8, 23.2, 23.1, 14.2, 12.9, 10.7.

**3α-Hydroxy-17β-(2-methyl-1,3-dioxolan-2-yl)-2β-propoxy-5α-androstan-11-one (5).** To a stirred solution of steroid **4** (901 mg, 2.31 mmol) in benzene (150 mL) was added ethylene glycol (2 mL) and PTSA (50 mg) at 23 °C. The reaction was refluxed in a flask equipped with a Dean-Stark apparatus under $N_2$ for 16 h. After cooling, solid $NaHCO_3$ (300 mg) was added and stirring continued for 10 min. Aqueous $NaHCO_3$ (100 mL) was added and the product was extracted into EtOAc (300 mL). The extract was washed with brine (100 mL x3), dried over anhydrous $Na_2SO_4$, filtered and the solvent removed. The residue was purified by flash column chromatography (silica gel, eluted with 25% EtOAc in hexanes) to give steroid **5** (964 mg, 96%): $^1$H NMR (400 MHz, $CDCl_3$) δ 3.96-3.86 (m, 5H), 3.84-3.83 (m, 1H), 3.58 (s, 1H), 3.31-3.21 (m, 1H), 2.69-2.19 (m, 3H), 2.01-1.07 (m, 19H), 1.21 (s, 3H), 1.07 (s, 3H), 0.90 (t, *J* = 7.6 Hz, 3H), 0.66 (s, 3H); $^{13}$C NMR (100 MHz, $CDCl_3$) δ 211.7, 111.2, 78.7, 70.3, 68.7, 64.8, 64.7, 63.1, 57.7, 56.8, 55.3, 45.9, 38.7, 35.6, 35.5, 34.4, 32.4, 31.7, 27.5, 24.2, 23.4, 23.2, 23.1, 14.0, 12.8, 10.7.

**3α-(Ethoxymethoxy)-17β-(2-methyl-1,3-dioxolan-2-yl)-2β-propoxy-5α-androstan-11-one (6).** To a stirred solution of steroid **5** (964 mg, 2.22 mmol) in $CH_2Cl_2$ (20 mL) was added chloromethyl ethyl ether (0.31 mL, 3,3 mmol) and (*i*-Pr)$_2$NEt (0.87 mL, 5 mmol) at 23 °C. After 16 h, solvent was removed and the residue was purified by flash column chromatography (silica gel, eluted with 10-20% EtOAc in hexanes) to give steroid **6** (1.03 mg, 94%): $^1$H NMR (400 MHz, $CDCl_3$) δ 4.69 (s, 2H), 3.98-3.83 (m, 4H), 3.72 (s, 1H), 3.60-3.54 (m 3H), 3.40 (s, 1H), 3.27-3.25 (m, 1H), 2.76-2.20 (m, 3H), 2.03-1.02 (m, 21H), 1.24 (s, 3H), 1.14 (s, 3H), 1.02 (t, *J* = 7.6 Hz, 3H), 0.69 (s, 3H); $^{13}$C NMR (100 MHz, $CDCl_3$) δ 211.7, 111.2, 93.9, 74.2, 70.4, 64.9, 64.8, 63.2, 63.1, 57.8, 56.9, 55.5, 45.9, 39.5 (2 x C), 35.7, 35.4, 34.9, 32.5, 29.6, 27.6, 24.3, 23.5, 23.3, 23.2, 15.1, 14.1, 12.9, 10.8.

**3α-(Ethoxymethoxy)-17β-(2-methyl-1,3-dioxolan-2-yl)-2β-propoxy-5α-androstan-11β-ol (7).** To a stirred solution of steroid **6** (1.03 g, 2.09 mmol) in anhydrous diethyl ether (60 mL) was added $LiAlH_4$ (2.4 M in THF, 5 mL, 12 mmol) at 23 °C. After 2 h, water (0.46 mL), 10% NaOH (0.92 mL), and water (1.38 mL) were slowly added sequentially and stirring continued for 1 h. The mixture was filtered through Celite and washed with EtOAc (200 mL). The EtOAc was removed and the residue was purified by flash column chromatography (silica gel, eluted with 40% EtOAc in hexanes) to give steroid **7** (1.01 g, 98%): $^1$H NMR (400 MHz, $CDCl_3$) δ 4.68 (s, 2H), 4.25 (s, 1H), 3.98-3.82 (m, 4H), 3.75 (s, 1H), 3.75-3.27 (m 5H), 2.19-0.75 (m, 25H), 1.24 (s, 3H), 1.11 (s,

3H), 0.95 (s, 3H), 0.93 (t, $J$ = 7.6, 3H); $^{13}$C NMR (100 MHz, $CDCl_3$) δ 111.8, 93.7, 73.6, 70.6, 68.2, 64.9, 63.1, 63.0, 58.8, 58.6, 57.9, 48.3, 41.2, 40.5, 36.3, 35.7 (2 x C), 32.1, 30.1, 29.0, 27.6, 24.4, 23.6, 23.3, 22.6, 16.6, 15.7, 15.1, 10.7.

**2-(3α-(Ethoxymethoxy)-11β-(2-propenyloxy)-2β-propoxy-5α-androstan-17β-yl)-2-methyl-1,3-dioxolane (8).** To a stirred solution of steroid **7** (1.10 g, 2.05 mmol) in THF (30 mL) was added KH in mineral oil (30%, 0.67 g, 5 mmol) at 23 °C. After 30 min, allyl bromide (0.87 mL, 10 mmol) was added and the mixture was refluxed for 16 h. After cooling, water was added and the product was extracted into EtOAc (150 mL x 2). The combined extracts were dried over anhydrous $Na_2SO_4$, filtered and the solvent removed. The residue was purified by flash column chromatography (silica gel, eluted with 25-40% EtOAc in hexanes) to give steroid **8** (966 mg, 88%): $^1$H NMR (400 MHz, $CDCl_3$) δ 5.89-5.85 (m, 1H), 5.22 (dd, $J$ = 10.8 Hz, 2H), 4.65 (s, 2H), 4.08-3.22 (m, 13H), 2.46-2.43 (m, 1H), 1.88-0.71 (m, 26H), 1.23 (s, 3H), 1.08 (s, 3H), 0.90 (s, 3H); $^{13}$C NMR (100 MHz, $CDCl_3$) δ 135.5, 135.4, 115.3, 111.8, 93.6, 75.1, 74.0, 70.2, 69.2, 64.9, 63.1, 62.9, 58.7, 58.4, 58.0, 41.4, 40.6, 40.4, 35.7, 35.6, 32.3, 30.7, 29.1, 27.5, 24.3, 23.5, 23.1, 22.5, 16.0, 15.0, 14.2, 10.6.

**3α-Hydroxy-11β-(2-propenyloxy)-2-propoxy-5α-pregnan-20-one (9, MQ390).** To a solution of steroid **8** (50 mg, 0.094 mmol) in THF (10 mL) was added 6 N HCl (10 mL) at 23 °C. After 2 h, the product was extracted into EtOAc (100 mL x2). The combined extracts were washed with aqueous $NaHCO_3$ (50 ml x2), dried over anhydrous $Na_2SO_4$, filtered and the solvent removed. The residue was purified by flash column chromatography (silica gel, eluted with 25-50% EtOAc in hexanes) to give **MQ390** (**9**, 36 mg, 89%): $^1$H NMR (400 MHz, $CDCl_3$) δ 5.95-5.89 (m, 1H), 5.22 (dd, $J$ = 10.8 Hz, 2H), 4.10-4.08 (m, 1H), 3.92-3.86 (m, 2H), 3.78-3.74 (m, 1H), 3.49-3.43 (m, 2H), 3.28-3.27 (m, 1H), 2.47-2.43 (m, 2H), 2.18-1.07 (m, 20H), 2.10 (s, 3H), 1.12 (s, 3H), 0.92 (t, $J$ = 7.6 Hz, 3H), 0.79 (s, 3H); $^{13}$C NMR (100 MHz, $CDCl_3$) δ 209.7, 135.2, 115.9, 78.54, 78.51, 74.9, 70.5, 69.5, 64.1, 58.8, 58.3, 43.6, 40.2, 39.9, 36.2, 35.1, 32.5, 31.6, 31.4, 27.5, 27.3, 24.3, 23.2, 22.6, 16.1, 14.7, 10.8.

**3-((3α-(Ethoxymethoxy)-17β-(2-methyl-1,3-dioxolan-2-yl)-2-propoxy-androstan-11-yl)oxy)propan-1-ol (10).** To a stirred solution of steroid **8** (916 mg, 1.72 mmol) in THF (30 mL) was added 9-BBN (0.5 M in THF, 10 mL, 5 mmol) at 23 °C. After 16 h, 3 N NaOH (30 mL, 90 mmol) and $H_2O_2$ (10 mL) were added and stirring continued at 23 °C for 1 h. The product was extracted into EtOAc (150 mL x 3). The combined extracts were washed with brine (100 mL x3), dried over anhydrous $Na_2SO_4$, filtered and the solvent removed. The residue was purified by flash column chromatography (silica gel, eluted with 25-50% EtOAc in hexanes) to give steroid **10** (903 mg, 95%): $^1$H NMR (400 MHz, $CDCl_3$) δ 4.71 (s, 2H), 3.99-3.82 (m, 4H), 3.77-3.29 (m, 11H), 2.51-2.47 (m, 1H), 2.11-0.76 (m, 29H), 1.26 (s, 3H), 1.21 (t, $J$ = 7.2 Hz, 3H), 1.13 (s, 3H); $^{13}$C NMR (100 MHz, $CDCl_3$) δ 111.9, 93.8, 76.7, 73.9, 70.6, 67.0, 65.0, 63.2, 61.8, 58.7, 58.0, 41.5, 40.7, 36.2, 35.7, 34.6, 32.7, 32.3, 30.7, 29.1, 27.6, 27.3, 25.1, 24.4, 23.5, 23.2, 22.6, 22.59, 16.4, 15.1, 14.5, 10.8.

**3-((3α-(Ethoxymethoxy)-17β-(2-methyl-1,3-dioxolan-2-yl)-2-propoxy-androstan-11-yl)oxy)propan-1-al (11).** To a stirred solution of oxalyl chloride (0.25 mL, 3 mmol) in $CH_2Cl_2$ (15 mL) was added DMSO (0.25 mL, 3.5 mmol) in $CH_2Cl_2$ (2 mL) at -78 °C and stirring was continued for 15 min. Steroid **10** (903 mg, 1.64 mmol) in $CH_2Cl_2$ (10 mL) was added and stirring was continued for 1 h at -78 °C. $Et_3N$ (0.7 mL, 5 mmol) was added and stirring was continued for 30 min. The reaction was warmed to 23 °C and water was added. The product was extracted into $CH_2Cl_2$ (100 mL x 3). The combined extracts were dried over anhydrous $Na_2SO_4$, filtered and the solvent removed. The residue was purified by flash column chromatography (silica gel, eluted with 25-40% EtOAc in hexanes) to give steroid **11** (720 mg, 80%): $^1$H NMR (400 MHz, $CDCl_3$) δ 9.75 (s, 1H), 4.66 (s, 2H), 3.93-3.24 (m, 13H), 2.56-0.69 (m, 32H), 1.22 (s, 3H), 0.94 (s, 3H); $^{13}$C NMR (100 MHz, $CDCl_3$) δ 201.6, 111.7, 93.7, 76.0, 73.7, 70.4, 64.9, 63.1, 62.9, 62.0, 58.6, 58.3, 57.9, 44.0, 41.4, 40.5, 40.4, 36.1, 35.7, 32.2, 30.6, 29.0, 27.5, 25.5, 24.3, 23.5, 23.2, 22.5, 16.2, 15.0, 14.2, 10.7.

**11β-(But-3-yn-1-yloxy)-3α-(ethoxymethoxy)-17β-(2-methyl-1,3-dioxolan-2-yl)-2β-propoxy-5α-androstane (12).** To a stirred solution of steroid **11** (720 mg, 1.31 mmol) in THF/MeOH (10 mL/10 mL) was added dimethyl-1-diazo-oxopropyl phosphonate (0.62 mL, 4 mmol) and $K_2CO_3$ (1.1 g, 8 mmol, fine powder) at 23 °C. After 16 h, water was added and the product was extracted into EtOAc (100 mL x3). The combined extracts were dried over anhydrous $Na_2SO_4$, filtered and the solvent removed. The residue was purified by flash column chromatography (silica gel, eluted with 25-35% EtOAc in hexanes) to give steroid **12** (480 mg, 67%): $^1$H NMR (400 MHz, $CDCl_3$) δ 4.65 (s, 2H), 3.95-3.25 (m, 13H), 2.44-0.69 (m, 33H), 1.24 (s, 3H), 1.09 (s, 3H); $^{13}$C NMR (100 MHz, $CDCl_3$) δ 111.8, 93.7, 81.8, 76.7, 75.7, 73.8, 70.4, 69.0, 66.5, 65.0, 63.1, 63.0, 58.7, 58.4,

58.0, 41.4, 40.7, 40.5, 36.0, 35.7, 32.3, 30.7, 29.1, 27.6, 24.4, 23.5, 23.2, 22.6, 20.0, 16.1, 15.0, 14.2, 10.7.

**3α-(Ethoxymethoxy)-17β-(2-methyl-1,3-dioxolan-2-yl)-11β-((1-phenyl-2,5,8,11-tetraoxaheptadec-14-yn-17-yl)oxy)-2β-propoxy-5α-androstane (13).** To a mixture of steroid **12** (480 mg, 0.88 mmol), 13-iodo-1-phenyl-2,5,8,11-tetraoxatridecane (591 mg, 1.5 mmol), bis[(2-dimethylamino)phenyl]amine nickel(ii) chloride (52 mg, 0.15 mmol), Cu(I)I (38 mg, 0.2 mmol) and $Cs_2CO_3$ (457 mg, 1.4 mmol) was added anhydrous dioxane (25 mL) at 23 °C. The stirred reaction was heated to 140 °C for 16 h. After cooling, dioxane was removed under reduced pressure and the residue was purified by flash column chromatography (silica gel, eluted with 25-60% EtOAc in hexanes) to give steroid **13** (520 mg, 73%): $^1$H NMR (400 MHz, $CDCl_3$) δ 7.32-7.26 (m, 5H), 4.68 (s, 2H), 4.54 (s, 2H), 3.96-3.26 (m, 27H), 2.45-0.71 (m, 34H), 1.25 (s, 3H), 1.06 (s, 3H); $^{13}$C NMR (100 MHz, $CDCl_3$) δ 138.1, 128.2 (2 x C), 127.6 (2 x C), 127.5, 111.9, 93.9, 78.3, 75.5, 73.9, 73.1, 70.5 (3 x C), 70.46, 70.1, 69.8, 69.3, 66.9, 65.0, 63.2, 63.0, 58.7, 58.4, 58.0, 41.4, 40.8 (2 x C), 40.5, 36.1, 35.7, 34.4, 32.3, 30.7, 29.1, 27.6, 24.4, 23.5, 23.2, 22.5, 20.4, 19.9, 16.1, 15.1, 14.3, 10.8.

**3α-(Ethoxymethoxy)-17β-(2-methyl-1,3-dioxolan-2-yl)-11β-((1-phenyl-2,5,8,11-tetraoxaheptadecan-17-yl)oxy)-2β-propoxy-5α-androstane (14).** To a solution of steroid **13** (520 mg, 0.64mmol) in EtOAc (60 mL) in a Parr hydrogenation flask was added Pd/C (300 mg) at 23 °C. The flask was evacuated and charged with $H_2$ three times. Hydrogenation was carried out 55 psi $H_2$ overnight. The mixture was filtered through Celite and washed with EtOAc (100 mL). The solvent was removed and the residue was purified by flash column chromatography (silica gel, eluted with 25-50% EtOAc in hexanes) to give steroid **14** (503 mg, 97%): $^1$H NMR (400 MHz, $CDCl_3$) δ 7.32-7.26 (m 5H), 4.69 (s, 2H), 4.55 (s, 2H), 3.98-3.08 (m, 27H), 2.46-2.43 (m, 1H), 1.86-0.71 (m, 37H), 1.28 (s, 3H), 1.12 (s, 3H); $^{13}$C NMR (100 MHz, $CDCl_3$) δ 138.2, 128.3 (2 x C), 127.7 (2 x C), 127.6, 112.0, 93.8, 76.8, 75.6, 74.2, 73.2, 71.4, 70.6 (2 x C), 70.4, 70.0, 69.4, 68.3, 65.2, 63.2, 63.1, 58.8, 58.6, 58.1, 41.6, 40.7, 40.6, 35.9 (2 x C), 35.7, 32.5, 30.8, 30.4, 29.5, 29.4, 29.3, 27.7, 26.5, 26.0, 24.5, 23.6, 23.3, 22.6, 16.1, 15.1, 14.3, 10.9.

**3α-(Ethoxymethoxy)-17β-(2-methyl-1,3-dioxolan-2-yl)-11β-((6-(2-(2-(2-hydroxyethoxy) ethoxy)ethoxy)hexyl)oxy)-2β-propoxy-5α-androstane (15).** To a solution of anhydrous liquid ammonia (50 mL) was added sodium metal (230 mg, 10 mmol) at -78 °C. The mixture was stirred for 15 min. Steroid **14** (503 mg, 0.62 mmol) in THF (40 mL) was added within 15 min and stirring was continued for 1 h. Solid $NaHCO_3$ was added and the mixture was warmed to 23 °C allowing the ammonia gas to evaporate. Aqueous $NaHCO_3$ was added. The product was extracted into EtOAc (200 mL x 2), The combined extracts were dried over anhydrous $Na_2SO_4$, filtered and the solvent removed. The residue was purified by flash column chromatography (silica gel, eluted with 25-50% EtOAc in hexanes) to give steroid **15** (320 mg, 67%): $^1$H NMR (400 MHz, $CDCl_3$) δ 4.68 (s, 2H), 3.98-3.06 (m, 27H), 2.77 (br, 1H), 2.45-2.42 (m, 1H), 1.88-0.69 (m, 37H), 1.28 (s, 3H), 1.08 (s, 3H); $^{13}$C NMR (100 MHz, $CDCl_3$) 112.0, 93.8 76.8, 75.6, 74.2, 72.5, 71.5, 70.6, 70.5, 70.4, 70.3, 70.0, 68.3, 65.2, 63.3, 63.1, 61.7, 58.8, 58.6, 58.1, 41.6, 40.7, 40.6, 35.9, 35.7, 32.4, 30.9, 30.4, 29.4, 29.3, 27.7, 26.5, 26.0, 24.5, 23.6, 23.3, 22.6, 16.1, 15.1, 14.3, 10.9.

**3α-(Ethoxymethoxy)-17β-(2-methyl-1,3-dioxolan-2-yl)-11β-((6-(2-(2-(2-oxoethoxy)ethoxy) ethoxy)hexyl)oxy)-2β-propoxy-5α-androstane (16).** To a stirred solution of oxalyl chloride (0.25 mL, 3 mmol) in $CH_2Cl_2$ (15 mL) was added DMSO (0.32 mL, 4.5 mmol) at -78 °C. After 15 min, steroid **15** (320 mg, 0.44 mmol) in $CH_2Cl_2$ (8 mL) was added and stirring was continued at -78 °C for 1 h. $Et_3N$ (0.84 mL, 6 mmol) was added at -78 °C and stirring continued for 30 min, then warmed to 23 °C and stirred an additional 30 min. Water (50 mL) was added and the product was extracted into $CH_2Cl_2$ (150 mL x2 ). The combined extracts were dried over anhydrous $Na_2SO_4$, filtered and the solvent removed. The residue was purified by flash column chromatography (silica gel, eluted with 25-50% EtOAc in hexanes) to give steroid **16** (265 mg, 83%): $^1$H NMR (400 MHz, $CDCl_3$) δ 9.66 (s, 1H), 4.64 (s, 2H), 4.10 (s, 2H), 3.92-3.05 (m, 23H), 2.41-2.38 (m, 1H), 1.85-0.67 (m, 37H), 1.29 (s, 3H), 1.04 (s, 3H); $^{13}$C NMR (100 MHz, $CDCl_3$) δ 200.7, 111.8, 93.6, 76.6, 75.4, 74.0, 71.2, 71.0, 70.6, 70.5, 70.2, 69.8, 68.1, 65.0, 63.1, 62.9, 58.6, 58.4, 57.9, 41.4, 40.5, 40.4, 35.7, 35.6, 32.3, 30.7, 30.2, 29.3, 29.1, 27.5, 26.3, 25.8, 24.3, 23.5, 23.1, 22.4, 15.9, 15.0, 14.1, 10.7.

**3α-(Ethoxymethoxy)-17β-(2-methyl-1,3-dioxolan-2-yl)-2β-propoxy-11β-((6-(2-(2-(prop-2-yn-1-yloxy)ethoxy)ethoxy)hexyl)oxy)-5α-androstane (17).** To a stirred solution of steroid **16** (320 mg, 0.44 mmol) in THF (5 mL) and MeOH (5mL) was added dimethyl-1-diazo-2-oxopropyl phosphonate (0.25 mL, 3 mmol) and $K_2CO_3$ fine powder (552 mg, 4 mmol) at 23 °C. After 16 h, water was added and the product was extracted into EtOAc (100 mL x 2). The combined extracts were dried over anhydrous $Na_2SO_4$, filtered and the solvent removed. The residue was purified by flash column chromatography (silica gel, eluted with 25-50% EtOAc in hexanes) to give steroid **17** (182 mg, 83%): $^{1}$H NMR (400 MHz, $CDCl_3$) δ 4.63 (s, 2H), 4.13 (s, 2H), 3.91-3.06 (m, 23H), 2.41-2.36 (m, 2H), 1.84-0.65 (m, 37H), 1.28 (s, 3H), 1.04 (s, 3H); $^{13}$C NMR (100 MHz, $CDCl_3$) δ 111.8, 93.6, 79.4, 77.0, 75.4, 75.3, 74.4, 74.0, 73.9, 71.2, 70.4, 70.2, 69.8, 68.9, 68.1, 65.0, 63.0, 62.9, 58.6, 58.4, 58.2, 57.9, 41.4, 40.6, 40.4, 35.7, 32.3, 30.7, 30.2, 29.3, 29.1, 27.5, 26.3, 25.8, 24.3, 23.5, 23.1, 22.4, 15.9, 15.0, 14.1, 10.7.

**3α-Hydroxy-2β-propoxy-11β-((6-(2-(2-(prop-2-yn-1-yloxy)ethoxy)ethoxy)hexyl)oxy)-5α-pregnan-20-one (18, MQ392).** To a stirred solution of steroid **17** (236 mg, 0.33 mmol) in THF (10 mL) was added 6 N HCl (10 mL) at 23 °C. After 2 h, solid $NaHCO_3$ was slowly added until pH~8 was reached. The product was extracted into EtOAc (150 mL x 3). The combined extracts were dried over anhydrous $Na_2SO_4$, filtered and the solvent removed. The residue was purified by flash column chromatography (silica gel, eluted with 25-70% EtOAc in hexanes) to give **MQ392** (**18**,195 mg, 96%): $^{1}$H NMR (400 MHz, $CDCl_3$) δ 4.16 (s, 2H), 3.86 (s, 1H), 3.73-3.08 (m, 16H), 2.41-0.74 (m, 34H), 2.07 (s, 3H), 1.05 (s, 3H), 0.71 (s, 3H); $^{13}$C NMR (100 MHz, $CDCl_3$) δ 209.7, 79.4, 78.4, 75.2, 74.5, 74.4, 71.2, 70.4, 70.3, 70.2, 69.8, 68.9, 68.6, 68.5, 68.3, 63.9, 58.5, 58.2, 43.5, 40.0, 39.7, 36.0, 34.9, 32.3, 31.4, 31.3, 30.1, 29.3, 27.4, 26.3, 25.8, 24.1, 23.1, 22.3, 15.8, 14.5, 10.7.

**Synthesis of YX88 and YX89**

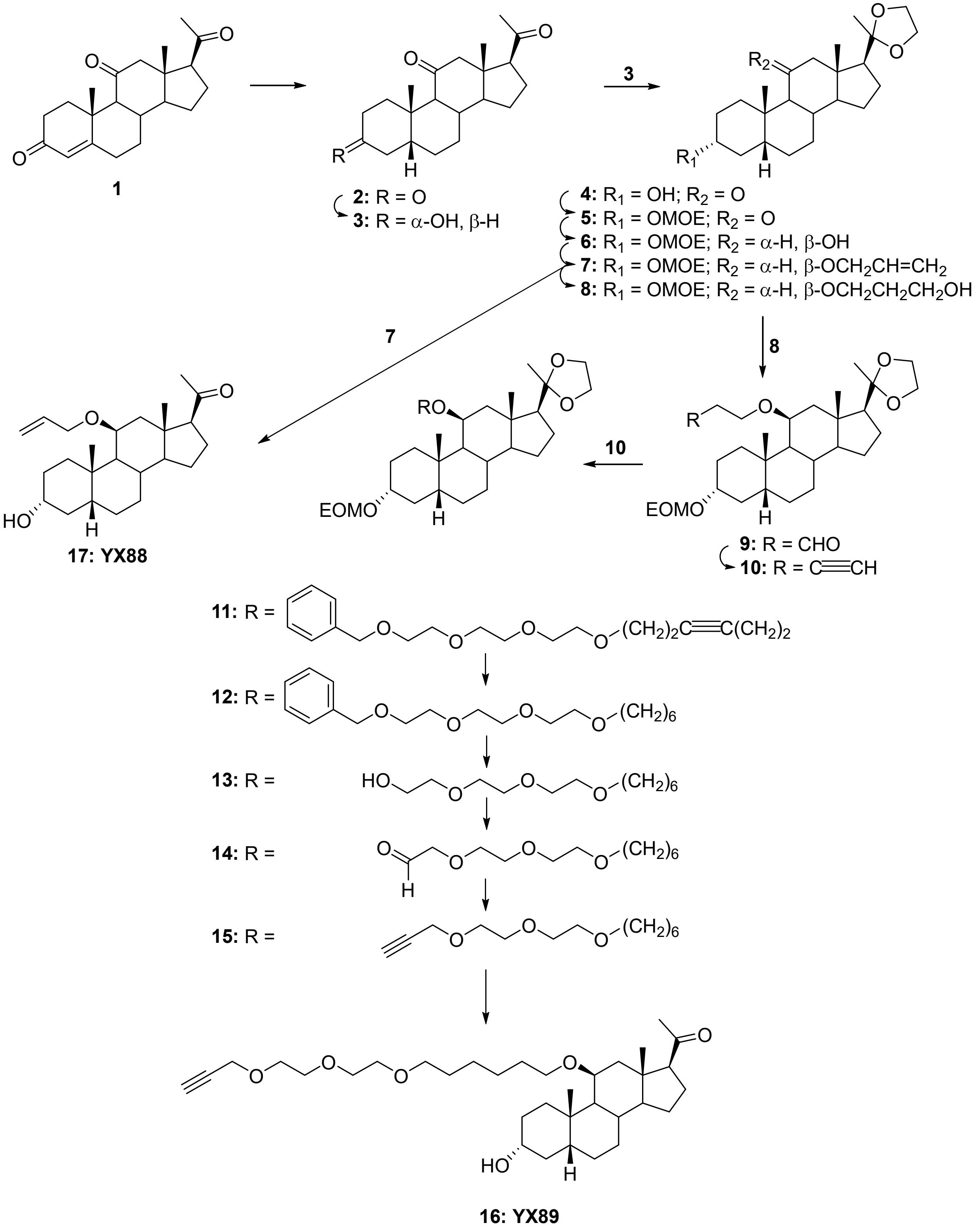

**5β-Pregnane-3,11,20-trione (2).** To a solution of pregn-4-ene-3,11,17-trione (**1**, 5 g, 15.2 mmol) in pyridine (300 mL) was added Pd/$CaCO_3$ (10%, 200 mg) in a Parr hydrogenation flask. Hydrogenation was carried out at 23 °C overnight under $H_2$ at 55 psi. The mixture was filtered through celite and the celite was washed with $CH_2Cl_2$. The solvent was removed and the residue was purified by flash column chromatography (silica gel, eluted with 50% EtOAc in hexanes) to give steroid **2** (4.5 g, 90%): $^1$H NMR (400 MHz, $CDCl_3$) δ 2.78-2.69 (m, 2H), 2.60-2.57 (m, 1H), 2.50-2.40 (m, 2H), 3.30-1.15 (m, 16H), 2.08 (s, 3H), 1.19 (s, 3H), 0.58 (s, 3H).

**3α-Hydroxy-5β-pregnane-11,20-dione (3).** To a stirred solution of steroid **2** (3.7 g, 11.2 mmol) in THF (200 mL) was added lithium tri-*tert*-butoxyaluminum hydride (22 mL, 22 mmol, 1.0 M in THF) at -40 °C. After 2 h, water was added and the reaction was allowed to warm to 23 °C. After 16 h, the product was extracted into $CH_2Cl_2$ (100 mL x 3). The combined extracts were washed with water (100 mL), dried over anhydrous $Na_2SO_4$, filtered, the solvent removed and the residue purified by flash column chromatography (silica gel, eluted with 20% EtOAc in hexanes) to give steroid **3** (2.90 g, 78 %): $^1$H NMR (400 MHz, $CDCl_3$) δ 3.55-3.47 (m, 1H) , 2.69-0.65 (m, 1H), 2.52-2.38 (m, 3H), 2.22-2.12 (m, 2H), 2.04 (s, 3H), 1.80-0.97 (m, 15H), 0.95 (s, 3H), 0.83-0.75 (m, 1H), 0.51 (s, 3H).

**3α-Hydroxy-17β-(2-methyl-1,3-dioxolan-2-yl)-5β-androstan-11-one (4).** To a stirred solution of steroid **3** (2.9 g, 8.7 mmol) in benzene (150 mL) was added ethylene glycol (10 mL) and PTSA (100 mg) at 23 °C. The reaction flask was equipped with a Dean-Stark apparatus and refluxed for 16 h, then allowed to cool to 23 °C. Aqueous $NaHCO_3$ was added and the product was extracted into EtOAc (300 mL). The extract was washed with brine (100 mL × 5), dried over anhydrous $Na_2SO_4$, filtered, the solvent removed and the residue was dried under high vacuum to give steroid **4** (3.28 g, 100%): $^1$H NMR (400 MHz, $CDCl_3$) δ 3.99-3.84 (m, 4H), 3.59-3.54 (m, 1H), 2.59-2.57 (m, 1H), 2.51-2.46 (m, 1H), 2.26-2.23 (m, 1H), 2.04-1.99 (m, 1H), 1.87-1.03 (m, 17H), 1.25 (s, 3H), 1.01 (s, 3H), 0.87-0.79 (m, 1H), 0.70 (s, 3H); $^{13}$C NMR (100 MHz, $CDCl_3$) δ 211.5, 111.2, 70.9, 64.8, 64.2, 63.1, 57.7, 56.8, 55.4, 46.0, 44.9, 37.6, 36.3, 35.6, 34.9, 32.5, 21.2, 28.0, 24.3, 23.4, 23.3, 14.1, 12.0.

**3α-(Ethoxymethoxy)-17β-(2-methyl-1,3-dioxolan-2-yl)-5β-androstan-11-one (5).** To a stirred solution of steroid **4** (3.28 g, 8.7 mmol) in $CH_2Cl_2$ (100 mL) was added chloromethyl ethyl ether (1.6 g, 17.4 mmol) and (*i*-Pr)$_2$NEt (4.5 g, 35 mmol) at 0 °C and the reaction was warmed to 23 °C. After 16 h, the solvent was removed and the residue was purified by flash column chromatography (silica gel, eluted with 20%-40% EtOAc in hexanes) to give steroid **5** (3.6 g , 95%): $^{1}$H NMR (400 MHz, $CDCl_3$) δ 4.64-4.59 (m, 2H), 3.89-3.73 (m, 4H), 3.53-3.43 (m, 2H), 3.42-3.35 (m, 1H), 2.50-2.47 (m, 1H), 2.42-2.37 (m, 1H), 2.18-2.10 (m, 1H), 1.94-1.90 (m, 1H), 1.79-0.94 (m, 19H), 1.15 (s, 3H), 0.92 (s, 3H), 0.76-0.64 (m, 1H), 0.61 (s, 3H); $^{13}$C NMR (100 MHz, $CDCl_3$) δ 211.0, 110.9, 92.7, 75.5, 64.6, 64.0, 62.9, 62.7, 57.5, 56.6, 55.2, 45.7, 44.7, 36.1, 35.4, 34.9, 34.6, 32.3, 28.2, 27.9, 24.1, 23.2, 23.1, 14.9, 13.9, 11.8.

**3α-(Ethoxymethoxy)-17β-(2-methyl-1,3-dioxolan-2-yl)-5β-androstan-11β-ol (6).** Steroid **5** (2.2 g, 5.2 mmol) was dissolved in dry $Et_2O$ and added dropwise to a stirred suspension of LAH (600 mg, 15.7 mmol) in $Et_2O$ at 0 °C. The mixture was warmed to 23 °C and stirring continued for 2 h. Water (0.6 mL) was added, then aqueous 15% NaOH (0.6 mL) was added and the mixture was stirred for 0.5 h. Additional water (1.8mL) was added and stirring continued for 15 min. Anhydrous $MgSO_4$ was added and after 15 min the mixture was filtered through celite. Solvent was removed from the filtrate and the residue was purified by flash column chromatography (silica gel, eluted with 20-30% EtOAc in hexanes) to give steroid **6** (1.45 g, 66%): $^{1}$H NMR (400 MHz, $CDCl_3$) δ 4.57 (s, 2H), 4.10 (s, 1H), 3.85-3.70 (m, 4H), 3.48-3.43 (m, 2H), 3.34-3.32 (m, 1H), 2.05-2.01 (m, 1H), 1.76-0.76 (m, 23H), 1.14 (s, 3H), 0.90 (s, 3H), 0.84 (s, 3H), 0.54-0.51 (m, 1H); $^{13}$C NMR (100 MHz, $CDCl_3$) δ 111.5, 92.6, 75.6, 67.9, 64.7, 62.9, 62.6, 58.3, 57.9, 57.6, 48.4, 45.4, 41.0, 36.4, 35.4, 34.5, 32.0, 30.5, 28.0, 28.0, 24.2, 23.4, 22.4, 15.4, 14.9, 14.86.

**3α-(Ethoxymethoxy)-11β-(2-propenyloxy)-17β-(2-methyl-1,3-dioxolan-2-yl)-5β-androstane (7).** To a stirred solution of steroid **6** (1.39 g, 3.18 mmol) in THF (50mL) was added KH (1 g, 25 mmol) at 23 °C. The mixture was refluxed for 50 min. Allyl bromide (2 mL) was added and stirring continued for 2 h. After cooling to 23 °, MeOH (5 mL) was added and stirring continued for 10 min. Water (50 mL) was added and the product was extracted into EtOAc (50 mL x 3). The combined extracts were dried over anhydrous $Na_2SO_4$, filtered, the solvent removed and the residue purified by flash column chromatography (silica gel, eluted with 5% EtOAc in hexanes) to give steroid **7** (1.32 g, 87%): $^{1}$H NMR (400 MHz, $CDCl_3$) δ 5.94-5.86 (m, 1H), 5.26-5.21 (m,

1H), 5.08-5.06 (m, 1H), 4.72-4.70 (m, 2H), 4.14-3.45 (m, 11H), 2.50-2.43 (m, 1H), 2.11-0.65 (m, 22H), 1.20 (s, 3H), 1.00 (s, 3H), 0.92 (s, 3H); $^{13}C$ NMR (100 MHz, $CDCl_3$) δ 135.6, 115.3, 112.0, 93.0, 76.1, 75.4, 69.1, 65.1, 63.7, 63.3, 63.0, 58.6, 58.4, 58.2, 45.8, 41.6, 40.6, 37.1, 35.8, 34.8, 32.5, 31.4, 28.4, 28.3, 24.5, 23.7, 22.7, 15.3, 14.5.

**3-(((3α-Ethoxymethoxy)-17β-(2-methyl-1,3-dioxolan-2-yl)-5β-androstan-11β-yl)oxy)-propan-1-ol (8).** To a stirred solution of steroid **7** (1.24 g, 2.6 mmol) in THF (10mL) was added 9-BBN (10 mL, 5 mmol, 0.5 M in THF) at 0 °C. After 4 h, the reaction was allowed to warm to 23 °C and stirring continued for 16 h. 3 N NaOH (5 mL, 15 mmol) and $H_2O_2$ (5 mL) were added, and the reaction was stirred for another 1 h. The product was extracted into EtOAc (100 mL x 2). The combined extracts were washed with brine (50 mL x 3), dried over anhydrous $Na_2SO_4$, filtered, the solvent removed and the residue purified by flash column chromatography (silica gel, eluted with 10-20% EtOAc in hexanes) to give steroid **8** (810 mg, 65%): $^1H$ NMR (400 MHz, $CDCl_3$) δ 4.73-4.70 (m, 2H), 4.13-3.85 (m, 4H), 3.76-3.67 (m, 4H), 3.62-3.58 (m, 2H), 3.56-3.45 (m, 1H), 3.30-3.25 (m, 1H), 2.52,-2.47 (m, 1H), 2.11- 0.65 (m, 26H), 1.29 (s, 3H), 0.99 (s, 3H), 0.91 (s, 3H); $^{13}C$ NMR (100 MHz, $CDCl_3$) δ 111.9, 93.0, 76.0, 67.1, 65.1, 63.7, 63.3, 63.0, 61.9, 58.6, 58.3, 58.1, 45.8, 41.5, 40.6, 37.1, 35.7, 34.8, 32.8, 32.5, 31.3, 28.4, 28.2, 24.5, 23.6, 22.7, 15.6, 15.1, 14.6.

**3-(((3α-Ethoxymethoxy)-17β-(2-methyl-1,3-dioxolan-2-yl)-5β-androstan-11β-yl)oxy)-propan-1-al (9).** To a stirred solution of oxalyl chloride (0.5 mL) in $CH_2Cl_2$ (30 mL) was added DMSO (0.6 mL) in $CH_2Cl_2$ (10mL) at -78 °C. After 10 min, steroid **8** (810 mg, 1.6 mmol) in $CH_2Cl_2$ (10 mL + 3 mL + 3 mL) was added. After 2 h, $Et_3N$ (2 mL) was added and the reaction was allowed to warm to 23 °C and stirring continued for 30 min. The product was extracted into $CH_2Cl_2$ (150 mL). The extract was dried over anhydrous $Na_2SO_4$, filtered, the solvent removed and he residue purified by flash column chromatography (silica gel, eluted with 20-50% EtOAc in hexanes) to give steroid **9** (500 mg, 62%): $^1H$ NMR (400 MHz, $CDCl_3$) δ 9.71-9.70 (m, 1H), 4.65-4.61 (m, 2H), 3.94-3.79 (m, 5H), 3.68-3.67 (m, 1H), 3.56-3.50 (m, 2H), 3.43-3.37 (m, 2H), 2.56-2.42 (m, 3H), 1.78-0.58 (m, 23H), 1.23 (s, 3H), 0.86 (s, 3H), 0.81 (s, 3H); $^{13}C$ NMR (100 MHz, $CDCl_3$) δ 201.8, 111.8, 93.0, 76.4, 75.9, 65.0, 63.2, 62.9, 62.0, 58.4, 58.1, 57.9, 45.8, 44.2, 41.5, 40.5, 37.0, 35.7, 34.8, 32.4, 31.3, 28.3, 28.2, 24.5, 23.6, 22.6, 15.3, 15.1, 14.4.

**11β-(But-3-yn-1-yloxy)-3α-(ethoxymethoxy)-17β-(2-methyl-1,3-dioxolan-2-yl)-5β-androstane (10).** To a stirred solution steroid **9** (500 mg, 1.0 mmol) in MeOH/THF (10 mL/10 mL) was added dimethyl (1-diazo-2-oxopropyl) phosphonate (340 mg, 1.76 mmol) and $K_2CO_3$ (1 g, fine powder) at 23 °C. After 16 h, water was added and the product was extracted into EtOAc (100 mL × 3). The combined extracts were dried over anhydrous $Na_2SO_4$, filtered, the solvent removed and the residue purified by flash column chromatography (silica gel, eluted with 20% EtOAc in hexanes) to give steroid **10** (340 mg, 68%): $^1$H NMR (400 MHz, $CDCl_3$) δ 4.68-4.64 (m, 2H), 3.94-3.79 (m, 4H), 3.68-3.60 (m, 2H), 3.57-3.51 (m, 2H), 3.45-3.39 (m, 1H), 3.22- 3.16 (m, 1H), 2.44-2.33 (m, 3H), 1.90-0.58 (m, 24H), 1.24 (s, 3H), 0.95 (s, 3H), 0.85 (s, 3H); $^{13}$C NMR (100 MHz, $CDCl_3$) δ 111.9, 93.0, 82.1, 76.1, 76.0, 69.1, 66.5, 65.1, 63.2, 62.9, 58.5, 58.2, 58.0, 45.8, 41.5, 40.6, 36.8, 35.8, 34.8, 32.5, 31.3, 28.3, 28.2, 24.5, 23.6, 22.6, 20.1, 15.2, 15.1, 14.2.

**3α-(Ethoxymethoxy)-17β-(2-methyl-1,3-dioxolan-2-yl)-11β-((1-phenyl-2,5,8,11-tetraoxaheptadec-14-yn-17-yl)oxy)-5β-androstane (11).** To a mixture of steroid **10** (340 mg, 0.7 mmol), 13-iodo-1-phenyl-2,5,8,11-tetraoxatridecane (40 0mg, 1.02 mmol), Cu(I)I (7.7 mg, 0.04 mmol), $CsCO_3$ (332 mg, 1.02 mmol) and bis[(2-dimethylamino)phenyl]amine nickel(II) chloride (48 mg, 0.14 mmol) was added dioxane (12 mL). The stirred mixture was heated to 140 °C for 16 h and then cooled to 23 °C. The solvent was removed and the residue was purified by flash column chromatography (silica gel, eluted with 20-50% EtOAc in hexanes) to give steroid **11** (460 mg, 87%): $^1$H NMR (400 MHz, $CDCl_3$) δ 7.34-7.27 (m, 5H), 4.74-4.70 (m, 2H), 4.56 (s, 2H), 3.99-3.84 (m, 4H), 3.75-3.44 (m, 20H), 3.21-3.15 (m, 1H), 2.47-2.35 (m, 5H), 1.82-0.62 (m, 22H), 1.29 (s, 3H), 0.99 (s, 3H), 0.89 (s, 3H); $^{13}$C NMR (100 MHz, $CDCl_3$) δ 138.2, 128.4 (2 x C), 127.8 (2 x C), 127.6, 112.0, 93.0, 78.8, 77.5, 76.0, 75.9, 73.2, 70.6 (4 x C), 70.5, 70.2, 69.9, 69.4, 67.0, 65.1, 63.3, 63.0, 58.6, 58.2, 58.1, 45.8, 41.6, 40.6, 36.9, 35.8, 34.8, 32.5, 31.4, 28.5, 24.5, 23.6, 22.7, 20.4, 20.0, 15.2, 15.1, 14.3.

**3α-(Ethoxymethoxy)-17β-(2-methyl-1,3-dioxolan-2-yl)-11β-((1-phenyl-2,5,8,11-tetraoxaheptadecan-17-yl)oxy)-5β-androstane (12).** To a stirred solution of steroid **11** (460 mg, 0.6 mmol) in EtOAc (30 mL) was added Pd/C (10%, 70 mg) in a Parr hydrogenation flask and hydrogenation was carried out overnight at 55 psi $H_2$. The mixture was filtered through celite and washed with EtOAc. The solvent was removed to give steroid **12** (460 mg, 100%): $^1$H NMR (400 MHz, $CDCl_3$) δ 7.33-7.26 (m, 5H), 4.58-4.59 (m, 2H), 4.72 (s, 1H), 4.56 (s, 1H), 3.99-3.41 (m,

23H), 3.08-3.04 (m, 1H), 2.47-2.44 (m, 1H), 1.92-0.06 (m, 31H), 1.29 (s, 3H), 0.98 (s, 3H), 0.89 (s, 3H); $^{13}$C NMR (100 MHz, $CDCl_3$) δ 138.2, 128.3 (2 x C), 127.7 (2 x C), 127.6, 120.0, 93.0, 76.1, 75.8, 73.2, 71.5, 70.6 (5 x C), 70.0, 69.4, 68.2, 65.2, 63.3, 63.0, 58.6, 58.3, 58.1, 45.8, 41.6, 40.6, 36.9, 35.8, 34.9, 32.5, 31.4, 30.3, 29.6, 28.4, 28.3, 26.5, 24.6, 23.7, 22.6, 15.2 (2 x C), 14.4.

**2-(2-(2-((6-((-3α-(Ethoxymethoxy)-17β-(2-methyl-1,3-dioxolan-2-yl)-5β-androstan-11β-yl)oxy)hexyl)oxy)ethoxy)ethoxy)-ethan-1-ol (13).** To a solution of anhydrous liquid ammonium (30 mL) was added sodium metal (100 mg) at -78 °C. The mixture was stirred 15 min. Steroid **12** (460 mg, 0.61 mmol) in THF (10 mL + 2 mL + 2 mL) was added dropwise. After 1 h, solid $NH_4Cl$ (2.0 g) was added and the mixture was allowed to warm to 23 °C allowing the ammonia to evaporate. Water was added and the product was extracted into EtOAc (100mL x 3). The combined extracts were dried over anhydrous $Na_2SO_4$, filtered, the solvent removed and the residue purified by flash column chromatography (silica gel, eluted with 3-5% MeOH in $CH_2Cl_2$) to give steroid **13** (310 mg, 76%): $^{1}$H NMR (400 MHz, $CDCl_3$) δ 4.70 (s, 2H), 3.99-3.82 (m, 4H), 3.71-3.40 (m, 19H), 3.05-3.03 (m, 1H), 2.46-2.42 (m, 1H), 1.80-0.59 (m, 32H), 1.27 (s, 3H), 0.96 (s, 3H), 0.87 (s, 3H); $^{13}$C NMR (100 MHz, $CDCl_3$) δ 112.0, 93.0, 76.1, 75.8, 72.5, 71.5, 70.6, 70.5, 70.3, 70.0, 68.2, 65.2, 63.2, 62.9, 61.9, 58.6, 58.3, 58.1, 45.8, 41.6, 40.6, 36.9, 35.8, 34.8, 32.5, 31.4, 30.3, 29.5, 28.4, 28.3, 26.4, 26.0, 24.5, 23.6, 22.6, 15.2 (2 x C), 14.3.

**2-(2-(2-((6-((-3α-(Ethoxymethoxy)-17β-(2-methyl-1,3-dioxolan-2-yl)-5β-androstan-11β-yl)oxy)hexyl)oxy)ethoxy)ethoxy)-acetaldehyde (14).** To a solution of oxalyl chloride (0.5 mL) in stirred $CH_2Cl_2$ (10 mL) was added DMSO (0.6 mL) in $CH_2Cl_2$ (10 mL) at -78°C. After 10 min, steroid **13** (310 mg, 0.46 mmol) in $CH_2Cl_2$ (3mL + 2mL + 1mL) was added. After 2 hours, $Et_3N$ (4 mL) was added and the reaction was allowed to warm to 23 °C and stirring continued for 30 min. The product was extracted into $CH_2Cl_2$ (50 mL). The extract was dried over anhydrous $Na_2SO_4$, filtered, the solvent removed and the residue was purified by flash column chromatography (silica gel, eluted with 20-50% EtOAc in hexanes) to give steroid **14** (200 mg, 65%): $^{1}$H NMR (400 MHz, $CDCl_3$) δ 9.67 (s, 1H), 4.69-4.65 (m, 2H), 4.11-3.30 (m, 21H), 3.04-2.99 (m, 1H), 2.43-2.39 (m, 1H), 1.78-0.57 (m, 31H), 1.15 (s, 3H), 0.94 (s, 3H), 0.85 (s, 3H); $^{13}$C NMR (100 MHz, $CDCl_3$) δ 200.9, 111.9, 93.0, 76.0, 75.7, 71.4, 71.2, 70.7, 70.6, 70.0, 68.1, 65.2, 63.2, 62.9, 58.6, 58.3, 58.1, 45.8, 41.5, 40.6, 36.9, 35.8, 34.8, 32.5, 31.4, 30.3, 29.5, 28.4, 28.2, 26.4, 25.9, 24.5, 23.6, 22.6, 15.1 (3 x C), 14.3.

**2-(3α-(Ethoxymethoxy)-11β-((6-(2-(2-(prop-2-yn-1-yloxy)ethoxy)ethoxy)hexyl)oxy)-5β-androstan-17β-yl)-2-methyl-1,3-dioxolane (15).** To a stirred solution of steroid **14** (190 mg, 0.29 mmol) in MeOH/THF (5 mL/5 mL) was added dimethyl (1-diazo-2-oxopropyl) phosphonate (340 mg, 1.76 mmol) and $K_2CO_3$ (1 g, fine powder) at 23 °C. After 16 h, water was added and the product was extracted into EtOAc (50 mL × 3). The combined extracts were dried over anhydrous $Na_2SO_4$, filtered, the solvent removed and the residue purified by flash column chromatography (silica gel, eluted with 20-50% EtOAc in hexanes) to give steroid **15** (150 mg, 79%): $^1H$ NMR (400 MHz, $CDCl_3$) δ 4.69 (s, 2H), 4.17-4.16 (m, 2H), 3.97-3.82 (m, 4H), 3.68-3.39 (s, 13H), 3.05-3.00 (m, 1H), 2.44-2.40 (m, 2H), 1.80-0.58 (m, 33H), 1.27 (s, 3H), 0.95 (s, 3H), 0.86 (s, 3H); $^{13}C$ NMR (100 MHz, $CDCl_3$) δ 112.0, 93.0, 79.6, 76.1, 75.8, 74.5, 71.4, 70.6, 70.4, 70.0, 69.1, 68.2, 65.2, 63.2, 62.9, 58.6, 58.4, 58.3, 58.1, 45.8, 41.6, 40.6, 36.9, 35.8, 34.8, 32.5, 31.4, 30.3, 29.5, 28.4, 28.3, 26.4, 26.0, 24.5, 23.6, 22.6, 15.1 (2 x C), 14.3.

**3α-(Hydroxy)-11β-((6-(2-(2-(prop-2-yn-1-yloxy)ethoxy)ethoxy)hexyl)oxy)-5β-pregnan-20-one (16, YX89).** To a stirred solution of steroid **15** steroid (150 mg, 0.23 mmol) in MeOH (20mL) was added 6 N HCl (5 mL). After 16 h, the product was extracted into EtOAc (100 mL x 3). The combined extracts were dried over anhydrous $Na_2SO_4$, filtered, the solvent removed and the residue purified by flash column chromatography (silica gel, eluted with 50-100% EtOAc in hexanes) to give **YX89** (**16**, 91 mg, 71%): $^1H$ NMR (400 MHz, $CDCl_3$) δ 4.18-4.17 (m, 2H), 3.71-3.39 (m, 13H), 3.09-3.03 (m, 1H), 2.42-2.38 (m, 3H), 2.14-0.63 (m, 28H), 2.09 (s, 3H), 0.96 (s, 3H), 0.73 (s, 3H); $^{13}C$ NMR (100 MHz, $CDCl_3$) δ 209.7, 79.6, 75.5, 74.6, 71.4, 71.0, 70.6, 70.4, 70.0, 69.1, 68.3, 64.1, 58.4, 58.3, 58.2, 45.8, 43.7, 40.0, 37.7, 37.0, 35.6, 32.5, 31.8, 31.5, 31.2, 30.2, 29.5, 28.1, 26.3, 25.9, 24.3, 22.5, 15.2, 14.7.

**3α-(Hydroxy)-11β-(2-propenyloxy)-5β-pregnan-20-one (17, YX88).** To a stirred solution of steroid **7** (80 mg, 0.17 mmol) in MeOH (10 mL) was added 6 N HCl (5 mL). After 16 h, the product was extracted into EtOAc (50 mL x 3). The combined extracts were dried over anhydrous $Na_2SO_4$, filtered, the solvent removed and the residue purified by flash column chromatography (silica gel, eluted with 50-100% EtOAc in hexanes) to give **YX88** (**17**, 63 mg, 99%): $^1H$ NMR (400 MHz, $CDCl_3$) δ 5.91-5.84 (m, 1H), 5.24-5.20 (m, 1H), 5.09-5.06 (m, 1H), 4.08-4.03 (m, 3H), 3.82-3.81 (m, 1H), 3.74-3.69 (m, 1H), 3.58-3.52 (m, 1H), 2.46-2.40 (m, 2H), 2.14-2.11 (m, 4H), 1.81-0.69

(m, 14H), 2.11 (s, 3H), 0.99 (s, 3H), 0.77 (s, 3H); $^{13}C$ NMR (100 MHz, $CDCl_3$) δ 209.7, 135.2, 115.7, 75.2, 71.1, 69.2, 64.1, 58.4, 58.3, 45.8, 43.6, 40.1, 37.7, 37.1, 35.7, 32.5, 31.7, 31.4, 31.2, 28.1, 24.3, 22.6, 15.4, 14.8.

**DART Compound synthesis Methods:**

### 1. General Methods

All reactions were conducted in oven-dried glassware under nitrogen. Unless otherwise stated, all reagents were purchased from commercial suppliers (Sigma–Aldrich, Acros, Fisher, Alfa Aesar, TCI, or Ambeed) and used without further purification. All solvents were American Chemical Society (ACS) grade or better and used without further purification. Analytical thin layer chromatography (TLC) was performed with glass-backed silica gel (60 Å) plates with fluorescent indication (Whatman). Visualization was accomplished by UV irradiation at 254 nm and/or by staining with p-anisaldehyde solution or potassium permanganate solution followed by heating. Flash column chromatography was performed by using silica gel (particle size 230−400 mesh, 60 Å) purchased from Silicycle. All $^{1}H$ NMR spectra were recorded with a Bruker spectrometer (500 MHz or 700 MHz). All NMR δ values are given in parts per million (ppm) and are referenced to the residual isotopomer solvent signals ($CD_3OD$: $\delta = 3.31$ ppm) for $^{1}H$ NMR spectra. Coupling constants (J) are given in Hertz (Hz) and multiplicities are indicated using the conventional abbreviation (s = singlet, d = doublet, t = triplet, m = multiplet or overlap of non-equivalent resonances, br = broad). Electrospray ionization (ESI) mass spectrometry (MS) was recorded with an Agilent 6224 series (LC/MS–TOF) spectrometer to obtain the molecular masses of the compounds.

### 2. Synthesis of Neurosteroid-DART Probes

#### (2.1) Preparation of azide-PEG$_{36}$-HTL.2

The **azide-PEG$_{36}$-HTL.2** was prepared following the synthetic procedure reported for the azide$^{DART.2}$ (HTL.2) module in Shields et al. (Shields et al., 2024)with only minor modifications to accommodate our reaction scale and purification conditions. The overall route, including construction of the PEG$_{36}$ linker and its conjugation to the HTL warhead, was identical to that described previously. The structure and analytical data ($^{1}H$ NMR and HRMS) of **azide-PEG$_{36}$-HTL.2** were consistent with those reported for azide$^{DART.2}$ (HTL.2).

#### (2.2) Preparation of MQ302$^{DART.2}$

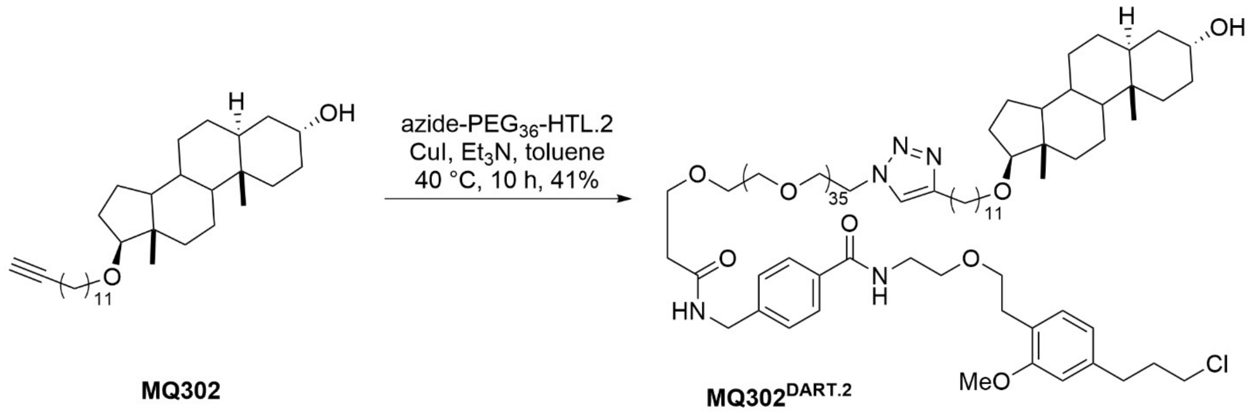


To a solution of **MQ302** (6 mg, 0.013 mmol) in toluene (2.0 mL) were added **azide-PEG$_{36}$-HTL.2** (27 mg, 0.013 mmol), CuI (7.3 mg, 0.039 mmol) and $Et_3N$ (318 mg, 438 μL, 3.12 mmol) at 25 °C. The resulting reaction mixture was stirred at 40 °C for 10 h and concentrated *in vacuo*. The obtained residue was then dissolved in $CH_2Cl_2$ and washed with ice-cold $NH_4OH$ and $H_2O$. The organic layer was dried over $Na_2SO_4$ and the solvent was evaporated *in vacuo*. The resulting residue was purified by column chromatography (silica gel, $CHCl_3$/MeOH/$NH_4OH$, 15:1:0.05) to afford **MQ302$^{DART.2}$** (13.6 mg, 0.0053 mmol, 41%): $^1$H NMR (500 MHz, $CD_3OD$) δ 7.83–7.73 (m, 3H), 7.40 (d, *J* = 8.1 Hz, 2H), 7.06 (d, *J* = 7.5 Hz, 1H), 6.76 (d, *J* = 1.6 Hz, 1H), 6.65 (dd, *J* = 7.6, 1.6 Hz, 1H), 4.58 (t, *J* = 5.5 Hz, 1H), 4.54 (t, *J* = 5.1 Hz, 1H), 4.47 (d, *J* = 4.8 Hz, 2H), 3.95 (dd, *J* = 6.5, 4.3 Hz, 2H), 3.88 (t, *J* = 5.1 Hz, 1H), 3.82–3.72 (m, 6H), 3.65–3.56 (m, 170H), 3.53 (t, *J* = 6.3 Hz, 5H), 3.50–3.40 (m, 4H), 2.84 (t, *J* = 6.9 Hz, 2H), 2.74–2.61 (m, 4H), 2.52 (t, *J* = 6.0 Hz, 2H), 2.07–1.93 (m, 3H), 1.88 (dt, *J* = 12.4, 3.4 Hz, 1H), 1.70–1.28 (m, 36H), 0.82 (s, 3H), 0.75 (s, 3H); HRMS (ESI-TOF) m/z: $[M + 3H]^{3+}$ Calcd for $C_{129}H_{230}ClN_5O_{42}$ 853.1974; Found 853.1964.

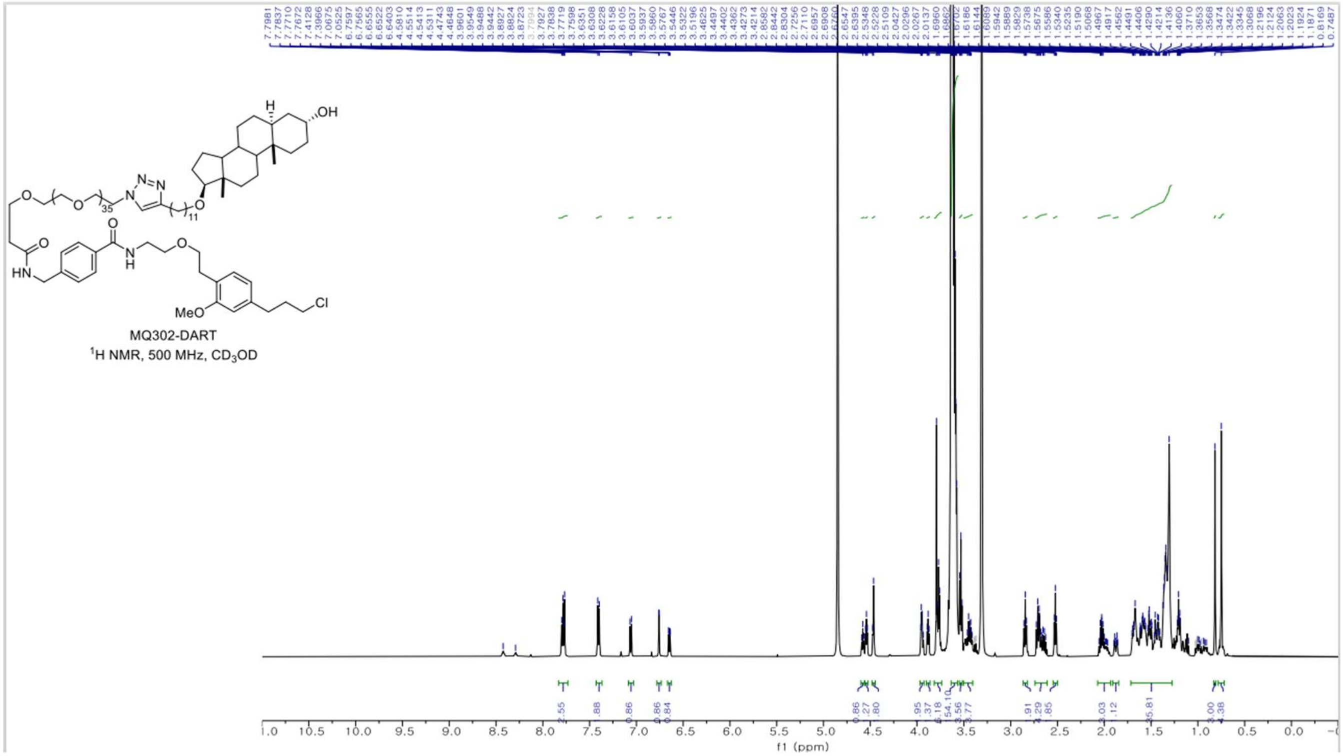


MQ302-DART
$^{1}$H NMR, 500 MHz, $CD_3OD$

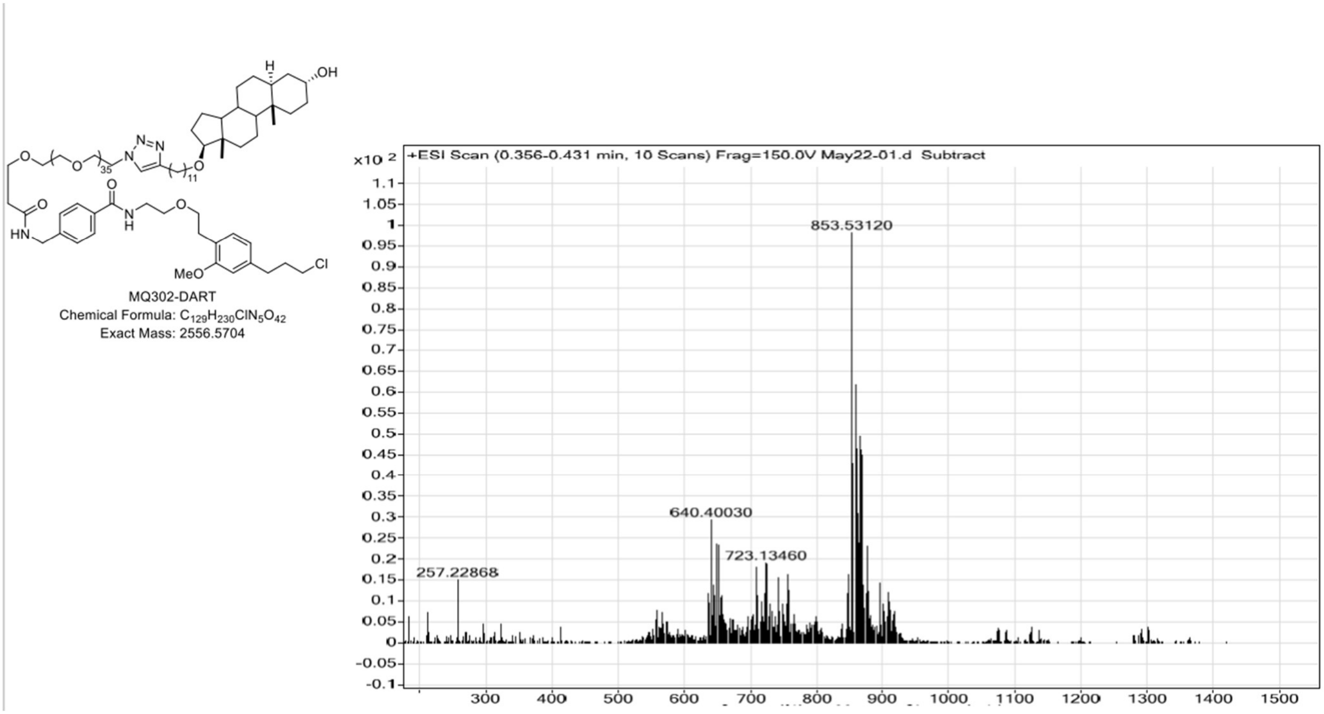


**(2.3) Preparation of MQ303$^{DART.2}$**

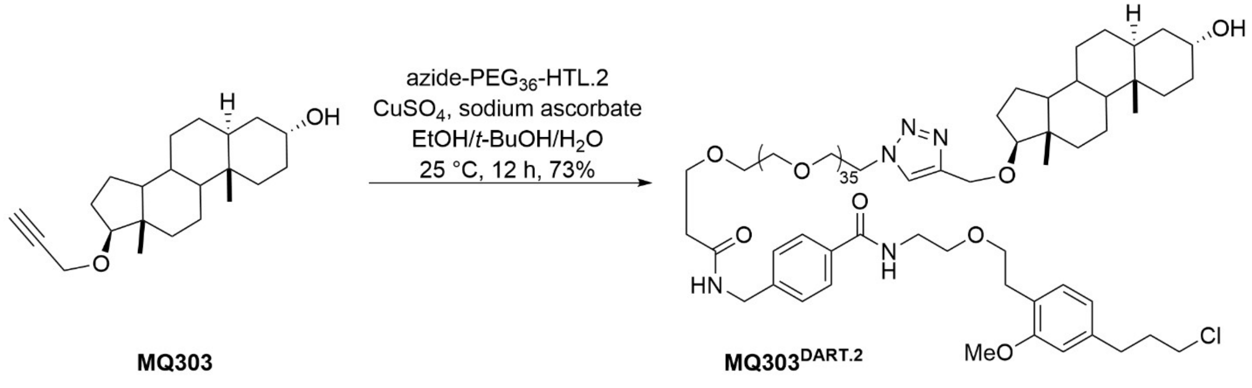


To a solution of **MQ303**(5.3 mg, 0.016 mmol) in EtOH/t-BuOH/$H_2O$ (2:1:1, 1.5 mL) were added $CuSO_4$ (3.2 mg, 0.02 mmol) and sodium ascorbate (4 mg, 0.02 mmol) at 25 °C. After stirring for 5 min, **azide-PEG$_{36}$-HTL.2** (33.5 mg, 0.016 mmol) was added. The resulting reaction mixture was stirred at 25 °C for 12 h and concentrated *in vacuo*. The residue was diluted with MeCN, filtered through a cotton plug, and washed with MeCN. The filtrate was concentrated in vacuo, and the resulting residue was purified by a reversed-phase semi-preparative HPLC column (gradient elution: 20–80% MeCN in $H_2O$ containing 0.1% TFA; detection wavelength: 254 nm; other parameters as described in the General Methods) to afford **MQ303$^{DART.2}$** (28.2 mg, 0.012 mmol, 73%): $^1$H NMR (500 MHz, $CD_3OD$) δ 7.99 (s, 1H), 7.77 (d, *J* = 8.0 Hz, 2H), 7.40 (d, *J* = 8.1 Hz, 2H), 7.06 (d, *J* = 7.5 Hz, 1H), 6.76 (s, 1H), 6.65 (d, *J* = 7.5 Hz, 1H), 4.61–4.56 (m, 4H), 4.47 (s, 2H), 3.97–3.94 (m, 1H), 3.89 (t, *J* = 5.1 Hz, 2H), 3.81–3.75 (m, 6H), 3.67–3.57 (m, 158H), 3.53 (t, *J* = 6.2 Hz, 4H), 3.50–3.44 (m, 2H), 2.84 (t, *J* = 7.0 Hz, 2H), 2.71 (t, *J* = 7.4 Hz, 2H), 2.52 (t, *J* = 6.0 Hz, 2H), 2.06–1.96 (m, 3H), 1.87 (d, *J* = 12.6 Hz, 1H), 1.72–1.18 (m, 18H), 1.18–1.10 (m, 1H), 1.05–0.88 (m, 2H), 0.82 (s, 3H), 0.80–0.71 (m, 4H); HRMS (ESI-TOF) m/z: $[M + 3H]^{3+}$ Calcd for $C_{119}H_{213}ClN_5O_{42}$ 806.4786; Found 806.4791.

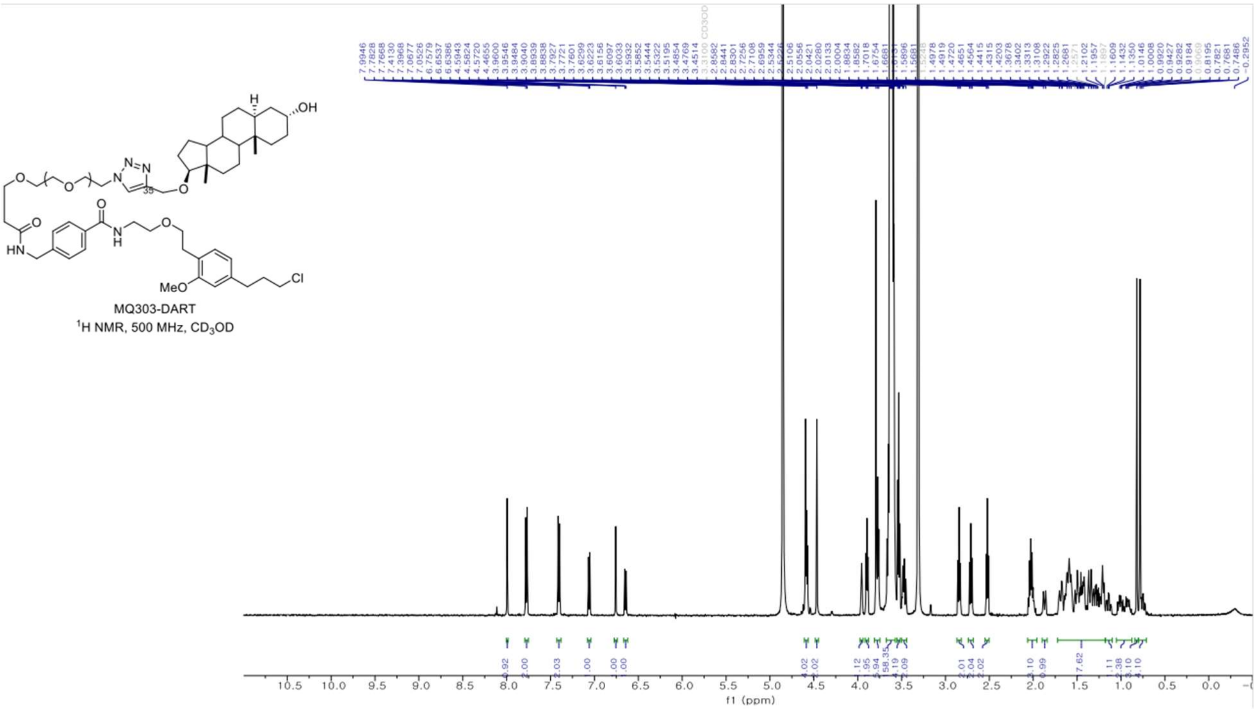

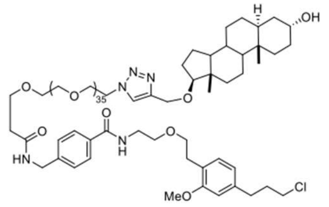

MQ303-DART
Chemical Formula: $C_{119}H_{210}ClN_5O_{42}$
Exact Mass: 2416.4139

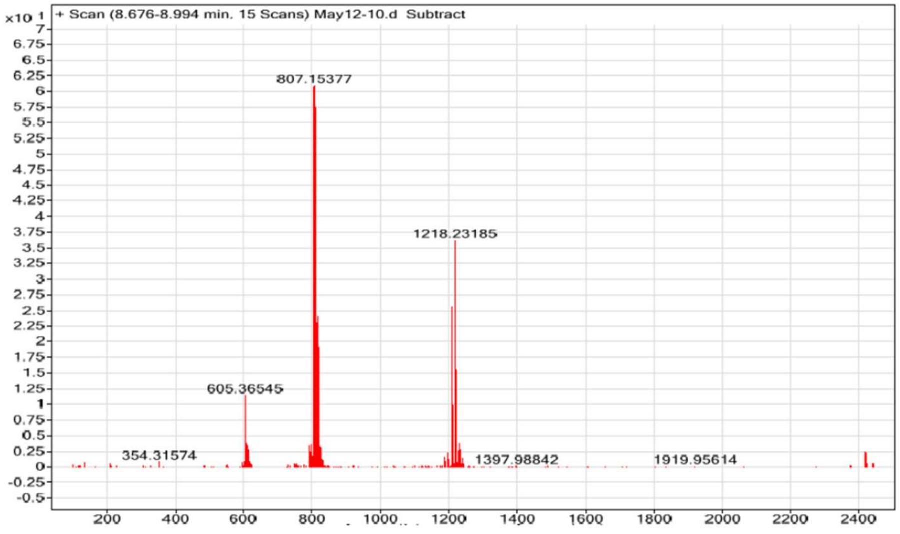


#### (2.4) Preparation of MQ311$^{DART.2}$

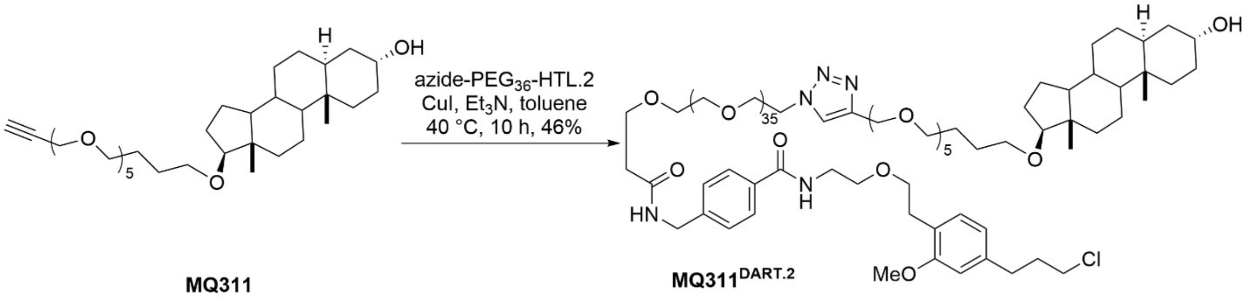


To a solution of **MQ311** (12 mg, 0.021 mmol) in toluene (2.0 mL) were added **azide-PEG$_{36}$-HTL.2** (20 mg, 0.01 mmol), CuI (12 mg, 0.063 mmol) and $Et_3N$ (214 mg, 294 µL, 2.1 mmol) at 25 °C. The resulting reaction mixture was stirred at 40 °C for 10 h and concentrated *in vacuo*. The obtained residue was then dissolved in $CH_2Cl_2$ and washed with ice-cold $NH_4OH$ and $H_2O$. The organic layer was dried over $Na_2SO_4$ and the solvent was evaporated *in vacuo*. The resulting residue was purified by column chromatography (silica gel, $CHCl_3$/MeOH/$NH_4OH$, 20:1:0.05) to afford **MQ311$^{DART.2}$** (25.4 mg, 0.0095 mmol, 46%); $^1$H NMR (500 MHz, $CD_3OD$) δ 8.05 (s, 1H), 7.80–7.74 (m, 2H), 7.41 (d, $J$ = 8.3 Hz, 2H), 7.06 (d, $J$ = 7.6 Hz, 1H), 6.76 (d, $J$ = 1.6 Hz, 1H), 6.65 (dd, $J$ = 7.5, 1.6 Hz, 1H), 4.66–4.56 (m, 4H), 4.47 (s, 2H), 3.98–3.94 (m, 2H), 3.90 (t, $J$ = 5.0 Hz, 2H), 3.79 (s, 3H), 3.77 (t, $J$ = 6.0 Hz, 3H), 3.63 (d, $J$ = 3.7 Hz, 133H), 3.53 (t, $J$ = 6.4 Hz,

4H), 3.51–3.43 (m, 6H), 2.84 (t, $J$ = 7.1 Hz, 2H), 2.71 (t, $J$ = 7.4 Hz, 2H), 2.52 (t, $J$ = 6.0 Hz, 2H), 2.06–1.95 (m, 3H), 1.91–1.84 (m, 1H), 1.72–1.56 (m, 12H), 1.53–1.41 (m, 5H), 1.39–1.17 (m, 10H), 1.04–0.89 (m, 3H), 0.82 (s, 3H), 0.75 (s, 3H); HRMS (ESI-TOF) m/z: $[M + 3H]^{3+}$ Calcd for $C_{131}H_{234}ClN_5O_{47}$ 888.1921; Found 889.1985.

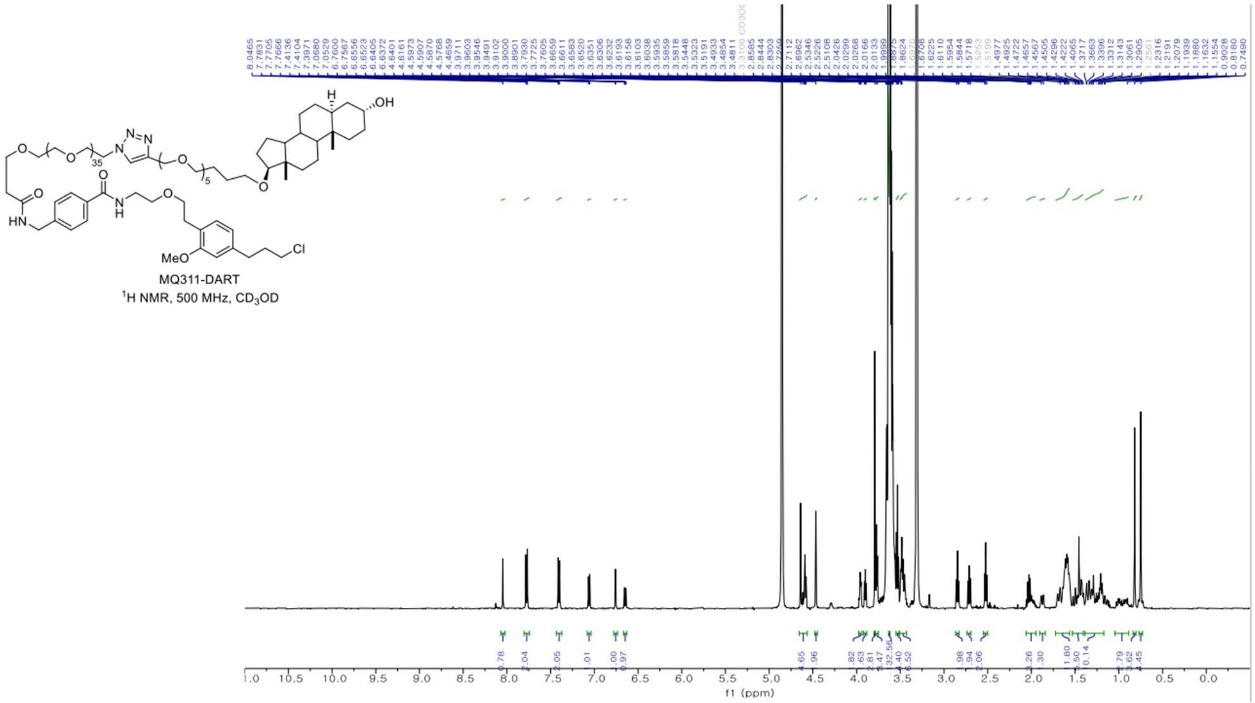

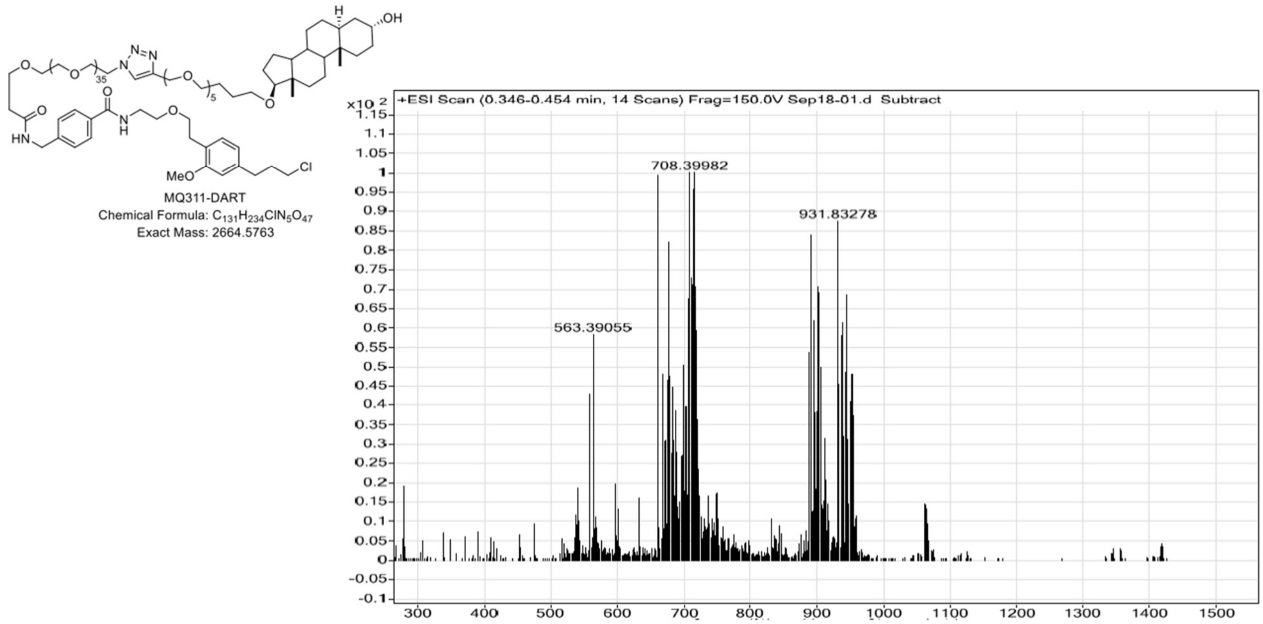


### (2.5) Preparation of MQ369$^{DART.2}$

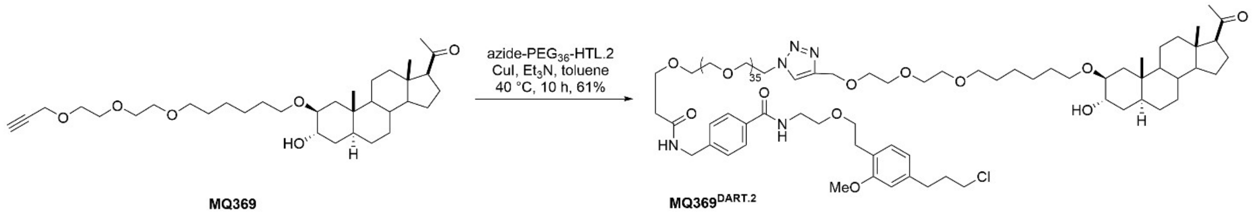


**MQ369** **MQ369$^{DART.2}$**

To a solution of **MQ369** (45 mg, 0.089 mmol) in toluene (4.4 mL) were added **azide-PEG$_{36}$-HTL.2** (185 mg, 0.089 mmol), CuI (51 mg, 0.27 mmol) and $Et_3N$ (453 mg, 624 μL, 4.45 mmol) at 25 °C. The resulting reaction mixture was stirred at 40 °C for 10 h and concentrated *in vacuo*. The obtained residue was then dissolved in $CH_2Cl_2$ and washed with ice-cold $NH_4OH$ and $H_2O$. The organic layer was dried over $Na_2SO_4$ and the solvent was evaporated *in vacuo*. The resulting residue was purified by column chromatography (silica gel, $CHCl_3$/MeOH, 20:1) to afford **MQ369$^{DART.2}$** (143.8 mg, 0.054 mmol, 61%); $^1$H NMR (700 MHz, $CD_3OD$) δ 8.04 (s, 1H), 7.77 (d, *J* = 7.9 Hz, 2H), 7.41 (d, *J* = 8.1 Hz, 2H), 7.06 (d, *J* = 7.5 Hz, 1H), 6.76 (s, 1H), 6.65 (d, *J* = 7.5 Hz, 1H), 4.64 (s, 2H), 4.59 (t, *J* = 5.1 Hz, 2H), 4.47 (s, 2H), 3.90 (t, *J* = 5.1 Hz, 2H), 3.84–3.75 (m, 7H), 3.72 (s, 1H), 3.69–3.55 (m, 138H), 3.55–3.44 (m, 10H), 3.42–3.34 (m, 3H), 2.84 (t, *J* = 7.1 Hz, 2H), 2.71 (t, *J* = 7.5 Hz, 2H), 2.62 (d, *J* = 9.0 Hz, 1H), 2.52 (t, *J* = 6.0 Hz, 2H), 2.11

(s, 4H), 2.06–1.99 (m, 3H), 1.86 (d, *J* = 14.3 Hz, 1H), 1.79–1.50 (m, 13H), 1.47–1.16 (m, 16H), 1.01–0.92 (m, 5H), 0.79–0.74 (m, 1H), 0.59 (s, 3H); HRMS (ESI-TOF) m/z: $[M + 3H]^{3+}$ Calcd for $C_{131}H_{235}ClN_5O_{46}$ 883.1958; Found 883.1944.

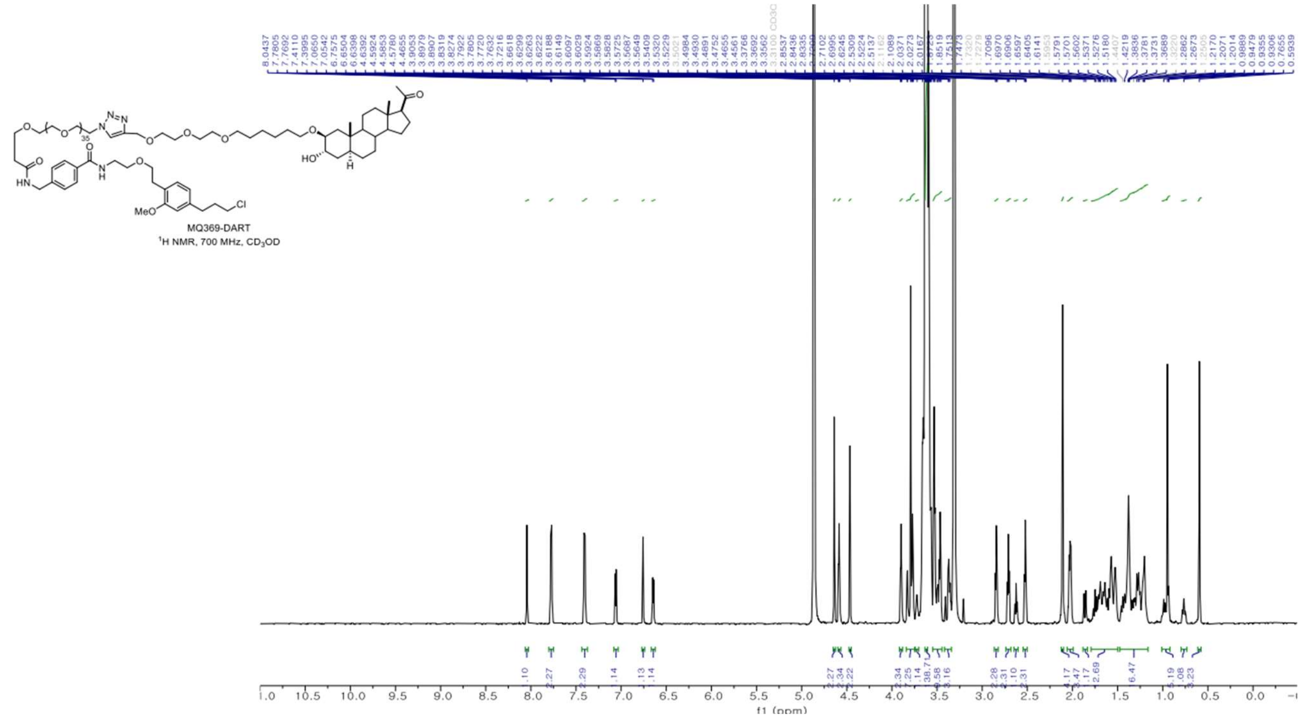

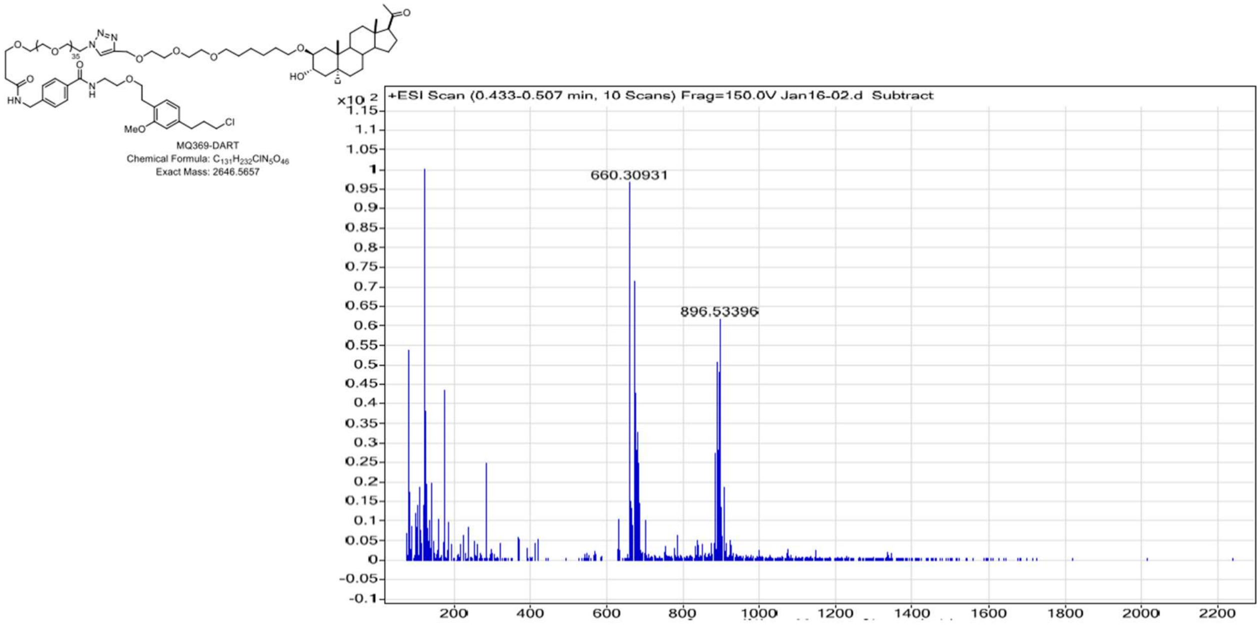


### (2.6) Preparation of MQ392$^{DART.2}$

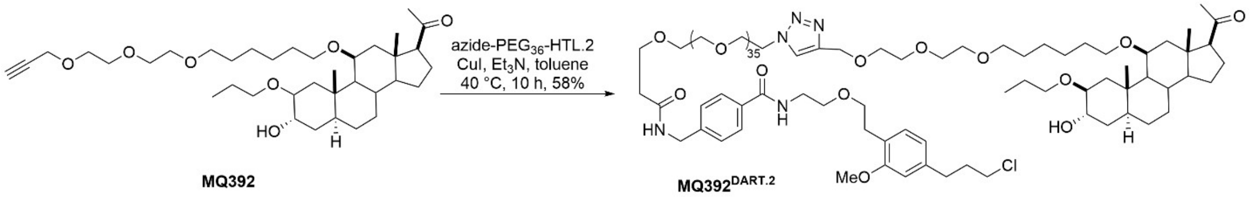


To a solution of **MQ392** (81 mg, 0.13 mmol) in toluene (6.5 mL) were added **azide-PEG$_{36}$-HTL.2** (271mg, 0.13 mmol), CuI (74 mg, 0.39 mmol) and $Et_3N$ (662 mg, 912 µL, 6.5 mmol) at 25 °C. The resulting reaction mixture was stirred at 40 °C for 10 h and concentrated *in vacuo*. The obtained residue was then dissolved in $CH_2Cl_2$ and washed with ice-cold $NH_4OH$ and $H_2O$. The organic layer was dried over $Na_2SO_4$ and the solvent was evaporated *in vacuo*. The resulting residue was purified by column chromatography (silica gel, $CHCl_3$/MeOH, 20:1) to afford **MQ392$^{DART.2}$** (204 mg, 0.075 mmol, 58%); $^1$H NMR (700 MHz, $CD_3OD$) δ 8.04 (s, 1H), 7.77 (d, *J* = 7.9 Hz, 2H), 7.40 (d, *J* = 7.9 Hz, 2H), 7.06 (d, *J* = 7.5 Hz, 1H), 6.76 (s, 1H), 6.64 (d, *J* = 7.5 Hz, 1H), 4.64 (s, 2H), 4.58 (t, *J* = 5.0 Hz, 2H), 4.46 (s, 2H), 3.90 (t, *J* = 5.2 Hz, 2H), 3.82 (d, *J* = 9.8 Hz, 2H), 3.79 (s, 3H), 3.77 (t, *J* = 6.0 Hz, 2H), 3.62 (d, *J* = 6.2 Hz, 155H), 3.45 (t, *J* = 6.7 Hz, 2H), 3.41 (s, 1H), 3.22–3.16 (m, 1H), 2.84 (t, *J* = 7.0 Hz, 2H), 2.71 (t, *J* = 7.5 Hz, 2H), 2.59–2.45

(m, 4H), 2.14–2.08 (m, 4H), 2.03 (p, *J* = 6.9 Hz, 2H), 1.92 (d, *J* = 14.1 Hz, 1H), 1.84–1.46 (m, 13H), 1.47–1.08 (m, 16H), 0.98 (dd, *J* = 12.5, 4.9 Hz, 1H), 0.93 (t, *J* = 7.4 Hz, 3H), 0.82 (dd, *J* = 10.7, 3.5 Hz, 1H), 0.76 (s, 3H); HRMS (ESI-TOF) m/z: $[M + 3H]^{3+}$ Calcd for $C_{134}H_{241}ClN_5O_{47}$ 902.5431; Found 902.5437.

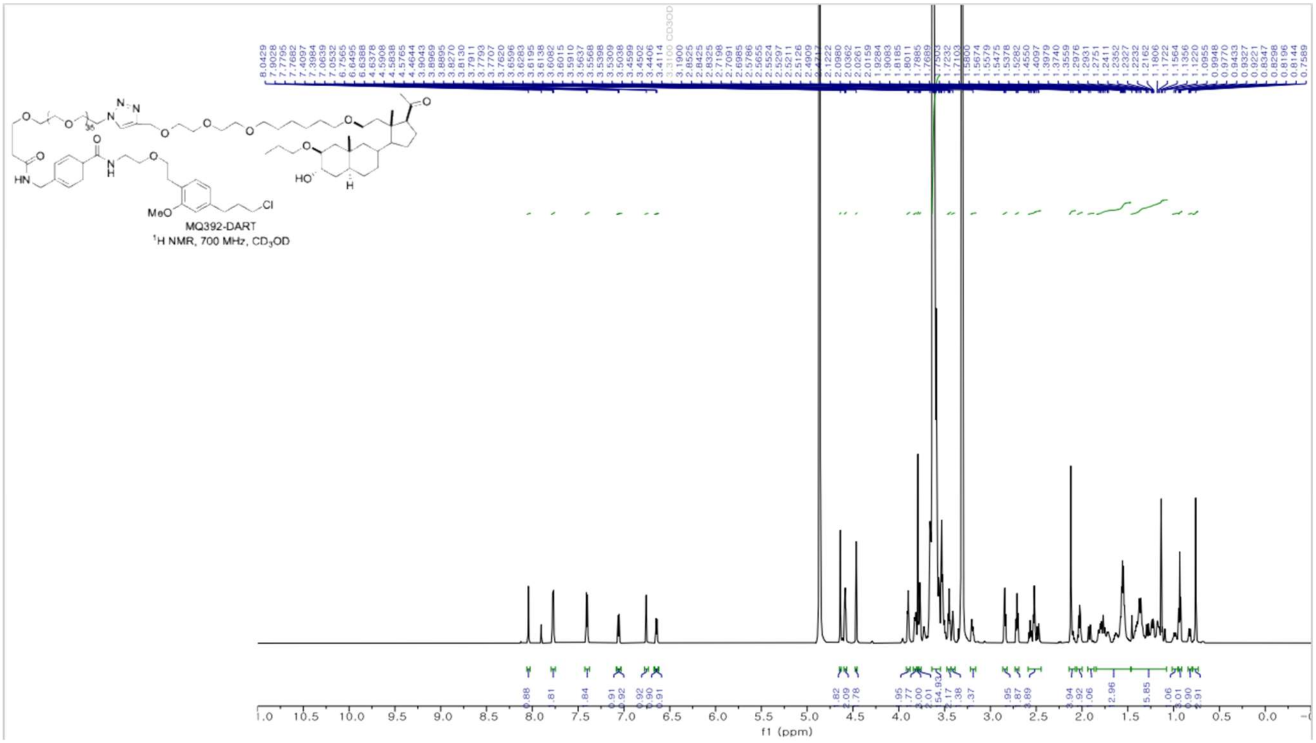

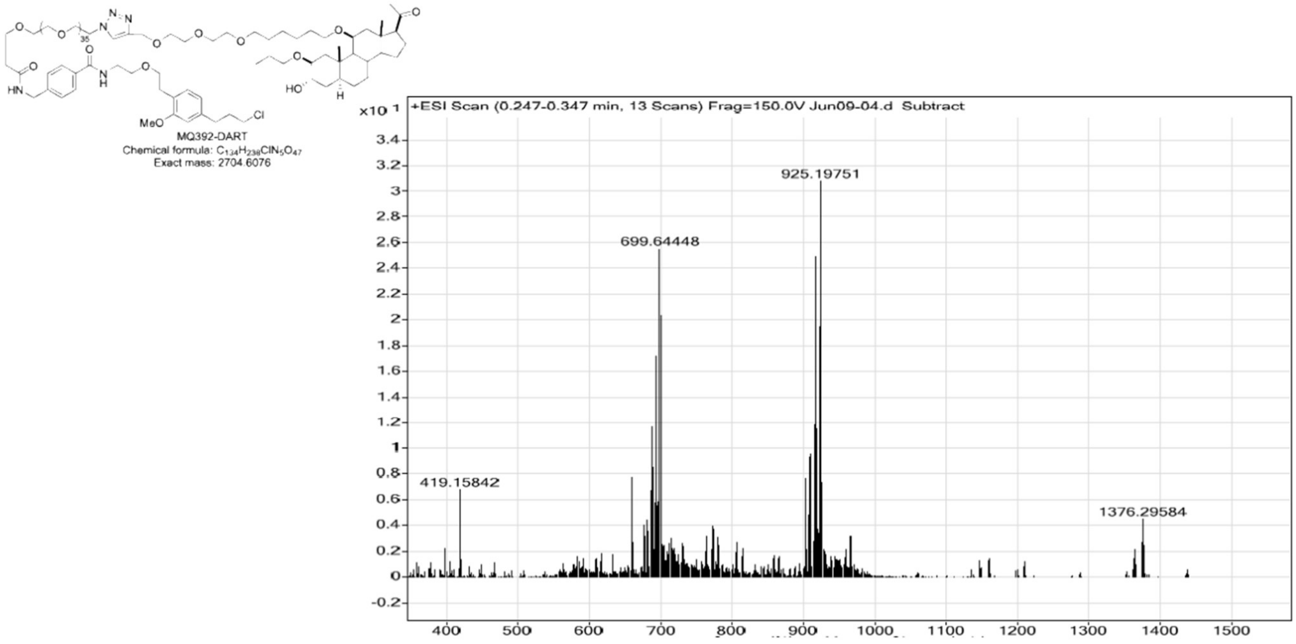


#### (2.7) Preparation of YX62[DART.2]

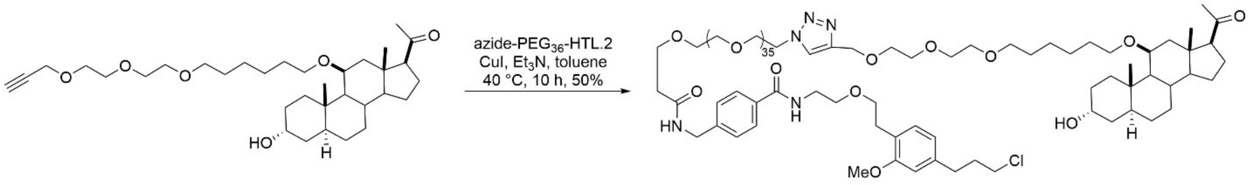


To a solution of **YX62** (62 mg, 0.11 mmol) in toluene (6.0 mL) were added **azide-PEG$_{36}$-HTL.2** (229mg, 0.11 mmol), CuI (62 mg, 0.33 mmol) and $Et_3N$ (560 mg, 722 µL, 5.5 mmol) at 25 °C. The resulting reaction mixture was stirred at 40 °C for 10 h and concentrated *in vacuo*. The obtained residue was then dissolved in $CH_2Cl_2$ and washed with ice-cold $NH_4OH$ and $H_2O$. The organic layer was dried over $Na_2SO_4$ and the solvent was evaporated *in vacuo*. The resulting residue was purified by column chromatography (silica gel, $CHCl_3$/MeOH, 20:1) to afford **YX62$^{DART.2}$** (145 mg, 0.055mmol, 50%); $^1$H NMR (700 MHz, $CD_3OD$) δ 8.05 (s, 1H), 7.78 (d, *J* = 8.0 Hz, 2H), 7.41 (d, *J* = 7.9 Hz, 2H), 7.06 (d, *J* = 7.5 Hz, 1H), 6.76 (s, 1H), 6.65 (d, *J* = 7.5 Hz, 1H), 4.64 (s, 2H), 4.59 (t, *J* = 5.1 Hz, 2H), 4.47 (s, 2H), 3.96 (d, *J* = 3.4 Hz, 1H), 3.90 (t, *J* = 5.2 Hz, 2H), 3.82 (d, *J* = 3.5 Hz, 1H), 3.81–3.75 (m, 5H), 3.65–3.52 (m, 160H), 3.45 (t, *J* = 6.8

Hz, 2H), 3.15 (q, $J$ = 6.9 Hz, 1H), 2.84 (t, $J$ = 7.1 Hz, 2H), 2.71 (t, $J$ = 7.4 Hz, 2H), 2.61–2.45 (m, 4H), 2.12 (s, 4H), 2.03 (p, $J$ = 6.9 Hz, 2H), 1.84–1.60 (m, 6H), 1.57–1.44 (m, 8H), 1.42–1.14 (m, 11H), 1.02 (s, 4H), 0.87 (dd, $J$ = 11.2, 3.4 Hz, 1H), 0.77 (s, 3H); HRMS (ESI-TOF) m/z: [M + 3H]$^{3+}$ Calcd for $C_{131}H_{235}ClN_5O_{46}$ 883.1958; Found 883.1945.

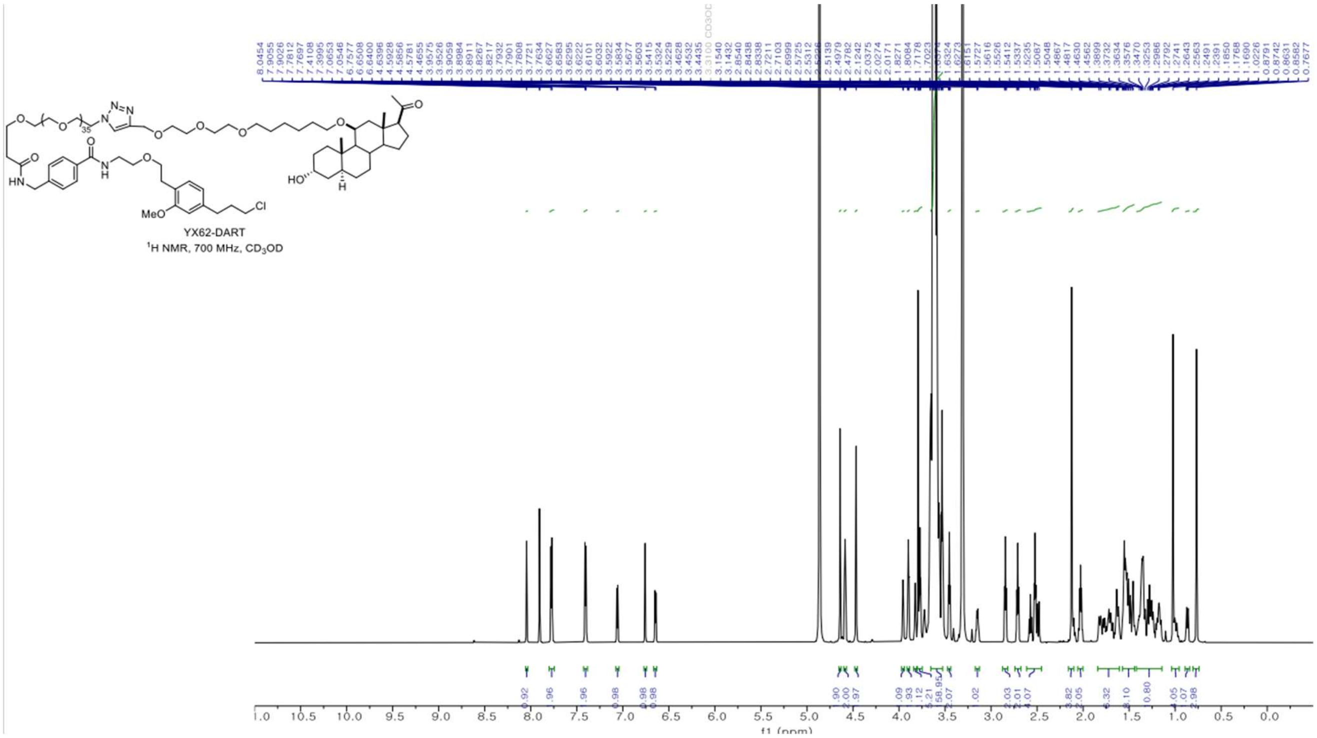

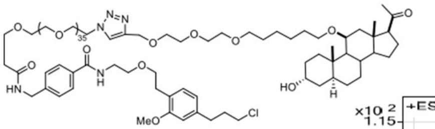

YX62-DART
Chemical Formula: $C_{131}H_{232}ClN_5O_{46}$
Exact Mass: 2646.5657

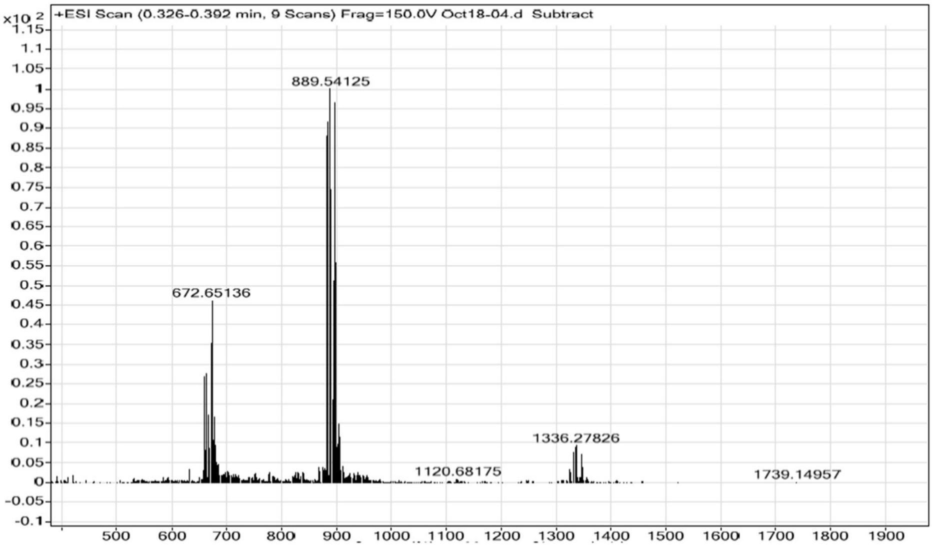


**(2.8) Preparation of YX85.1$^{DART.2}$**

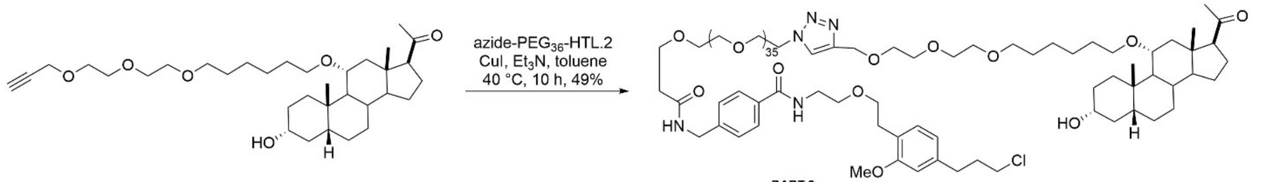


YX85 YX85.1$^{DART.2}$

To a solution of **YX85** (65 mg, 0.12 mmol) in toluene (6.0 mL) were added **azide-PEG$_{36}$-HTL.2** (250mg, 0.12 mmol), CuI (68 mg, 0.36 mmol) and $Et_3N$ (611 mg, 842 µL, 6.0 mmol) at 25 °C. The resulting reaction mixture was stirred at 40 °C for 10 h and concentrated *in vacuo*. The obtained residue was then dissolved in $CH_2Cl_2$ and washed with ice-cold $NH_4OH$ and $H_2O$. The organic layer was dried over $Na_2SO_4$ and the solvent was evaporated *in vacuo*. The resulting residue was purified by column chromatography (silica gel, $CHCl_3$/MeOH, 20:1) to afford **YX85.1$^{DART.2}$** (156 mg, 0.059mmol, 49%); $^1$H NMR (500 MHz, $CD_3OD$) δ 8.05 (s, 1H), 7.77 (d, *J* = 8.4 Hz, 2H), 7.41 (d, *J* = 8.2 Hz, 2H), 7.06 (d, *J* = 7.6 Hz, 1H), 6.76 (d, *J* = 1.6 Hz, 1H), 6.65 (dd, *J* = 7.6, 1.6 Hz, 1H), 4.64 (s, 2H), 4.61–4.57 (m, 2H), 4.47 (s, 2H), 3.92–3.89 (m, 2H), 3.80–3.75 (m, 6H), 3.65–3.60 (m, 154H), 3.47 (d, *J* = 6.6 Hz, 4H), 2.84 (t, *J* = 7.0 Hz, 2H), 2.74–2.64

(m, 3H), 2.53 (dt, $J$ = 11.8, 5.5 Hz, 3H), 2.33 (d, $J$ = 13.6 Hz, 1H), 2.14 (s, 3H), 2.07–1.99 (m, 2H), 1.94–1.86 (m, 1H), 1.79–1.43 (m, 16H), 1.40–1.17 (m, 12H), 1.04 (s, 3H), 1.01–0.91 (m, 1H), 0.59 (s, 3H); HRMS (ESI-TOF) m/z: $[M + 3H]^{3+}$ Calcd for $C_{131}H_{235}ClN_5O_{46}$ 883.1958; Found 833.1958.

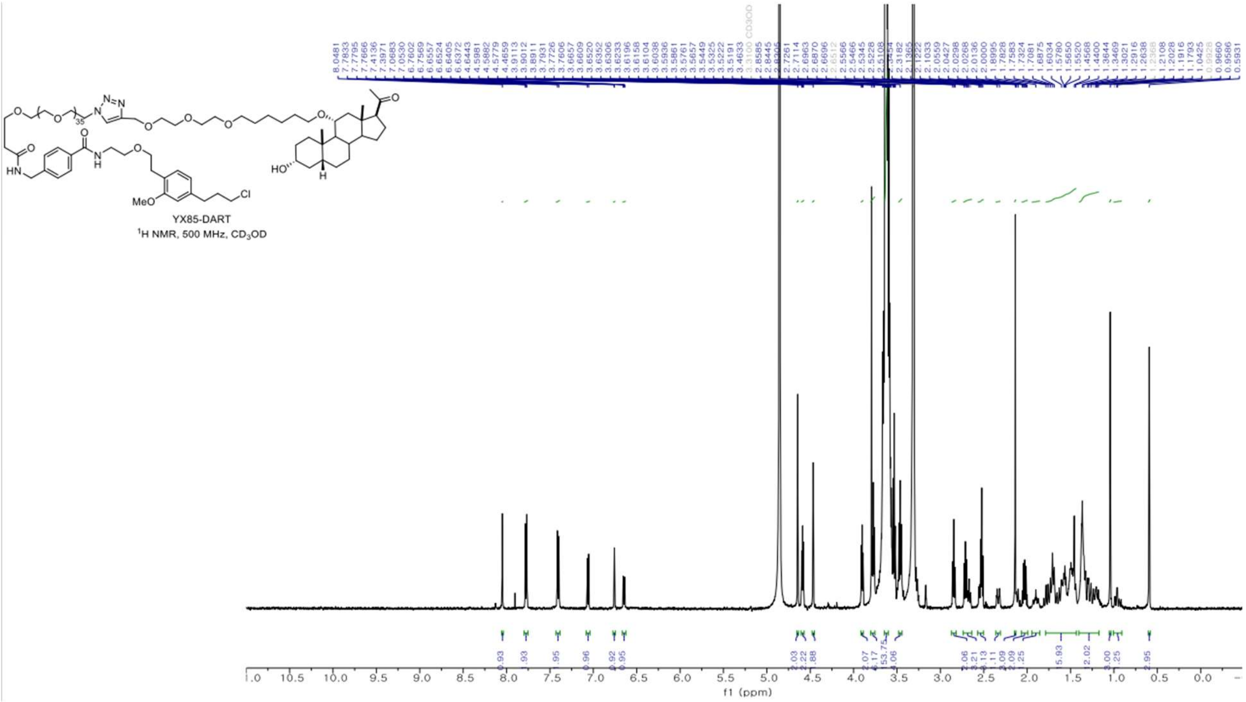

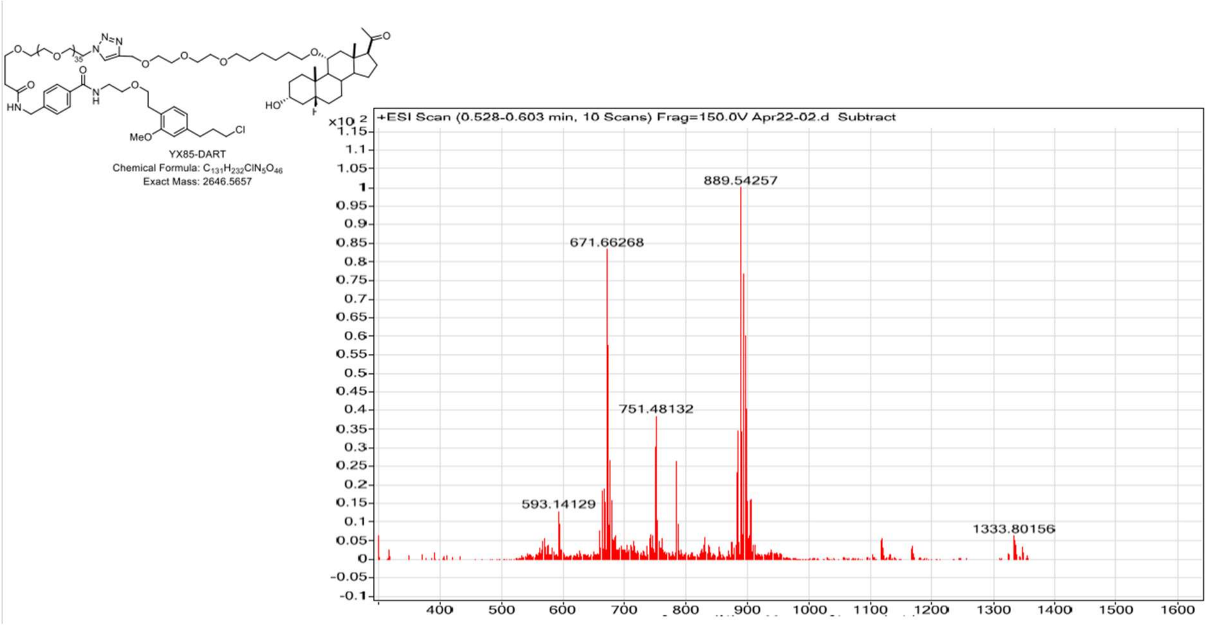


### 4. Reference


Shields, B.C., Yan, H., Lim, S.S.X., Burwell, S.C.V., Cammarata, C.M., Fleming, E.A., et al. (2024). DART.2: bidirectional synaptic pharmacology with thousandfold cellular specificity. Nat. Methods *21*: 1288–1297.
